# Low-ABC: a robust demographic inference from low-coverage whole-genome data through ABC

**DOI:** 10.1101/2024.08.01.606209

**Authors:** Maria Teresa Vizzari, Silvia Ghirotto, Rajiv Boscolo Agostini, Pierpaolo Maisano Delser, Lara Cassidy, Andrea Manica, Andrea Benazzo

## Abstract

The reconstruction of past demographic histories relies on the pattern of genetic variation shown by the sampled populations; this means that an accurate estimation of genotypes is crucial for a reliable inference of past processes. A commonly adopted approach to reconstruct complex demographic dynamics is the Approximate Bayesian Computation (ABC) framework. It exploits coalescent simulations to generate the expected level of variation under different evolutionary scenarios. Demographic inference is then performed by comparing the simulated data with the genotypes called in the sampled individuals. Low sequencing coverage drastically affects the ability to reliably call genotypes, thus making low-coverage data unsuitable for such powerful inferential approaches.

Here, we present Low-ABC, a new ABC approach to infer past population processes using low-coverage whole-genome data. Under this framework, both observed and simulated genetic variation are not directly compared using called genotypes, but rather obtained using genotype likelihoods to consider the uncertainty caused by the low sequencing coverage. We first evaluated the inferential power of this procedure in distinguishing among different demographic models and in inferring model parameters under different experimental conditions, including a wide spectrum of sequencing coverage (1x to 30x), number of individuals, number, and size of genetic loci.

We showed that the use of genotype likelihoods integrated into an ABC framework provides a reliable inference of past population dynamics, thus making possible the application of model-based inference also for low-coverage data. We then applied Low-ABC to shed light on the relationship between Mesolithic and Early Neolithic European populations.

## Introduction

A robust inference of past population dynamics is crucial to correctly reconstruct demographic and evolutionary trajectories. The constant increase in genome data availability has promoted the development of inferential algorithms based on a variety of statistical techniques, exploiting various aspects of genomic information (Marchi et al. 2021). Hidden Markov models methods (Li and Durbin 2011; Schiffels and Durbin 2014) use patterns of variation along the individuals’ genomes to estimate the age of the common ancestors of the sampled segments along the chromosomes; full-likelihood methods work assessing the probability of observing DNA sequences under relatively simple evolutionary models; site frequency spectrum (SFS) based methods (Gutenkunst et al. 2010; Jouganous et al. 2017; Terhorst et al. 2017; Excoffier et al. 2021) exploit the information provided by the distribution of the number of sites in a genome that have given allele frequencies in one or several populations to reconstruct past population histories; and Approximate Bayesian computations (ABC) methods (Bertorelle et al. 2010; Csilléry et al. 2010) which allow for complex scenarios for which the likelihood cannot be explicitly computed by replacing it with a distance measure between an observed and a simulated set of summary statistics, chosen to be informative about the model parameters.

Regardless of the method used to perform the inference, all these methods assume that the individual genotypes are known. Whilst this assumption is not problematic for high coverage data, the level of uncertainty associated with the genotype calling increases with decreasing sequencing coverage levels (Nielsen et al. 2012). Low-coverage data are common for non-model species, due to budget limitations, and in the study of ancient DNA, where the limited amount of endogenous content makes high coverage genomes unfeasible for many samples. The lack of methods for demographic inference able to cope with the loss of information related to low-coverage data greatly limits the kind of questions that we can ask under these scenarios (Lou et al. 2021).

A number of approaches designed to describe population differences based on low-coverage data have been developed by taking a probabilistic approach in which polymorphic sites and genotypes are not directly characterised from the data, but rather estimated from the so-called genotype likelihoods (GLs, Nielsen et al. 2011). The GLs are generally computed from the aligned reads to a reference genome and express the probability of the observed read configuration given a certain genotype, at a particular genomic site, for a particular individual, while also integrating the error rate of the sequencing machine. It is also possible, using a Bayesian framework, to incorporate prior information to the inference (e.g. the population allele frequencies) in order to produce posterior probabilities for the genotypes (Nielsen et al. 2011). Under these approaches, the genotype calling phase is avoided and the GLs have been directly used to estimate the SFS (Korneliussen et al. 2014), to characterize population’s structure (Meisner and Albrechtsen 2018), to test for introgression (Soraggi et al. 2018) and to reliably use ancient DNA by explicitly incorporating post-mortem DNA damage in the estimation of GLs (Kousathanas et al. 2017; Lou et al. 2021).

Whilst these methods have been shown to produce a higher number of true positive polymorphic sites compared to direct calling of low coverage data, thus reducing the bias in downstream population genetic analyses (Han et al. 2014; Korneliussen et al. 2014; Kousathanas et al. 2017), GLs have yet to be incorporated within a model-based inferential framework. To date, there are no inference of complex demographic models (i.e. including population sizes and split times, as well as migration) that leverage low-coverage genomes.

In this paper we developed an ABC-based procedure (called Low-ABC) which enables the fitting of complex demographic models to low coverage data, explicitly accounting for the amount of data available for each sample. The idea behind this framework is to integrate the calculation of genotype likelihoods for a specific sample sequencing coverage level and error rate in the simulation step, to perform a fair comparison with the observed data. In the implementation we present, we then estimated directly from the genotype likelihood of observed and simulated data the FDSS, a statistic that has proven to be effective in the reconstruction of past population dynamics (Ghirotto et al. 2021), to perform an ABC-Random Forest (ABC-RF) model comparison and parameters estimation. Integrating the GLs within simulations, we can explicitly account for differences in the coverage level among samples and to also consider the effect of platform-specific sequencing error rates. We assessed the effectiveness of this conceptual framework in selecting the true demographic history using pseudo-observed datasets, under different sampling strategies and accounting for decreasing levels of sequencing coverage. We evaluated the power of the procedure considering a mean coverage of 30x (thus mimicking the robustness of the estimates achievable using high coverage data), 5x, 2x and 1x, for different sets of evolutionary scenarios. We show that Low-ABC had a high success rate at selecting the correct model and recovering the underlying demographic parameters, and that it is superior to comparing coalescent simulations directly to GL-based summary statistics of the observed low-coverage.

Finally, we applied Low-ABC to the analysis of alternative hypotheses depicting the relationships among Neolithic farmers and Mesolithic hunter-gatherer populations during the late Pleistocene and early Holocene in Eurasia (Jones et al. 2015; Lazaridis et al. 2016a; Marchi et al. 2022; Stoneking et al. 2023).

## Materials and Methods

Approximate Bayesian Computation (ABC) is a simulation-based inferential framework developed to quantitatively compare alternative demographic models and to infer their parameters. Under this Bayesian approach, the likelihood function is approximated by simulation, incorporating prior information about the parameters that define the population dynamics under investigation. The whole ABC procedure is based on the comparison between simulated and observed genomic data, summarized by the same set of summary statistics selected to be informative about the evolutionary processes of interests (Bertorelle et al., 2010; Csilléry et al., 2010). The classical ABC procedure requires the simulation of millions of data sets of the same size as those observed, thus becoming computationally expensive as increasing the size of the datasets and the complexity of the models tested. A recent integration of ABC, the ABC-Random Forest (ABC-RF), based on a machine learning algorithm, allowed to overcome the above-cited limitations, and facilitated the application of ABC to investigate complex demographic models through the analysis of complete genomes. ABC-RF requires fewer simulations per model, and it accommodates large dimensional summary statistics with no consequences on the estimation performances (Pudlo et al. 2016; Raynal et al. 2019; Ghirotto et al. 2021).

We summarized the genetic data through the FDSS, i.e. the frequency distributions of genomic loci carrying the four mutually exclusive categories of segregating sites for pair of populations (i.e., private polymorphisms in either population, shared polymorphisms, and fixed differences (Wakeley and Hey 1997). To calculate the FDSS we considered the genome as subdivided into *k* independent fragments of length *m*, and for each fragment, we counted the number of segregating sites belonging to each of the categories defined above. Each distribution describes the observed frequency of *k* genomic loci having from 0 (monomorphic loci) to *n* segregating sites in each category, for each pair of populations considered (see Ghirotto et al 2021 for details).

Under the Low-ABC approach presented here, the FDSS is not directly calculated from known genotypes, but rather estimated using genotype likelihoods, as detailed in the next section.

### Calculate the FDSS from Genotype Likelihoods

We calculated the FDSS based on GLs from the output of the *ms* coalescent simulator (Hudson, 2002), following the steps detailed below:

1. Simulate a certain number of independent loci of length *w* base pairs, using the *ms* coalescent simulator.
2. For each locus, for each one of the *w* sites (both polymorphic and monomorphic), we sample the number of simulated reads covering that site for each diploid individual according to a Poisson distribution with a mean equal to a user-defined mean coverage. A specific mean coverage for each simulated individual could be optionally set.
3. Sampled “reads” are then assigned to the two chromosomes (assuming diploidy) of each individual accordingly to a binomial distribution with a probability of success of 0.5.

By sampling the number of available reads from a Poisson distribution given a mean coverage, we take into account the possibility that some nucleotide positions may be not covered, thus making our method able to deal with the presence of missing data (typically observed in low-coverage genomes).

4. At this point, a Phred quality score is assigned to each sampled read. The Phred quality score is a measure of the quality of the identification of the nucleobases generated by DNA sequencing, and it represents the chances that a base is incorrectly called and is defined as *Q* = − 10*log*_10_ *P*, were *P* represent the base-calling error probability. In the simulations, we assumed that every base has the same quality score of 30. A Phred quality score Q30 means that the probability of an incorrect base call is 1 in 1,000.
5. A certain amount of errors (wrong nucleotides) are introduced according to a binomial distribution *B*(*n,p*), where *n* is the number of reads covering each site and *p* is the NGS error rate (that we fixed to 1% as observed for the Illumina platform).
6. Given the error rate, the Phred quality scores and the list of nucleotides (sampled “reads” *D*), the genotype likelihoods, *P*(*D|G*), for all the 10 possible diploid genotypes are finally calculated as follows:

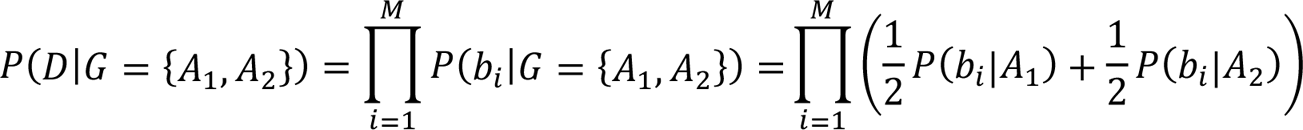

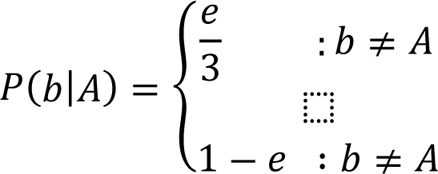

where *M* is the sequencing coverage, *_bi_* is the observed base in read *i* and *e* is the probability of error calculated from the Phred quality score (Van der Auwera et al. 2013).

7. The conditional posterior probability of the genotype *G* given the observed data *D*, formally *P*(*G|D*), is finally computed according to the Bayes’ Theorem:

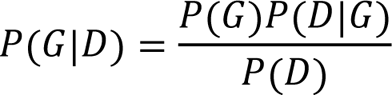

where *D* is the list of nucleotides observed at each site. This probability depends on the prior probability of the genotype, *P*(*G*), and on the conditional probability of the data given the genotype, *P*(*D|G*), computed in 6.

The prior probability of a genotype, *P*(*G*), represents out expectation to see a certain genotype in the population according to the evolutionary model. In this framework the prior of the genotype frequency for each population is computed using the allele frequency counts from reads under the assumption of Hardy-Weinberg Equilibrium.

8. In the case of a single population, the genotype posterior probabilities across all individuals are used to estimate the probability of the site to be segregating in the population, *P*(*POP_xpoly_*), as follow:

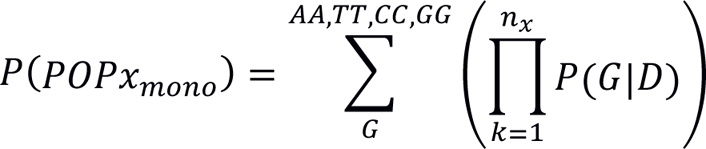

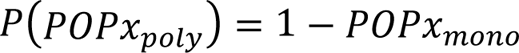

where *P(G|D)* represents the posterior probabilities of the homozygous genotypes and *n_x_* indicates the number of individuals sampled from the population *x*.

Single-site probabilities were combined to determine the expected number of segregating sites, _E_[*ns*], in a single locus (of length *w*) and the FDSS was finally computed across all the simulated loci.

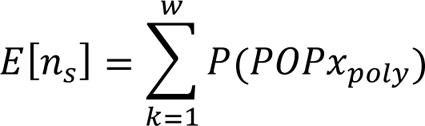

In the case of two or more populations, we computed for all the possible pairs of populations the probability of a site to be:

a) Monomorphic in *Pop1* and *Pop2*:

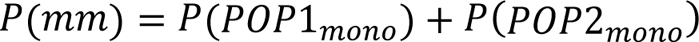

b) Segregating in *Pop1* but monomorphic in *Pop2*:

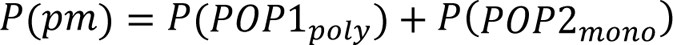

c) Monomorphic in *Pop1* but segregating in *Pop2*:

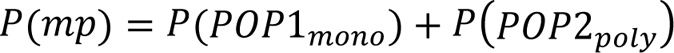

d) Segregating in *Pop1* and *Pop2*:

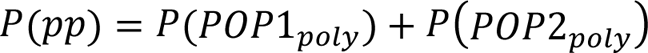

e) Fixed for different alleles in *Pop1* and *Pop2*:

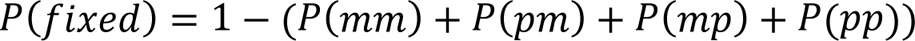

The FDSS was then computed across loci for each segregating-sites category as previously described.

The approach described above works by position, generating ten genotype posterior probabilities at both polymorphic and monomorphic sites, thus it may become computationally expensive and time consuming when analysing thousands of independent loci of several Kb in length for each coalescent simulation. To partially overcome this issue, we decided to approximate the estimation of the GLs for the monomorphic fraction of each simulated locus, which is the most demanding phase of the framework. Since the genotype posterior probabilities for a monomorphic site only depend on the error rate and the mean coverage, we repeated steps 2-7 in a large sample of monomorphic sites (10,000 sites). Then, we used the formulas in 8 to compute the expected probability for a monomorphic site to be correctly identified as monomorphic (a) or polymorphic in every segregating site category (b, c, d, e) due to the error rate and mean coverage. These expected probabilities were generated once and then used to model the monomorphic part of each locus in all the simulated dataset generated under a demographic model.

### Power Analysis

To evaluate the robustness of our procedure, we conducted an extensive simulation study conducting a power analysis at different coverage levels. We also explored the inferential power of Low-ABC with respect to different experimental conditions, evaluating the consequences of sampling strategies involving different numbers of chromosomes, different numbers of loci, and different locus lengths. We tested all the possible combinations of locus length (bp) {200; 1,000}, number of independent genomic loci {1,000; 5,000}, number of chromosomes sampled per population {10, 20, 50} and four different coverage levels {1x, 2x, 5x, 30x} for a total of 48 combinations of sampling strategies tested. For each combination, we generated 100,000 simulated datasets with a fixed intra-locus recombination rate (1x10-8/bp/generation), and with a fixed mutation rate (1x10-8 /bp/generation). We evaluated the power considering two sets of demographic models (detailed below). The FDSS were estimated from the genotype likelihood calculated from the *ms* (Hudson 2002) output of each simulation. For each combination of experimental conditions, we compared alternative models within each set of demographic scenarios evaluating the associated classification error; to evaluate the power of our framework in estimating demographic parameters, we generated 1,000 pods for each model and combination of experimental condition tested. All the ABC-RF estimates have been obtained using the functions *abcrf*, for model selection, and *regAbcrf*, for parameters estimation, and employing forests of 2,000 trees; both functions are integrated in the R-package *abcrf* (Pudlo et al. 2016; Raynal et al. 2019). To assess the power of the model selection procedure, we evaluated the out-of-bag Classification Error (CE), representing the proportion of simulated datasets not assigned to the models that actually produced them, and the proportion of true positives (1-CE), representing the proportion of correctly assigned simulated datasets to the true model. To determine the power of our procedure in correctly estimating demographic parameters we calculated: 1) the coefficient of determination (R^2^), that represents the proportion of variance of the parameters explained by the summary statistics used in the regression model construction by the random forest algorithm. Although there is no universally defined threshold, we considered parameters with an R^2^ below 10% unreliably estimated, since the summary statistics do not adequately capture the information necessary for parameter estimation (Neuenschwander et al. 2008); 2) the relative bias, calculated by summing the differences between the 1,000 estimates of each demographic parameter and the true known (true) value, as:

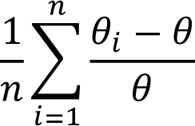

where *θ_i_* is the estimator of the parameter *θ* (true value), and *n* is the number of pods used (1,000 in our case). Because bias is relative, a value of 1 corresponds to a bias equal to 100% of the true value; 3) the root mean square error (RMSE), calculated by summing the squared differences between the 1,000 parameter estimates and the corresponding true value as:

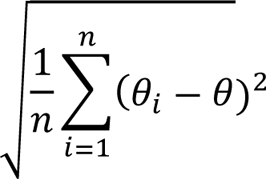

4) the factor *2*, representing the proportion of the 1,000 estimated median values lying between 50% and 200% of the true value, and 5) the 50% coverage, defined as the proportion of times that the known value lies within the 50% credible interval of the 1,000 estimates.

#### Tested models: One-population models

We first analyzed a set of models involving a single population evolving under three different scenarios (Figure 1). The first model (*Constant*), representing a population with a constant effective population size through time (*N1*). The second model (*Bottleneck*), describing a population that *T* generations ago has undergone an instantaneous bottleneck event, with the effective population size decreasing from *NaBott* to *N1Bott*. The third model (*Exponential Growth*), representing an exponentially growing population. The expansion started *T* generations ago, with the effective population size increasing from *NaExp* to *N1Exp*. The demographic parameters associated to each demographic model are drawn from uniform prior distributions (Table 1).

**Figure 1.**
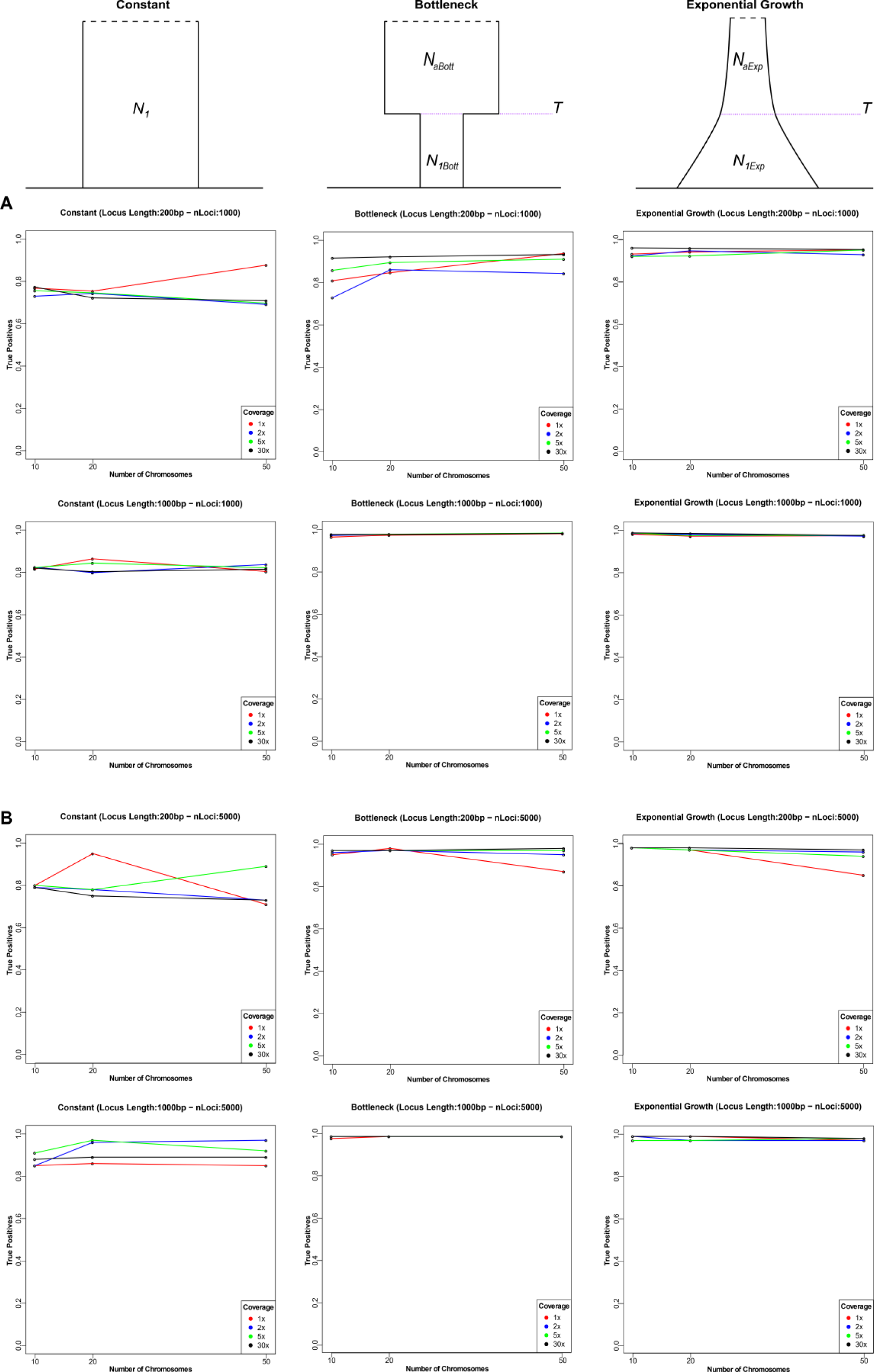
Proportion of True Positives for the one-population models. The plot below represents the proportion of TPs obtained analyzing pods coming from the three models under 48 combinations of experimental parameters. A) Combinations considering 1,000 loci. B) Combinations considering 5,000 loci. Number of chromosomes is in the x-axis, coverage levels are represented by different colors.

**Table 1.** Demographic parameters and prior distributions of One-Population models. Mutation and Recombination rates are expressed per nucleotide per generation.

| Demographic Parameters | Prior Distributions |
| --- | --- |
| Effective population size ($N_I$) | Uniform {500:50,000} |
| Effective population size ($Na_{Bott}$) | Uniform {25,000:100,000} |
| Effective population size ($NI_{Bott}$) | Uniform {500:5,000} |
| Effective population size ($Na_{Exp}$) | Uniform {500:5,000} |
| Effective population size ($NI_{Exp}$) | Uniform {25,000:100,000} |
| Time bottleneck ($T$) | Uniform {100:20,000} |
| Time exponential growth ($T$) | Uniform {100:20,000} |
| Mutation rate | $1 \times 10^{-8}$ {Fixed} |
| Recombination rate | $1 \times 10^{-8}$ {Fixed} |

#### Tested models: Two-populations models

We then moved to considering three demographic models with two populations (Figure 2). The first model (*Divergence*) describing an ancestral population of size *Nanc* that split *Tsep* generations ago into two different populations. These two derived populations evolve with a constant population size (*N1* and *N2*) until present time. The second model (*Divergence with Migration*) also included a continuous and bidirectional migration event between the two derived populations, from the divergence to the present. Under the third model (*Divergence with Admixture*), a single pulse of bidirectional admixture occurred at time *Tadm* after the divergence. Admixture rates (*adm12, adm21*), and event’ times are drawn from uniform priors; migration rates (*m12, m21*) are drawn from an exponential prior with mean 0.1 (Table 2).

**Figure 2.**
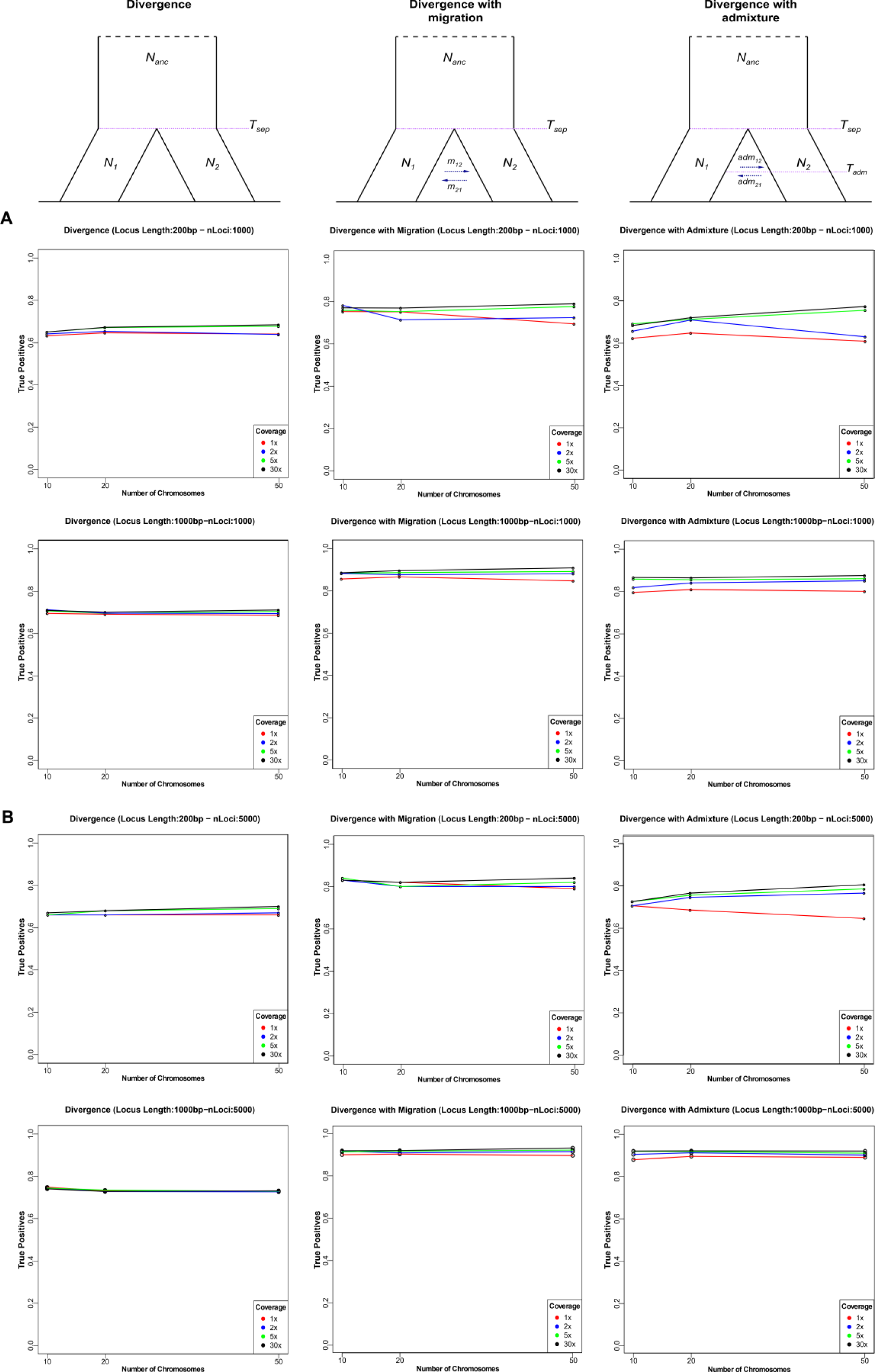
Proportion of True Positives for the two-populations models. A) Combinations considering 1,000 loci. B) Combinations considering 5,000 loci. The plots have the same features of Figure 1.

**Table 2.** Demographic parameters and prior distributions of Two-Population models. Mutation and Recombination rates are expressed per nucleotide per generation.

| Demographic Parameters | Prior Distributions |
| --- | --- |
| Effective population size ($N_{anc}$, $N_I$, $N_2$) | Uniform {500:50,000} |
| Time split ($T_{sep}$) | Uniform {300:20,000} |
| Migration rate ($m_{I2}$, $m_{2I}$) | Exponential {0.1} |
| Time admixture ($T_{adm}$) | Uniform {50:2,500} |
| Admixture rate ($adm_{I2}$, $adm_{2I}$) | Uniform {0.05:0.20} |
| Mutation rate | $1 \times 10^{-8}$ {Fixed} |
| Recombination rate | $1 \times 10^{-8}$ {Fixed} |

#### Comparison with using exact simulations and GL-based summary statistics from ANGSD

We finally compared the robustness and performance of Low-ABC with simpler approach of comparing the coalescent simulations directly to GL-based summary statistics of the low coverage data. In this approach, there is mismatch between simulated and observed data (the former are without error, whilst the latter attempt to correct for the uncertainty via GL). We produced 1,000 pseudo-observed datasets, for each combination of experimental condition tested, we employed the *msToGlf* tool implemented in ANGSD (Korneliussen et al. 2014) to compute the simulated genotype likelihoods directly from the *ms* output. From each pod we simulate GLs according to the different coverage levels tested (1x, 2x, 5x and 30x) and a fixed sequencing error of 1%. We then estimated the FDSS from the simulated GLs and compared these pseudo-observed FDSS from genotype likelihood with the FDSS directly calculated from *ms* simulations. To assess the power of this procedure, we evaluated the out-of-bag Classification Error (CE), the proportion of true positives (1-CE).

### Real Case: Mesolithic hunter-gatherer and Early Farmers relationships

We applied Low-ABC procedure to investigate the evolutionary relationship between Eurasian Mesolithic hunter-gatherer and Early Farmers. We considered ancient complete genomes already available for these groups (Gamba et al. 2014; Lazaridis et al. 2014; Olalde et al. 2014; Jones et al. 2015; Hofmanová et al. 2016; Kılınç et al. 2016; González-Fortes et al. 2017; de Barros Damgaard et al. 2018; Günther et al. 2018; Valdiosera et al. 2018; Cassidy et al. 2020; Maisano Delser et al. 2021; Saag et al. 2021), detailed in Table 3. All the collected samples were analyzed following the bioinformatic pipeline detailed in Maisano Delser et al. (2021).

**Table 3.**
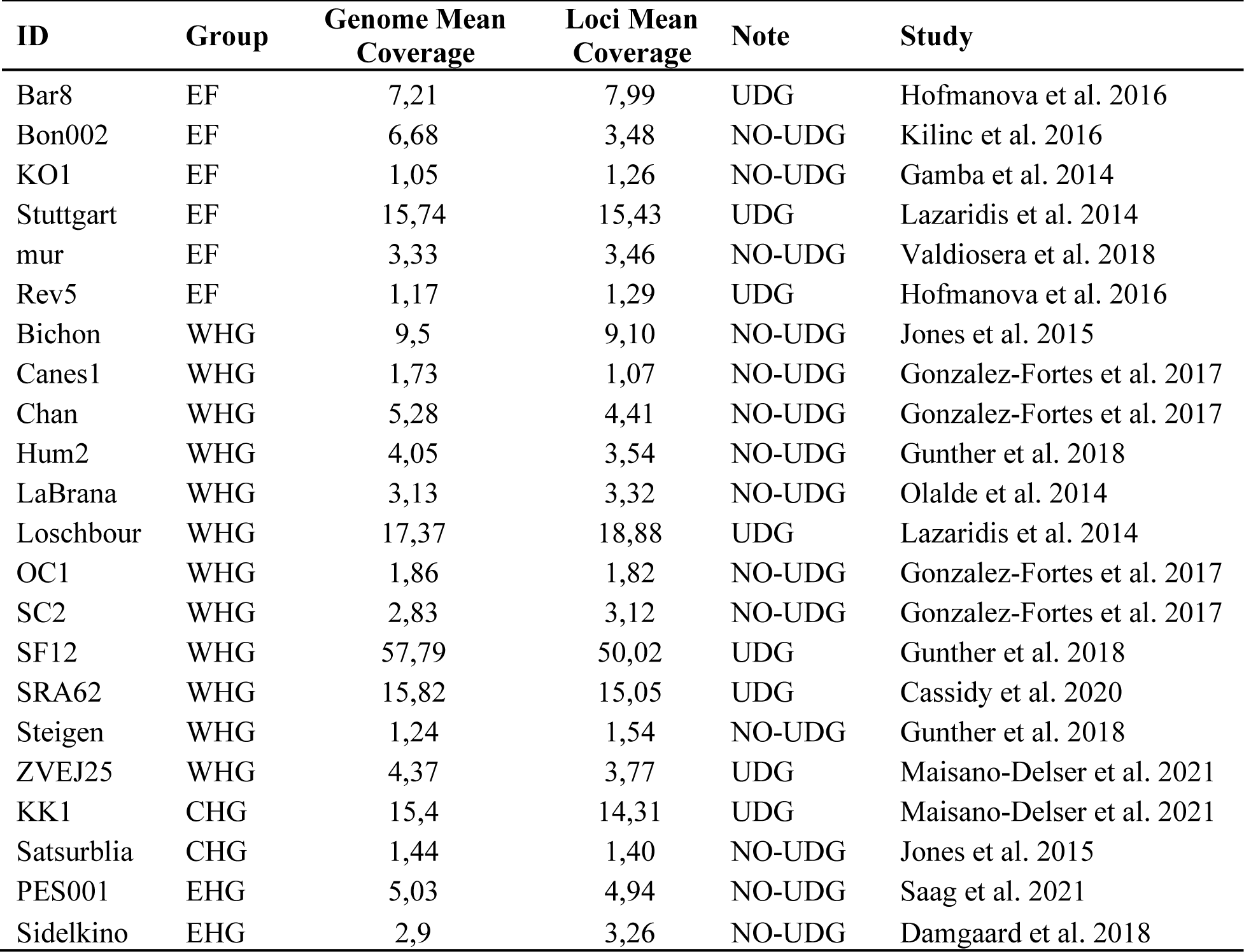
Ancient individuals analyzed.

Adapters were trimmed with cutadapt v1.9.1 (Martin 2011) and then raw reads were aligned to human reference sequence hg19/GRCh37 using bwa aln v0.7.12 (Li and Durbin 2010) with seeding disabled (-l 1000), maximum edit distance set to -n 0.01 and maximum number of gap opens set to -o 2. Aligned reads were filtered for minimum mapping quality 20 with samtools view v1.9 (Li et al. 2009). Indexing, sorting and duplicate removal (rmdup) were performed with samtools v1.9. Indels were realigned using The Genome Analysis Toolkit v3.7 (GATK, (Van der Auwera et al. 2013), employing the modules RealignerTargetCreator and IndelRealigner, and 2 bp at the start and ends of reads were softclipped using a custom python script (https://github.com/EvolEcolGroup/data_paper_genetic_pipeline).

From the aligned Bam files, we generated a bed file that describes the regions that were callable for each sample using the CallableLoci tool from GATK (version 3.7.0); we considered genomic regions with a minBaseQuality score of 30, a minMappingQuality score of 30, and a minDepth of 1. Next, we intersected the callable loci regions (i.e. regions marked with CALLABLE flag) of all the samples with bedtools multiIntersect, v2.26.0, (Quinlan and Hall 2010). To calculate the observed *FDSS* we only considered autosomal regions outside known and predicted genes ± 10,000 bp and outside CpG islands and repeated regions (as defined by the UCSC tracks, (Hinrichs et al. 2016). We extracted 12,507 independent fragments of 1,000 bp length, covered in at least the 80% of the samples (18 out of 22 individuals). We then generated genotype likelihoods at the selected loci with ANGSD setting a minimum mapping quality of 30 (-minMapQ 30), a minimum base quality of 30 (-minQ 30), the gatk genotype likelihood model (-GL 2) and emitted the genotype likelihoods for all the 10 possible genotypes at each position (-doGlf 4). The minor allele frequencies of the populations under investigation were estimated with ANGSD based on the allele counts from reads (-doCounts 1 -doMajorMinor 2 -doMaf 8) setting a minimum mapping quality of 30 (-minMapQ 30) and a minimum base quality of 30 (-minQ 30). Finally, we calculate the observed FDSS from GLs.

To investigate human past population dynamics during the early Holocene, we designed two alternative scenarios (Figure 3) that summarize the key events that characterized the relationships among the four major Eurasia groups at that time: Western Hunter-Gatherers (WHG), Eastern Hunter-Gatherers (EHG), Caucasus Hunter-Gatherers (CHG) and Anatolian and European Early Farmers (EF).

**Figure 3.**
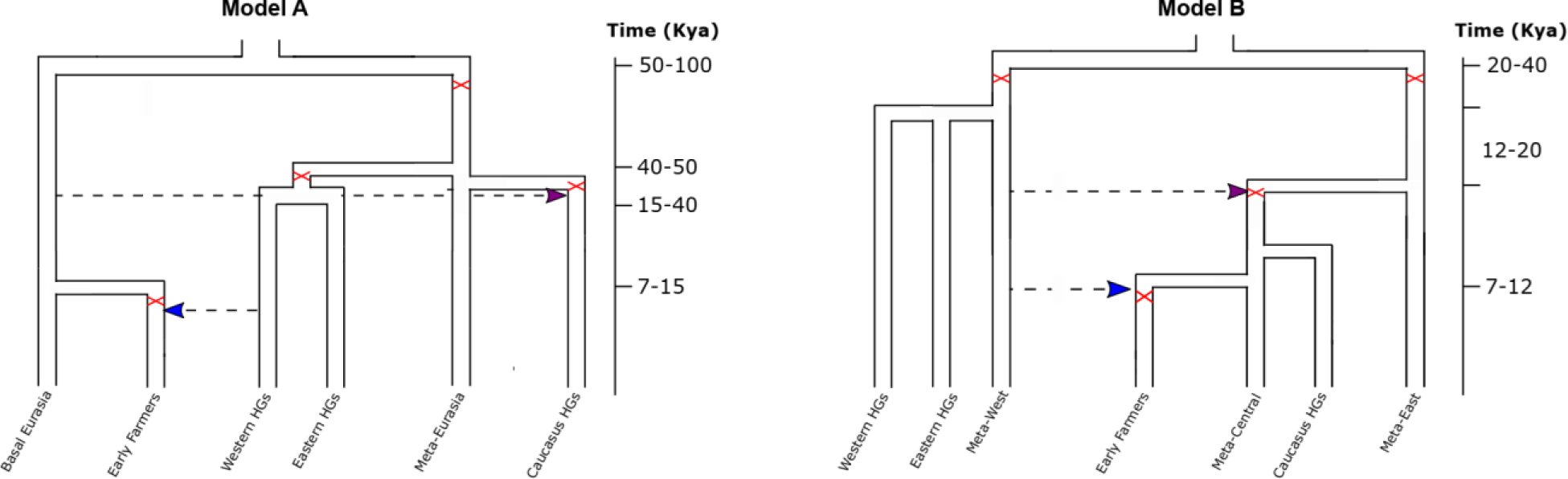
Demographic models defining alternative historical relationships between Early Farmers and Mesolithic Hunter-Gatherers.

The first model (Model A) summarizes the evidence that emerged from Jones et al (2015), Lazaridis et al. (2016) and Wang et al. (2019). We defined two ancestral ghost populations that existed before the differentiation of all other Eurasian lineages. The first population, called Meta-Eurasia population, gave rise to the three main groups of hunter-gatherers shortly after anatomically modern humans expanded from Africa into Europe (∼60-50 kya). The second population, known as Basal Eurasia, was the source of the first Anatolian/European Farmers. We account for the genetic contribution of Western HGs in the genomic ancestry of EFs by modelling an admixture event that took place during their differentiation and modelled with another admixture event the contribution of the Basal Eurasia in the Caucasus HGs gene pool.

To design the second model (Model B), we followed the parametrization recently proposed by Marchi et al. (2022) with minor modifications detailed below. Unlike Marchi et al (2022), we modelled the samples by considering them within populations and not as single genomes. We modelled three metapopulations that represent the ancestral genetic pools of our ancient individuals: the Meta-West related to the Western and Eastern HGs populations, the Meta-Central from which both Caucasus HGs and Anatolian/European EFs descend, and finally the Meta-East from which the Meta-Central population differentiated. Under this model the divergence between the Western and Eastern metapopulations is dated back to the Last Glacial Maximum (LGM ∼25 kya). We also account for two different pulses of admixture from the Meta-West population in both Meta-Central and in the Early Farmer populations.

The prior distributions associated with the demographic parameters that define these models were set following what was proposed in the recent literature (Jones et al. 2015; Wang et al. 2019; Marchi et al. 2022), and are reported in Tables 4 and 5. We considered a generation time of 29 years, and we fixed the mutation rate at 1.25×10^-8^ bp/generation and the intra-locus recombination rate at 1.12×10^-8^ (Scally and Durbin 2012).

**Table 4.** Demographic parameters and prior distributions of Model A. Mutation and Recombination rates are expressed per nucleotide per generation, Times are expressed in years.

| Parameters | Prior | Min | Max | Parameter | Value (Fixed) |
| --- | --- | --- | --- | --- | --- |
| Ne (Sampled pops) | logunif | 500 | 50,000 | tgen | 29 |
| Ne (Meta pops) | unif | 1,000 | 100,000 | mu | 1.25e-8 |
| Intensity (Bottleneck) | unif | 2 | 100 | recomb | 1.12e-8 |
|  |  |  |  | tsampEF | 6,500 |
| admxEFWHG | unif | 10% | 99% | tsampWHG | 7,500 |
| admXCHGBE | unif | 10% | 99% | tsampCHG | 9,000 |
|  |  |  |  | tsampEHG | 10,000 |
| tsepMETAUEBE | unif | 50,000 | 100,000 | tBott | 29/ 1gen |
| tsepCHGMETAUE | unif | 40,000 | tbottMETAUEBE |  |  |
| tsepHGMETAUE | unif | 15,000 | tbottMETAUEBE |  |  |
| tsepEFBE | unif | 7,000 | 15,000 |  |  |
| tadmXCHG | equal to | tsepCHGMETAUE |  |  |  |
| tadmXEF | equal to | tsepEFBE |  |  |  |

**Table 5.**
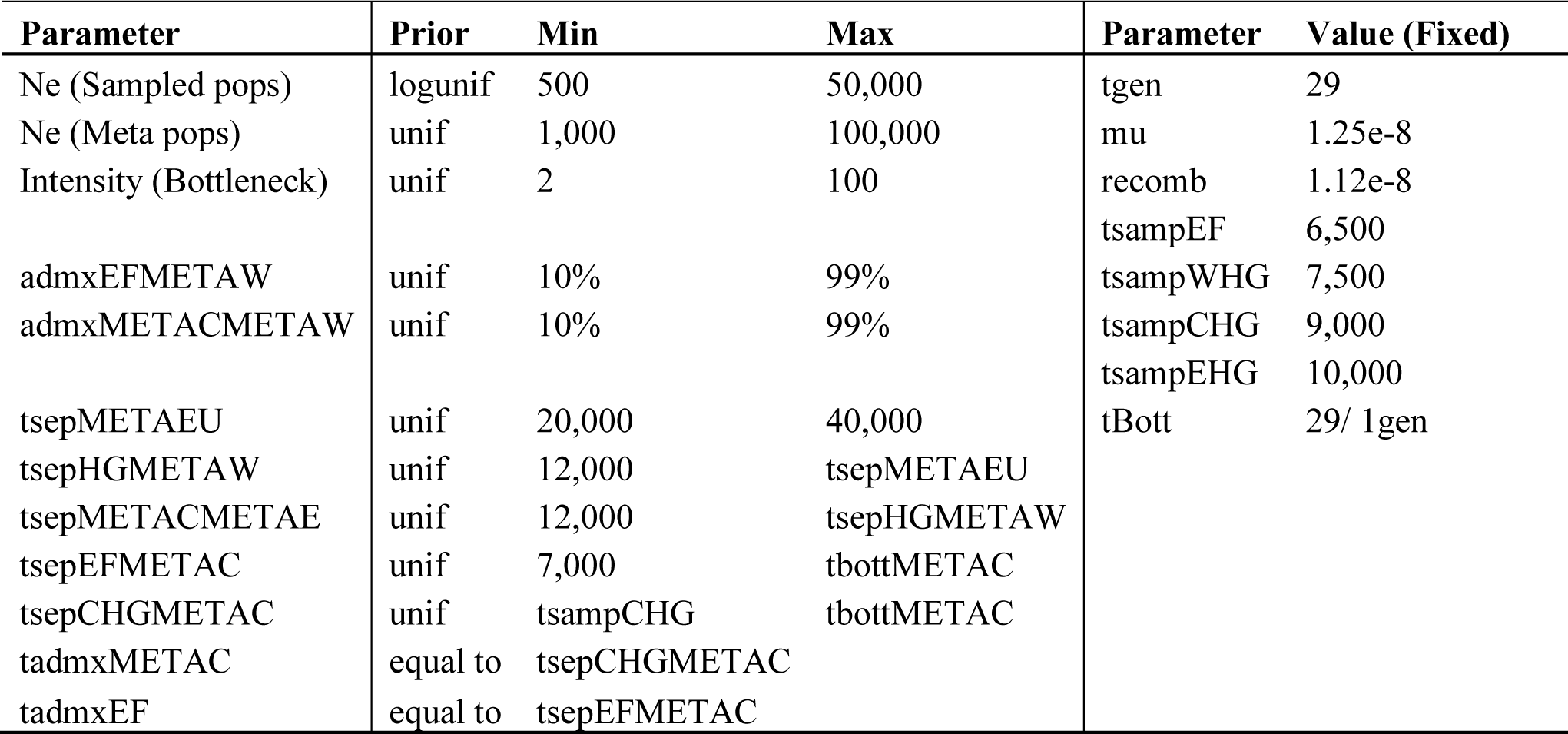
Demographic parameters and prior distributions of Model B. Mutation and Recombination rates are expressed per nucleotide per generation, Times are expressed in years.

| Parameter | Prior | Min | Max | Parameter | Value (Fixed) |
| --- | --- | --- | --- | --- | --- |
| Ne (Sampled pops) | logunif | 500 | 50,000 | tgen | 29 |
| Ne (Meta pops) | unif | 1,000 | 100,000 | mu | 1.25e-8 |
| Intensity (Bottleneck) | unif | 2 | 100 | recomb | 1.12e-8 |
|  |  |  |  | tsampEF | 6,500 |
| admxEFMETAW | unif | 10% | 99% | tsampWHG | 7,500 |
| admXMETACMETAW | unif | 10% | 99% | tsampCHG | 9,000 |
|  |  |  |  | tsampEHG | 10,000 |
| tsepMETAUE | unif | 20,000 | 40,000 | tBott | 29/ 1gen |
| tsepHGMETAW | unif | 12,000 | tsepMETAUE |  |  |
| tsepMETACMETAE | unif | 12,000 | tsepHGMETAW |  |  |
| tsepEFMETAC | unif | 7,000 | tbottMETAC |  |  |
| tsepCHGMETAC | unif | tsampCHG | tbottMETAC |  |  |
| tadmXMETAC | equal to | tsepCHGMETAC |  |  |  |
| tadmXEF | equal to | tsepEFMETAC |  |  |  |

We performed 100,000 simulations for each model with *ms*. All the ABC-RF estimates have been obtained using the functions *abcrf*, for model selection, and *regAbcrf*, for parameters estimation, employing a forest of 4,000 trees (Pudlo et al. 2016; Raynal et al. 2019). Before calculating the posterior probabilities of the most supported model and estimating its parameters, we evaluated the out-of-bag Classification Error (CE) and the proportion of True Positives (1-CE) as a measure of the power of the model selection procedure. To determine the power of our procedure in estimating the demographic parameters of the most supported model, we calculated the coefficient of determination (R^2^).

## Results

### Power Analysis Low-ABC

#### Model Selection

##### One-population models

The plots of Figure 1 report the results of the power analyses obtained summarizing the data through the FDSS estimated from genotype likelihoods (GLs). In each plot, we reported the True Positive (TP) rates for each demographic model and the combination of experimental parameters tested; plots in Panel A show the TP proportion obtained simulating 1,000 loci, whereas in Panel B those obtained simulating 5,000 loci.

For all three scenarios, we found the Low-ABC had very high True Positive rates, ranging from almost 80% to 100%, irrespective of the combination of parameters generating the data.

The proportion of true positives generally increase with the increase of both locus length and number of loci considered; the number of chromosomes does not seem to remarkably affect the true positives rate. These observations are consistent with the results detailed in Ghirotto et al. (2021).

Remarkably, for both *Bottleneck* and *Exponential Growth* models, the TP proportions obtained simulating the data at low coverage (1x, 2x and 5x) are comparable with those obtained for the high coverage (30x), ranging from 95% to 100% for almost all the combinations of parameters tested; differences are observed only for the *Bottleneck* model when considering 1,000 loci of 200bp. In this specific case the proportion of TP varies between 0.78 and 0.95. For the *Constant* model, the true positives percentage considering data with the lower coverage levels (about ∼80%-90% TP) is even higher than that obtained with a coverage level of 30x (∼80% TP); this difference is more pronounced when analyzing loci of 200bp.

##### Two-population models

The plots in Figure 2 show the results for the two-populations models. The proportion of TP is generally quite high, with the *Divergence with Migration* and the *Divergence with Admixture* models showing the highest proportion of TP, ranging from 62% to 90%. The true positives rate for the *Divergence* model is slightly lower, ranging from 60% to 78%. As observed for the one-population models, the proportion of TP increases as increasing locus length and number of loci. The results do not show significant differences in the percentage of true positives when simulating the data at different coverage levels.

#### Parameters Estimation

##### One-population models

Supplementary Tables 1-6 report the results of the quality assessment of the parameters estimation procedure for the one-population models. The R^2^, Bias, RMSE, Factor2 and 50% Coverage associated to each demographic parameter were estimated exploiting 1,000 pseudo-observed datasets (pods) and reference tables of 100,000 simulated datasets. In each box we listed the indices calculated for each coverage level (1x, 2x, 5x and 30x) together with the number of chromosomes and the locus lengths tested; in Supplementary Tables 1, 3 and 5 we reported all the combinations considering 1,000 loci, whereas in Supplementary Tables 2, 4 and 6 those considering 5,000 loci. Figure 4-6 report the distribution of the relative Bias valued over 1,000 pods as a general index quantifying the goodness of the estimates; we reported in these figures only the combinations of experimental parameters considering 50 chromosomes per population, as the quality of parameters estimates is not significantly affected by the number of chromosomes used.

The *Constant* model is defined by a single demographic parameter, i.e. the effective population size (N1); the quality assessment procedure indicates a good ability to estimates N1, with R^2^ values reaching 100% and median Bias values always close to 0 for all the combinations of experimental conditions tested. The variance of the estimates decreases when using more loci (Figure 4 left vs right panel) or when increasing their length. In general, the coverage level seems to not affect the median quality estimate measures, indeed the results obtained considering low-coverage levels (1x, 2x and 5x) are comparable with those obtained with a 30x coverage (Supplementary Tables 1-2 and Figure 4). However, the dispersion of the relative bias is considerably reduced for shorter loci (200bp in length) for coverage levels higher than 2x; for longer loci (1,000bp) a small dispersion of the relative bias was already observed at 1x coverage.

The quality estimates of the three demographic parameters defining the *Bottleneck* model (N1, NaBott and T) shows similar results with R^2^ values >50% and median Bias values ranging from 0.01 to 0.4. As regarding to the coverage levels, we observed a slight improvement of the estimates with the increase of the sequencing coverage (median Bias 30x: 0.01-0.2; median Bias 1x: 0.04-0.4). Even in this case, the dispersion of the relative Bias decreases with the increase of the number and length of loci across almost all the experimental conditions analyzed and among demographic parameters (Supplementary Tables 3-4 and Figure 5).

**Figure 4.**
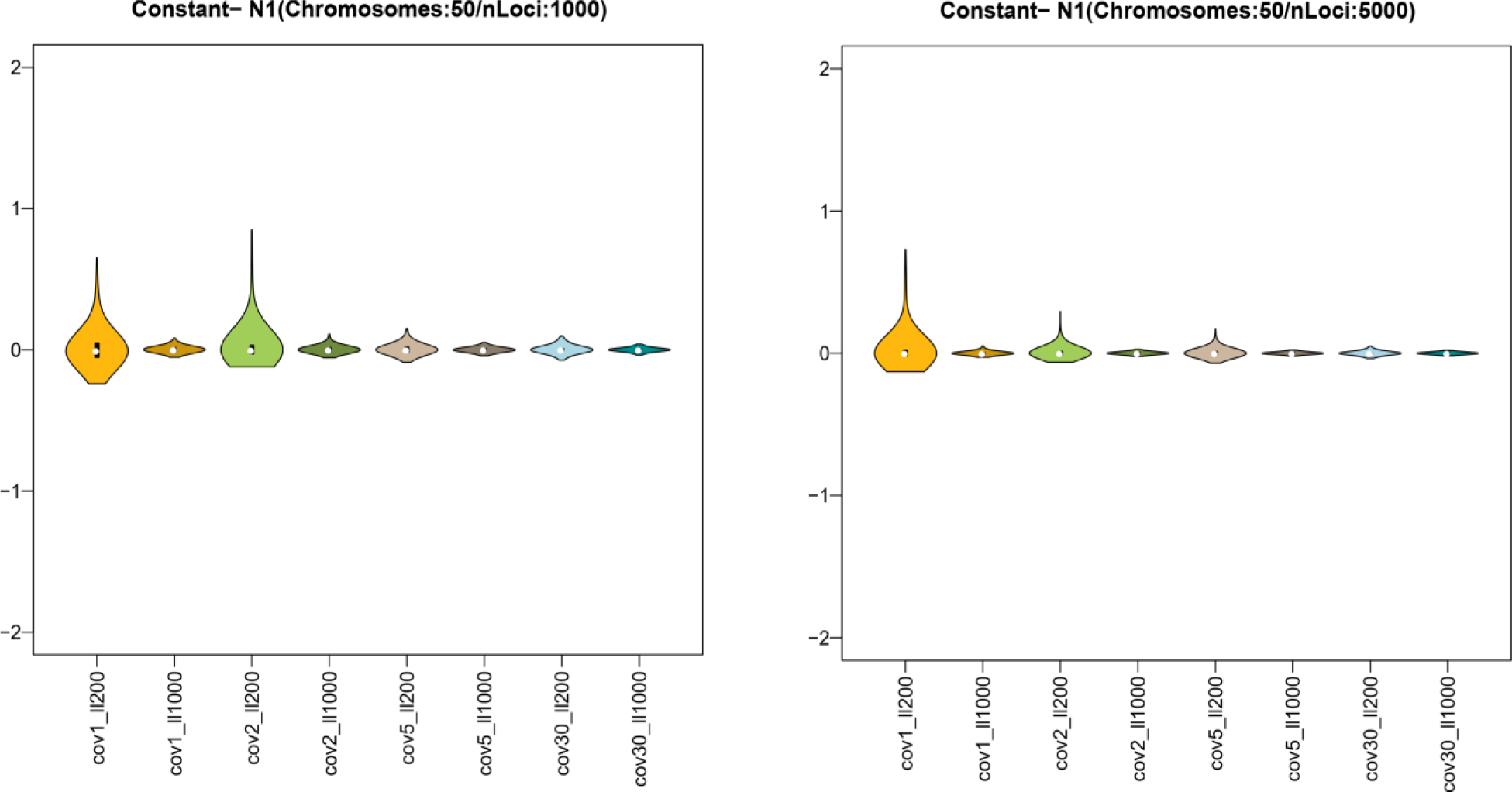
Relative Bias distributions for the Constant model’s demographic parameters. Coverage levels are represented by different colors. Lighter colors indicate short loci (200bp), darker colors indicate longer loci (1,000bp). Left panel: combinations considering 1,000 loci. Right panel: combinations considering 5,000 loci.

**Figure 5.**
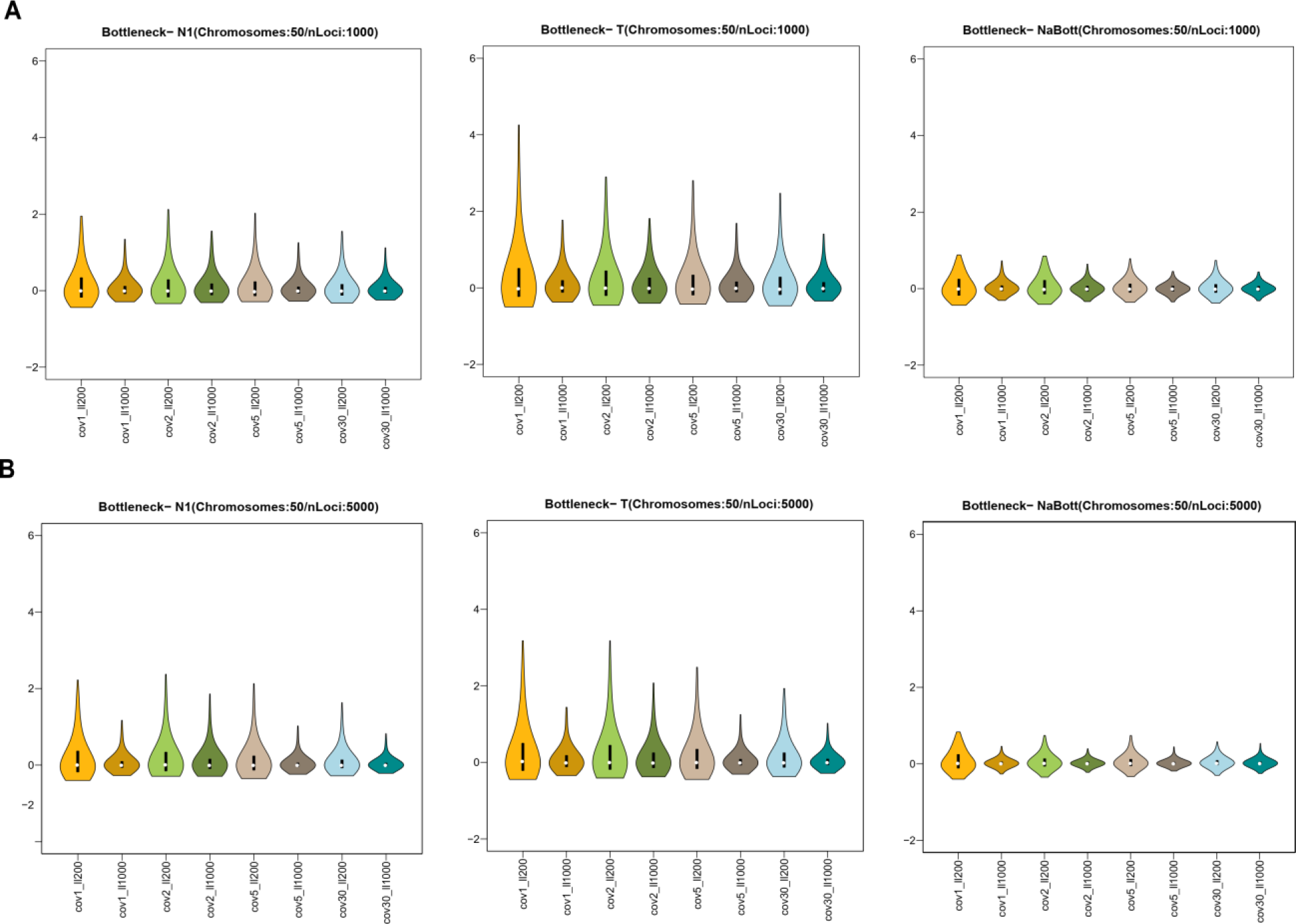
Relative Bias distributions for the Bottleneck model’s demographic parameters. These plots have the same features of Figure 4. A) Combinations considering 1,000 loci. B) Combinations considering 5,000 loci.

For the *Exponential Growth* model, we observed the same general results with R^2^ >10% and median Bias that varies between 0.06 and 0.5 for most of the combinations of experimental conditions tested. The demographic parameter that shows lower quality indices is the current effective population size (N1) with R^2^ values ranging from 2% to 30% depending on the experimental conditions; in particular, the R^2^ is lower than 10% when considering only 1,000 loci and 10 sampled chromosomes (Supplementary Tables 5-6). We did not observe significant differences in the quality of the estimates with respect to the coverage levels analyzed (Supplementary Tables 5-6 and Figure 6).

**Figure 6.**
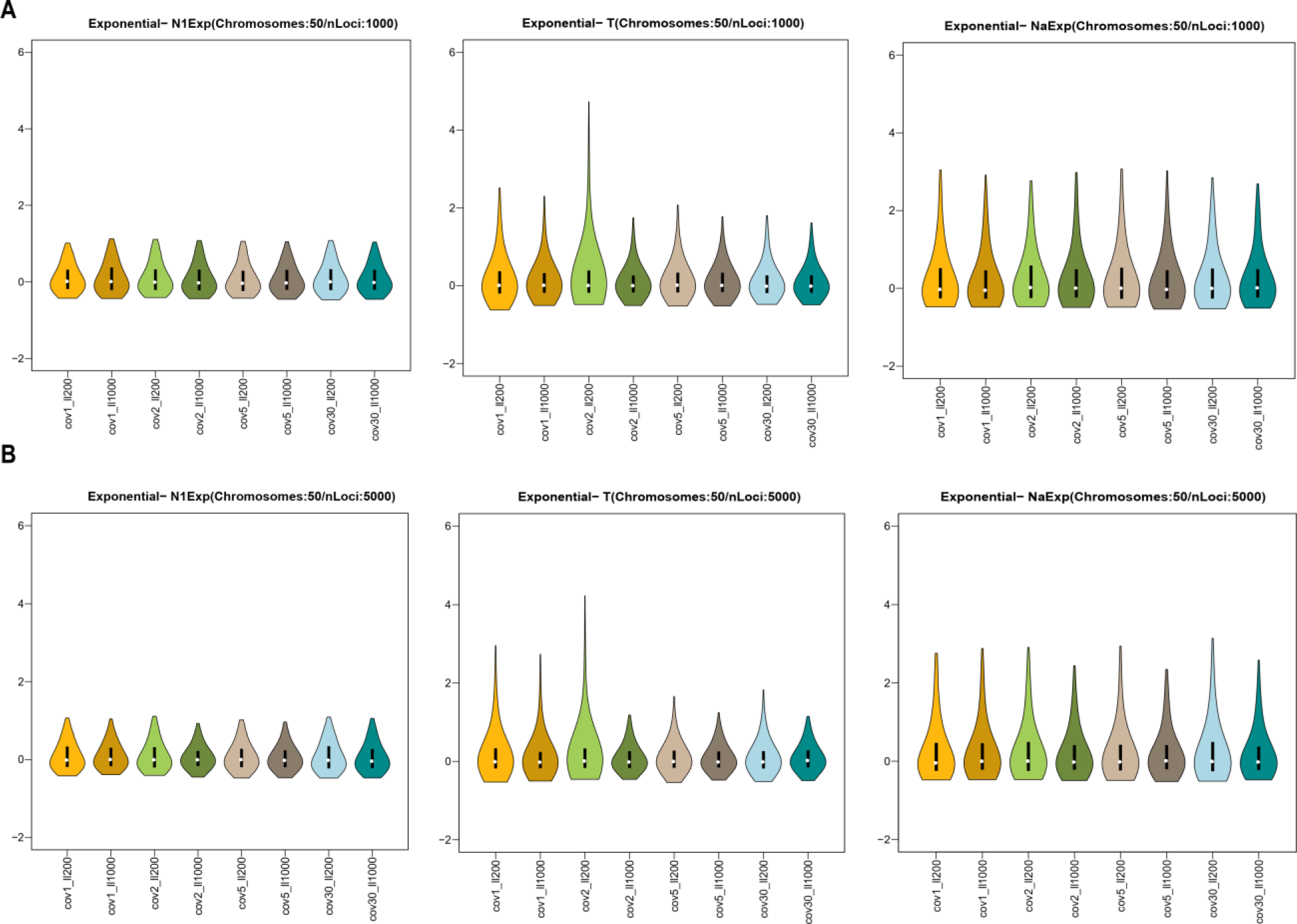
Relative Bias distributions for the Exponential Growth model’s demographic parameters. These plots have the same features of Figure 4. A) Combinations considering 1,000 loci. B) Combinations considering 5,000 loci.

##### Two-population models

Supplementary Tables 7-12 show the results of the quality assessment of the parameters estimation procedure for the two-populations models. Figures 7-9 report the distribution of the relative Bias values over the 1,000 pods, for the combinations of experimental parameters considering 50 chromosomes.

**Figure 7.**
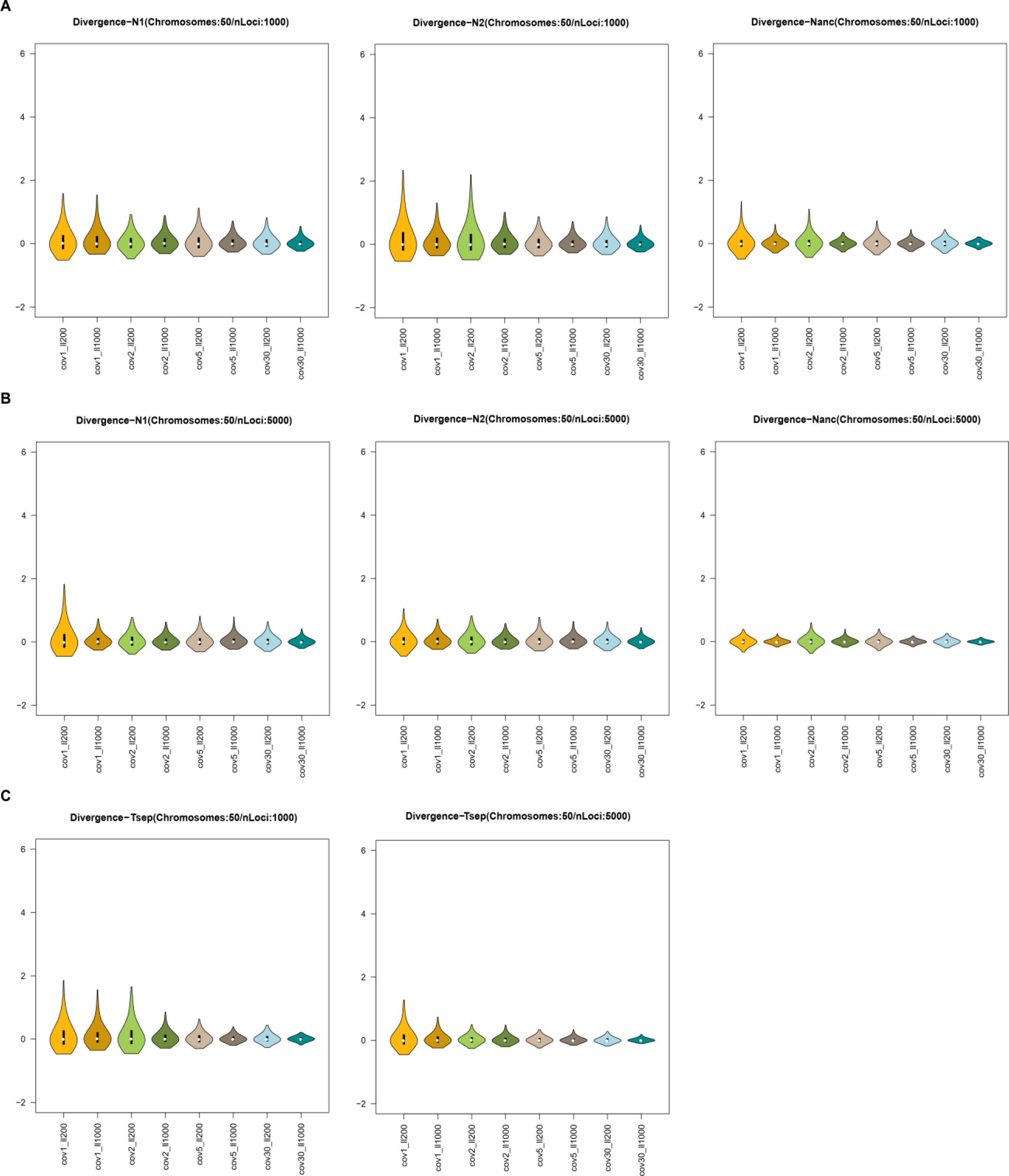
Relative Bias distributions for the Divergence model’s demographic parameters. These plots have the same features of Figure 4. Panel A and left of Panel C: combinations considering 1,000 loci. Panel B and right of Panel C: combinations considering 5,000 loci.

The general results of the quality of the estimates are similar to those obtained for the one-population models, with R^2^ > 50% and median Bias values ranging from 0.003 to 0.4 for most of the demographic parameters estimated (Supplementary Tables 7-12). The estimates improve with the increase of the number of chromosomes, number and length of loci considered in the analysis (Figures 7-9). As for the one-population models, the effect of the coverage level on the quality of the estimates is negligible, the results remain indeed consistent regardless the sequencing coverage (Figures 7-9).

The model showing the best quality indices is the *Divergence* model (Figure 7). In this scenario the demography is defined by four demographic parameters: the effective population sizes of the ancestral population (Nanc) and of the two derived populations (N1 and N2) and the divergence time (Tsep). All these parameters show R^2^ values ranging from 65% to almost 100% and low median Bias (Supplementary Tables 7-8).

The *Divergence with Migration* model shows lower quality indices compared to the other two-population models. The R^2^ of the three effective population sizes varies between 40% and almost 100%, whether the median Bias’s values vary between 0.06 and 0.4. The worst estimated parameters are the two migration rates (m12 and m21) with R^2^ < 40% and higher levels of Bias with values ranging from 1 to 34 (Supplementary Tables 9-10). In general, the two migration rates and the separation time (Tsep), show greater levels of variance in the dispersion of the relative Bias that slightly improves with the increase of the sequencing coverage, the number and length of loci (Figure 8).

**Figure 8.**
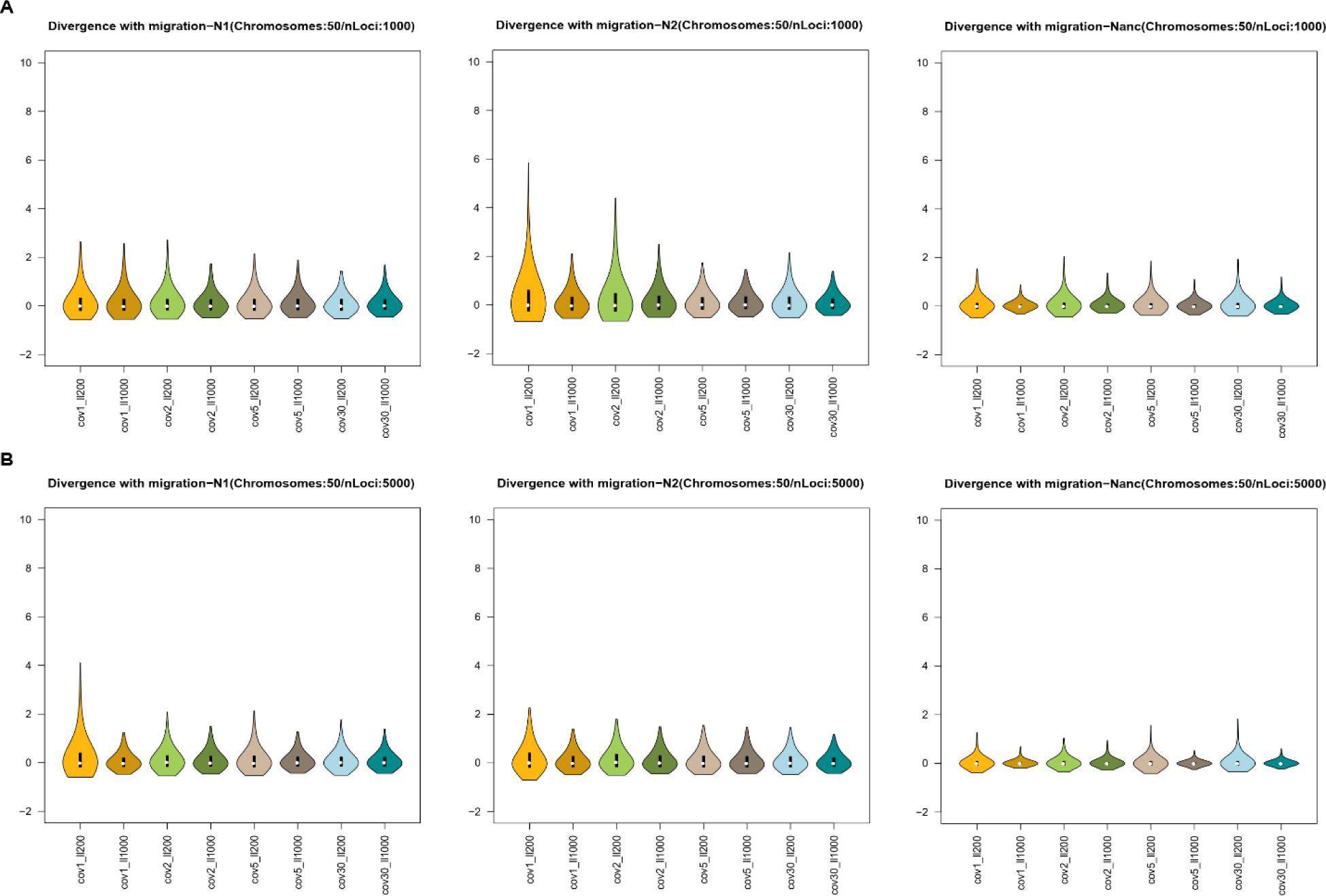
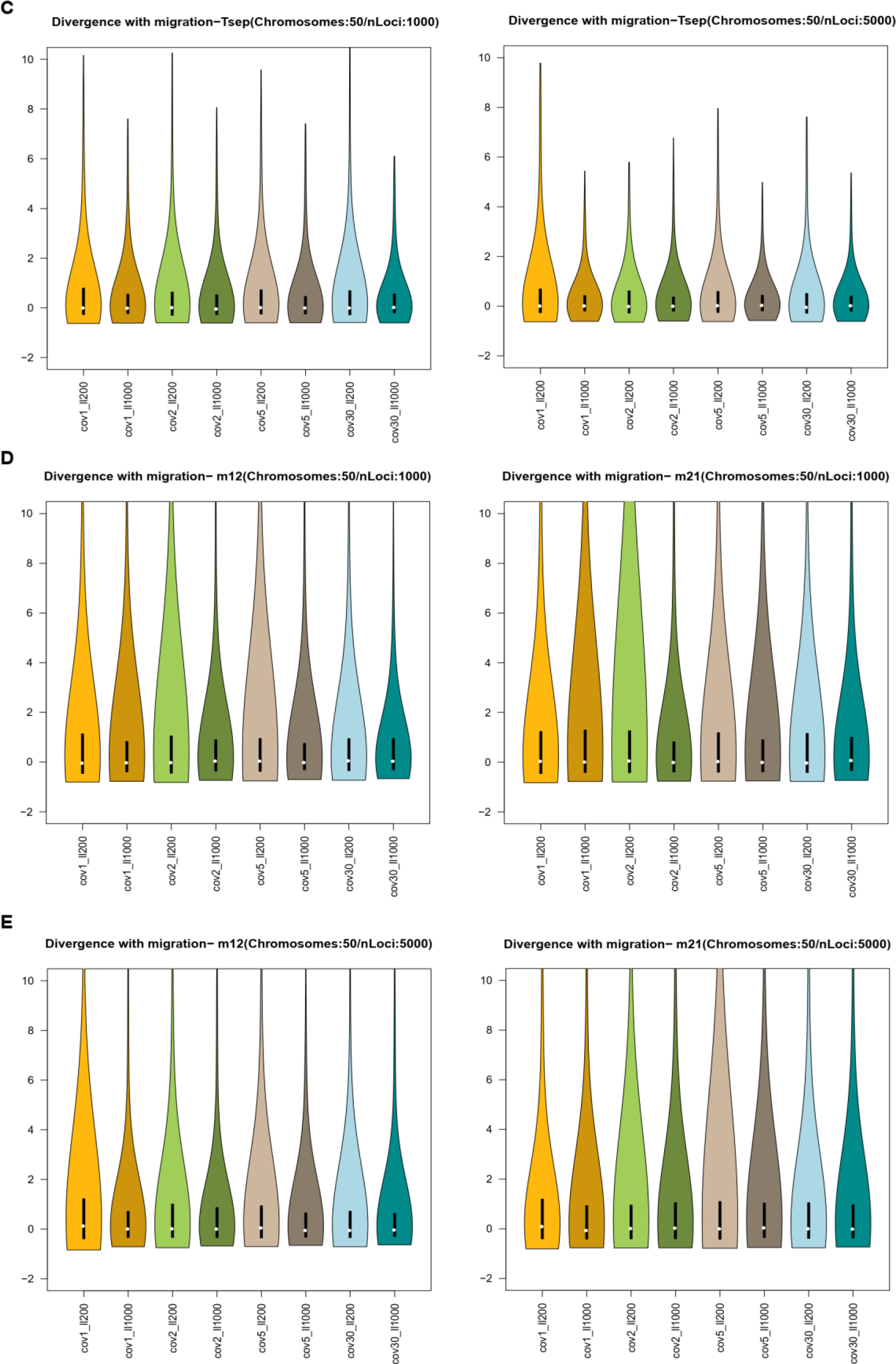
Relative Bias distributions for the Divergence with migration model’s demographic parameters. These plots have the same features of Figure 4. Panels A, C and D: combinations considering 1,000 loci. Panels B, C and E: combinations considering 5,000 loci.

Finally, for the *Divergence with Admixture* model we observed good quality indices for the three population sizes (N1, N2 and Nanc), the time of the admixture event (Tadm) and the divergence time (Tsep) with R^2^> 50% and low level of median Bias with values ranging from 0.005 to 0.1. The worst estimated parameters are the two admixture rates (adm12 and adm21) with R^2^ <10% for most of the combination of experimental parameters tested (Supplementary Tables 11-12). Even in this case, the variance of the relatives Bias decrease with the increase of locus length, number of loci and coverage levels (Figure 9); this pattern is more evident for the three effective population sizes than for the divergence time (Tsep), the admixture time (Tadm) and the two admixture rates (Figure 9).

**Figure 9.**
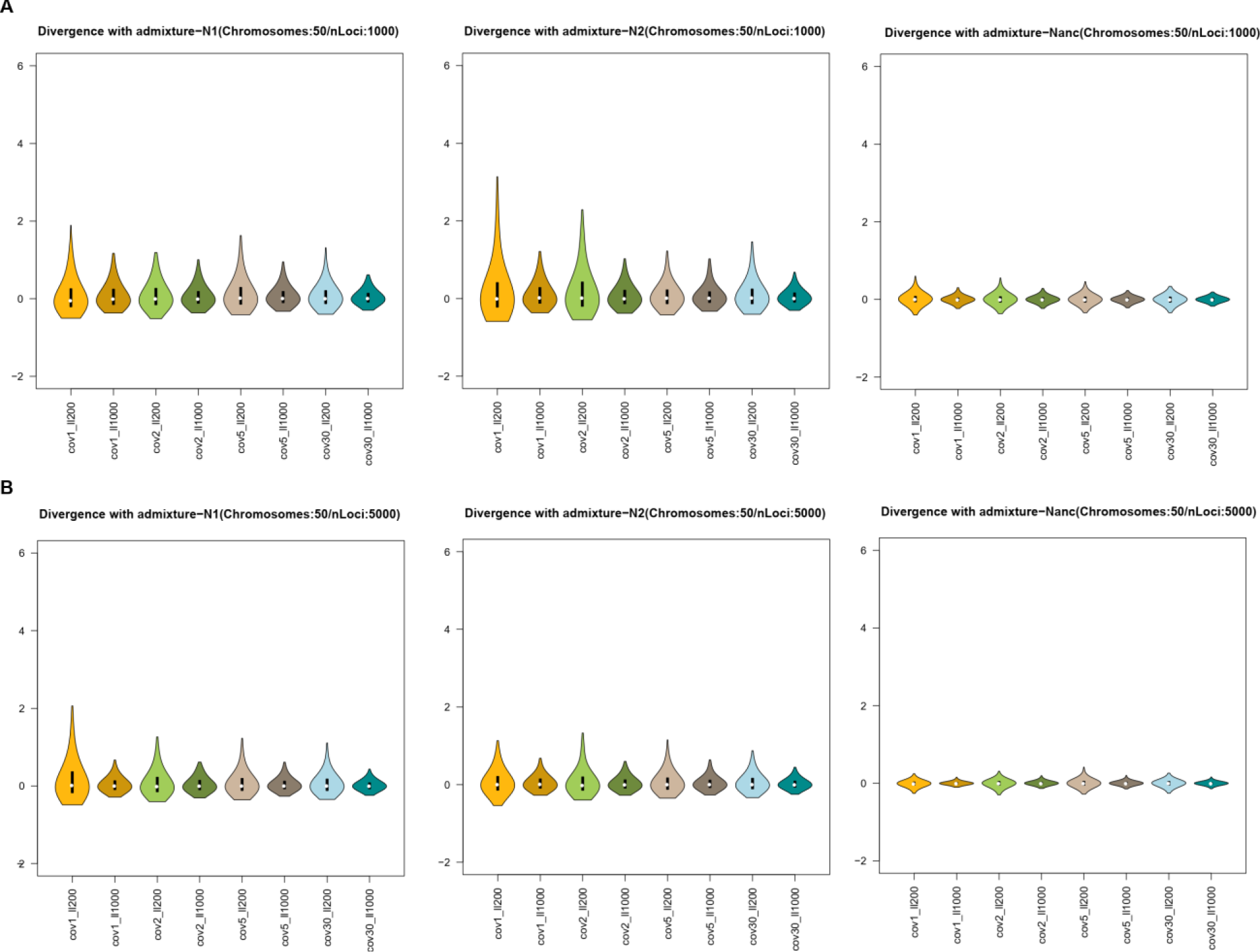
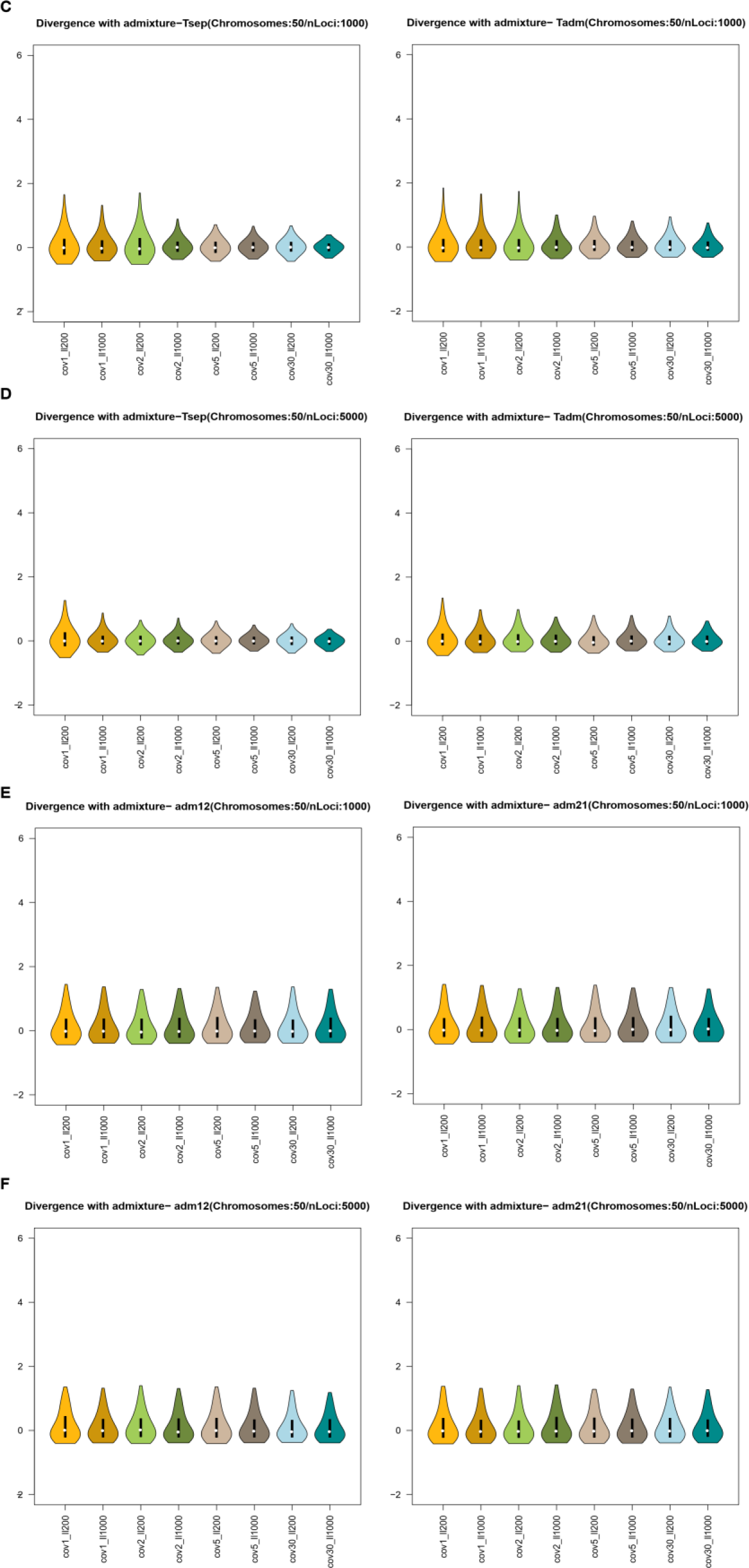
Relative Bias distributions for the Divergence with admixture model’s demographic parameters. These plots have the same features of Figure 4. Panels A, C and E: combinations considering 1,000 loci. Panels B, D and F: combination considering 5,000 loci.

#### Comparison with using exact simulations and GL-based summary statistics from ANGSD

In Supplementary Figures 1 and 2 we reported the results obtained from the power analysis conducted to evaluate the effectiveness, at different coverage levels, of considering the uncertainty in genotype calling by calculating observed statistics using GLs and directly comparing them with simulated data. In each figure, we reported the True Positive (TP) rates for each model and combination of parameters tested: Panel A shows the TP proportion obtained by simulating 1,000 genomic loci, while Panel B presents the TP rate obtained through simulations involving 5,000 loci.

We explored two sets of models, namely the one-population and two-population models. For both sets of models, the datasets generated with a 30x coverage level were assigned to the correct demographic history with TP rates ranging from 70% to nearly 100%.

When comparing the one-population models considering a coverage of 5x, the proportion of TP decreases from ∼80-97% to 0-20% for both the *Exponential* and the *Constant* model. Furthermore, the TP proportion significantly declines when analyzing lower coverage levels (1x and 2x), resulting in a TP rate of 0%. For the *Bottleneck* model, we observed a higher proportion of TPs, equal to ∼90-100%, regardless of the coverage level considered (Supplementary Figure 1).

When comparing the two-population models considering a coverage of 5x we observed a slight reduction in the power for the *Divergence* model, with a TP rate declining from ∼65-80% to ∼60-80%. The TP rate further decreases with lower coverage levels, 1x and 2x (0-40%). For the *Divergence with Migration* model, we observed a decrease in TP proportion from ∼80-90% to 20-80% when analyzing data at 5x coverage. The TP rate further decreases with 1x and 2x coverage levels (0-20%). The *Divergence with Admixture* model, instead, showed higher TP proportions, with values ranging from 60% to 90% regardless of the coverage level considered (Supplementary Figure 2).

### Real Case: Mesolithic hunter-gatherer and Early Farmers relationships

Table 6 reports the results of the power test for the comparison between the two alternative demographic scenarios under investigation. For both models, we obtained a Classification Error of ∼2% and a proportion of True Positives of ∼98%, showing that our procedure is effective in discriminating between the two models. The model selection procedure supported Model A as the scenario that best explains the evolutionary relationships between Early Farmers and Mesolithic Hunter-Gatherers, with a posterior probability of 70% (Table 6).

**Table 6.** Model selection. Best model selected (highlighted in bold) by the ABC-RF procedure.

|  | Classification Error (%) | True Positives (%) | Votes | Posterior Probability |
| --- | --- | --- | --- | --- |
| **Model A** | **2.25** | **97.75** | **2,439** | **70%** |
| Model B | 2.31 | 97.69 | 1,561 |  |

Once Model A was identified as the most probable, we moved to estimate its parameter values, maximizing the fit between observed and simulated genomic data. The estimated posterior distributions for all parameters are shown in Table 7. In general, the demographic parameters were estimated reasonably well, with most R^2^ values above 10%.

**Table 7.**
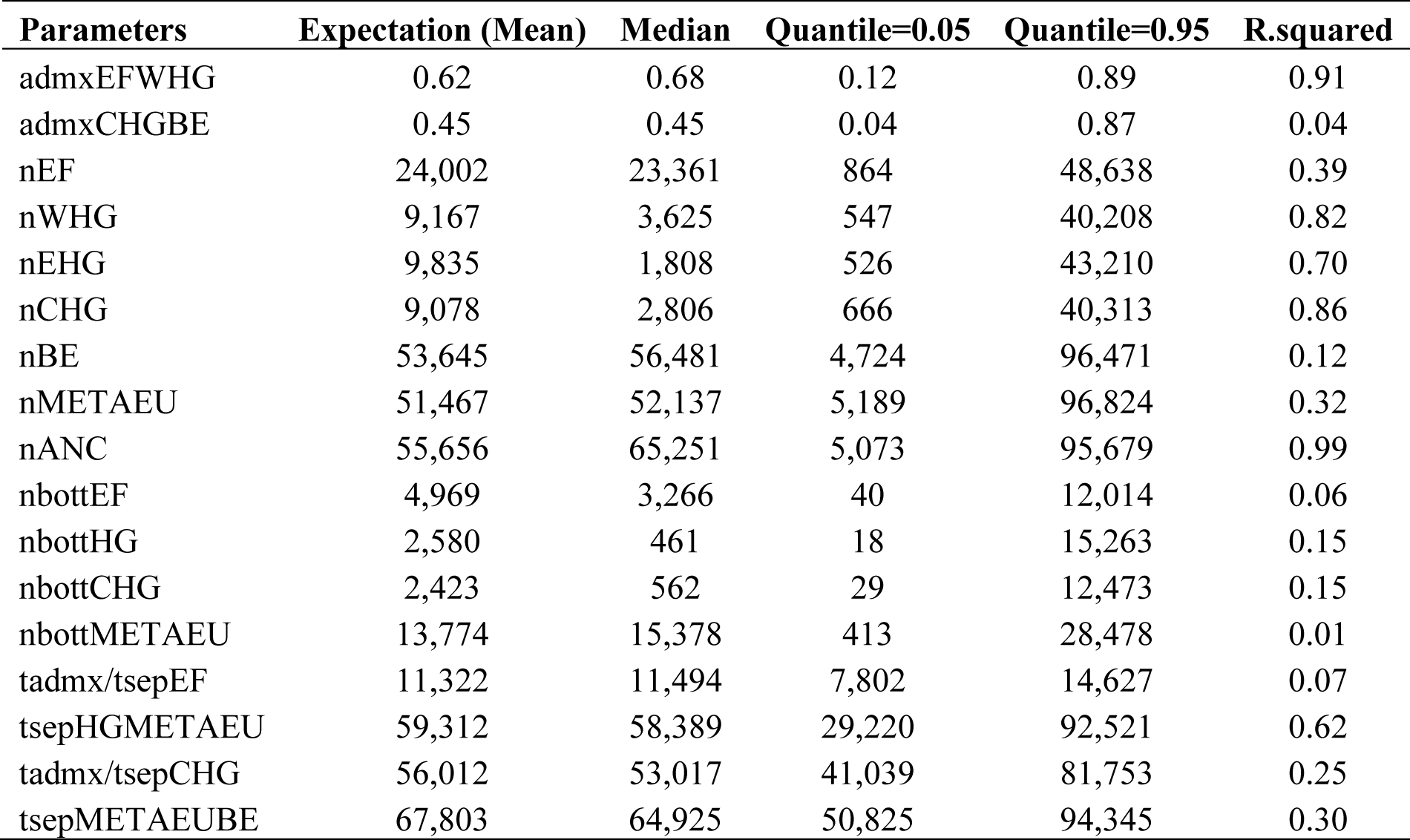
Estimated parameters of the most supported model (Model A).

The divergence time between the Basal Eurasia and the Meta-Eurasian population was estimated at 64,925 years ago (95% CI: 50,825-94,345; R^2^=0.30). The ancestral population of western and eastern HGs diverged from the Meta-Eurasian population 58,389 years ago (95% CI: 29,220-92,521; R^2^=0.62), whereas the separation time of the Caucasus hunter-gatherers was estimated at 53,017 years ago (95% CI: 41,039-81,753; R^2^=0.25). We also estimated that the EFs received 68% of their ancestry from the Western HGs (95% CI: 12%-89%; R^2^=0.91). We could not confidentially estimate the Basal Eurasian contribution to the Caucasus HGs gene pool (∼45%; 95% CI: 5%-87%; R^2^=0.04) and the Anatolian/European Early Farmers’ divergence time from the Basal Eurasian population (∼11,494 years ago; 95% CI: 7,802-14,627; R^2^=0.07).

## Discussion

When analyzing low-coverage genomes in a population genetic context we should consider the high genotype uncertainty and the low ability to identify sequencing errors (Korneliussen et al. 2014; Kousathanas et al. 2017; Meisner and Albrechtsen, 2018). Unfortunately, statistical inference methods able to exploit low-coverage genomes in an inferential context with the aim of making accurate demographic inference of complex models, are still missing.

A possible strategy might be the incorporation of low coverage data within an Approximate Bayesian Computation (ABC) context, integrating the uncertainty related to the sequencing coverage level when computing the observed summary statistics; however, our simulation experiments show that the use of GLs-corrected summary statistics within ABC is not always effective. In our power analysis, indeed, this approach failed most of the time to recognize the true demographic history for sequencing coverage levels equal to or below 5x, regardless of the combination of experimental parameters tested. When comparing the one-population models, the *Bottleneck* model showed, instead, a high proportion of TP even at the lower coverage levels; however, we verified that this unexpected behaviour is caused by a higher number of sites erroneously identified as heterozygous at low coverage levels. This biased prediction of genomic variation causes a distortion in the summary statistic distribution towards a pattern of variation more easily generable from the *Bottleneck* model. All the low-coverage pods generated through the *Bottleneck* model were hence correctly assigned, thus explaining the high TP proportion observed for this scenario.

When comparing the two-population models, only the *Divergence with Admixture* model showed good TP proportions, with values ranging from 60% to 90%. Even in this case with very low-coverage levels, 1-2x, the genotype uncertainty prevents indeed the classification of private polymorphic sites. This results in an artificially higher proportion of loci containing shared polymorphic sites with respect to sites belonging to the other three categories, that is what is expected under the *Divergence with Admixture* model. Such distortion results in a higher proportion of pods assigned to the *Divergence with Admixture* model, and consequently in a high TP rate when pods actually come from the same model. Taken together, these results highlight that the correction of low coverage observed data through the method embedded in ANGSD to perform ABC model choice, does not produce reliable results when the sequencing coverage is 5x or less. This drawback severely limits the possibility to exploit this approach to study past demographic processes through the analysis of genomic data from high degraded samples, as those extracted from ancient remains (whose achievable coverage is often lower than 5x) or through non-invasive sampling.

To implement the use of low sequencing coverage genomic data in the ABC inferential process, we developed Low-ABC, a new inferential framework in which, rather than correcting the low-covered observed data, coalescent simulations are generated according to a specific coverage level. Observed and simulated data are hence directly comparable, and summary statistics are calculated in both cases through genotype likelihoods.

We evaluated the inferential power of low-ABC in distinguishing among different demographic models and in inferring model parameters under different experimental conditions. When we compared one-population scenarios, the model identifiability, calculated as the proportion of TPs over 100,000 pods, reached values between 80% and 100%, regardless of the coverage level considered. The true positives rates obtained simulating the data at a low sequencing coverage (1x to 5x) are comparable with those obtained simulating the data at 30x coverage. The proportion of true positives for the *Constant* model, considering data with the lower coverage levels, is slightly higher than that obtained with a coverage level of 30x (∼80%-90% vs ∼80% TP). This result may be related to the skewed level of polymorphism typical of low-coverage conditions (Nielsen et al. 2012; Fumagalli et al. 2013) that may amplify differences in the polymorphism levels generated by the models, especially in some regions of the parameter’s space. This behaviour is not observed in the *Bottleneck* and the *Exponential Growth* models, suggesting that the performance of the inferential framework at lower coverage levels could be model dependent and hence a careful analysis of the model choice performance should be always performed before analysing real datasets.

Among the two-population models, the True Positive (TP) proportions ranged from 60% to 90%, irrespective of the coverage level. Remarkably, when simulating data at low sequencing depths (1x to 5x), the true positive rates are always comparable to those obtained at a higher coverage (30x), especially when simulating 5,000 genomic loci of 1,000 base pairs length.

We then assessed for both sets of models the power of our framework in estimating the models’ parameters. The overall quality of the estimates was quite good, with high R^2^ values, low Bias and RMSE, and consequently high values of factor2 and 50% coverage. Once again, the quality of the estimates is not influenced by the coverage, indeed, the performances of the estimation process are comparable regardless of whether the coverage was 1x or 30x. The only case in which we observed lower quality indices was when we estimated the current effective population size (N1) under the one-population *Exponential Growth* model. These results were somehow expected: the ability to accurately estimates the present effective population size of an exponentially growing population strictly depends on the time of the beginning of the growth; if the growth start in recent times, we need larger samples to characterise the effective population size before and after the expansion (Boitard et al. 2016). Among the two-population models, the estimated demographic parameters showed good quality indices, except for both migration and admixture rates, two classes of demographic parameters known to be difficult to estimate (Robinson et al. 2014).

Increasing the number of individuals sampled from each population and the number of genomic loci, we observed a general improvement of the quality of the estimates with 5000 loci characterised in 10 chromosomes per population, showing a good balance between sampling effort and precision/accuracy of the estimates.

We finally applied Low-ABC to the analysis of two demographic models depicting alternative historical relationships between Early Farmers and Mesolithic Hunter-Gatherers. Under the selected model (posterior probability of 70%), all the HGs groups diverged from an ancestral population shortly after the human expansion into Europe, approximately 58,000 years ago, whereas the Anatolian/European early Farmers descended from the Basal Eurasia population and admixed with the Western HGs during the Neolithic expansion (approximately 11,000 years ago), consistent with findings from previous research (Jones et al. 2015; Lazaridis et al. 2016b; Wang et al. 2019). We also estimated the demographic parameters of the supported model. Many of the estimated parameters exhibit R^2^ values above 10%; however, their confidence intervals are wide, and their posterior distributions often reflect the prior range (Table 7). The uncertainty arising from the parameter estimation procedure may reflect the limitations of the highly heterogeneous genomic dataset used in this study. Many of these genomes were not subjected to UDG treatment, leading to an increased rate of sequencing errors in the untreated sequences compared to the treated genomes while our procedure assumes a uniform sequencing error rate across all simulated individuals.

The Low-ABC framework presented here paves the ground for a new era of effective comparisons among alternative evolutionary and demographic histories through the analysis of samples sequenced at low coverage. The impressive amount of ancient genomic sequences produced at a coverage level of at least 1x can now be robustly integrated for the first time within an ABC framework to explicitly test hypotheses that arose from descriptive analyses of sequences’ variation, as to the peopling dynamics of the Americas (Willerslev and Meltzer 2021) or the characterization of genetic shifts and regionally-specific patterns of admixture across western Eurasia (Fernandes et al. 2020; Feldman et al. 2021).

The presented approach may also be used to disentangle past demographic dynamics of cryptic or elusive species, where the lack of availability of samples may only lead to low-coverage sequencing, or to significantly reduce sequencing costs of projects requiring extensive sampling and sequencing, as those aiming to determine the worldwide colonization routes of invasive species.

## Author contributions

A.B. conceived the study; A.B., S.G. and A.M. designed the study; P.M.D. and L.C., collected and processed the ancient whole-genome data; M.T.V. analyzed the data; M.T.V., A.M, S.G, R.B.A. and A.B. discussed the results. M.T.V., and S.G. wrote the paper with contributions from all authors; A.B. and A.M. supervised research.

## Data and resource availability

The data underlying this article are available in the Zenodo repository, at http//[link to the repository will be added upon acceptance].

## Acknowledgements

Maria Teresa Vizzari’s, Rajiv Boscolo Agostini’s and Silvia Ghirotto’s contribution to this research was financially supported by PRIN 2017 (grant 20177PJ9XF), and PRIN 2020 (grant 2020TACEZR) from the Italian Ministry of Education, University and Research (MIUR). Andrea Benazzo’s contribution to this research was financially supported by PRIN 2022 from the Italian Ministry of Education, University and Research (MIUR), funded by the European Union -

NextGenerationEU -mission 4, component 2, investment 1.1. - project 202292P4R7 - CUP F53D23003960006.

