## Supplementary files for "Low-ABC: a robust demographic inference from low-coverage whole-genome data through ABC"

**Supplementary Table 1. Accuracy of the estimated parameters of the Constant model assessed by 1,000 pods.** Combinations of experimental parameters considering 1,000 loci. The number of chromosomes is indicated with *nc*, whereas *ll* indicates the locus length.

| | | Coverage 1x | | | | | |
|---|---|---|---|---|---|---|---|
| | | Parameter | $R^2$ | Bias | RMSE | Factor2 | Coverage50% |
| nc10 | ll200 | *N1* | 0.955 | 0.033 | 1545.833 | 0.989 | 0.644 |
| | ll1000 | | 0.994 | 0.002 | 766.837 | 1.000 | 0.570 |
| nc20 | ll200 | | 0.984 | 0.016 | 1308.498 | 0.995 | 0.603 |
| | ll1000 | | 0.999 | 0.004 | 609.009 | 1.000 | 0.554 |
| nc50 | ll200 | | 0.990 | 0.036 | 1713.427 | 0.983 | 0.589 |
| | ll1000 | | 1.000 | 0.005 | 503.096 | 1.000 | 0.615 |

| | | Coverage 2x | | | | | |
|---|---|---|---|---|---|---|---|
| | | Parameter | $R^2$ | Bias | RMSE | Factor2 | Coverage50% |
| nc10 | ll200 | *N1* | 0.997 | 0.033 | 1499.297 | 0.985 | 0.573 |
| | ll1000 | | 0.996 | 0.002 | 617.687 | 1.000 | 0.594 |
| nc20 | ll200 | | 0.992 | 0.017 | 1247.473 | 0.992 | 0.609 |
| | ll1000 | | 0.998 | 0.001 | 511.822 | 1.000 | 0.600 |
| nc50 | ll200 | | 0.966 | 0.079 | 1175.160 | 0.979 | 0.755 |
| | ll1000 | | 0.999 | 0.007 | 472.530 | 0.999 | 0.618 |

| | | Coverage 5x | | | | | |
|---|---|---|---|---|---|---|---|
| | | Parameter | $R^2$ | Bias | RMSE | Factor2 | Coverage50% |
| nc10 | ll200 | *N1* | 1.000 | 0.013 | 1211.403 | 0.996 | 0.533 |
| | ll1000 | | 1.000 | 0.000 | 518.848 | 1.000 | 0.663 |
| nc20 | ll200 | | 1.006 | 0.011 | 1167.460 | 0.994 | 0.531 |
| | ll1000 | | 0.996 | 0.002 | 548.235 | 1.000 | 0.609 |
| nc50 | ll200 | | 1.000 | 0.014 | 775.294 | 0.995 | 0.667 |
| | ll1000 | | 0.999 | 0.000 | 440.270 | 1.000 | 0.619 |

| | | Coverage 30x | | | | | |
|---|---|---|---|---|---|---|---|
| | | Parameter | $R^2$ | Bias | RMSE | Factor2 | Coverage50% |
| nc10 | ll200 | *N1* | 0.998 | 0.001 | 1188.937 | 1.000 | 0.508 |
| | ll1000 | | 0.995 | 0.002 | 549.645 | 1.000 | 0.558 |
| nc20 | ll200 | | 0.994 | 0.001 | 920.206 | 1.000 | 0.561 |
| | ll1000 | | 0.999 | 0.001 | 499.945 | 1.000 | 0.577 |
| nc50 | ll200 | | 1.000 | 0.004 | 700.686 | 1.000 | 0.620 |
| | ll1000 | | 1.000 | 0.000 | 382.375 | 1.000 | 0.591 |

**Supplementary Table 2. Accuracy of the estimated parameters of the Constant model assessed by 1,000 pods.** Combinations of experimental parameters considering 5,000 loci.

| | | Coverage 1x | | | | | |
|---|---|---|---|---|---|---|---|
| | | Parameter | $R^2$ | Bias | RMSE | Factor2 | Coverage50% |
| nc10 | ll200 | *N1* | 1.000 | 0.007 | 590.230 | 1.000 | 0.565 |
| | ll1000 | | 0.999 | 0.001 | 334.277 | 1.000 | 0.570 |
| nc20 | ll200 | | 0.996 | 0.005 | 622.665 | 0.999 | 0.590 |
| | ll1000 | | 0.998 | 0.002 | 285.148 | 1.000 | 0.554 |
| nc50 | ll200 | | 0.989 | 0.053 | 805.546 | 0.979 | 0.571 |
| | ll1000 | | 0.998 | 0.010 | 246.237 | 0.997 | 0.669 |

| | | Coverage 2x | | | | | |
|---|---|---|---|---|---|---|---|
| | | Parameter | $R^2$ | Bias | RMSE | Factor2 | Coverage50% |
| nc10 | ll200 | *N1* | 0.996 | 0.008 | 524.767 | 0.999 | 0.605 |
| | ll1000 | | 1.000 | 0.001 | 307.994 | 1.000 | 0.610 |
| nc20 | ll200 | | 0.999 | 0.012 | 482.244 | 0.998 | 0.647 |
| | ll1000 | | 0.999 | 0.000 | 268.772 | 1.000 | 0.594 |
| nc50 | ll200 | | 0.995 | 0.020 | 455.324 | 0.994 | 0.670 |
| | ll1000 | | 1.000 | 0.002 | 229.755 | 1.000 | 0.575 |

| | | Coverage 5x | | | | | |
|---|---|---|---|---|---|---|---|
| | | Parameter | $R^2$ | Bias | RMSE | Factor2 | Coverage50% |
| nc10 | ll200 | *N1* | 1.000 | 0.003 | 462.882 | 1.000 | 0.595 |
| | ll1000 | | 0.998 | 0.000 | 253.277 | 1.000 | 0.580 |
| nc20 | ll200 | | 1.001 | 0.003 | 417.779 | 1.000 | 0.592 |
| | ll1000 | | 1.000 | 0.001 | 229.886 | 1.000 | 0.638 |
| nc50 | ll200 | | 0.998 | 0.012 | 450.187 | 0.996 | 0.596 |
| | ll1000 | | 1.000 | 0.000 | 208.397 | 1.000 | 0.597 |

| | | Coverage 30x | | | | | |
|---|---|---|---|---|---|---|---|
| | | Parameter | $R^2$ | Bias | RMSE | Factor2 | Coverage50% |
| nc10 | ll200 | *N1* | 0.999 | 0.002 | 408.336 | 1.000 | 0.643 |
| | ll1000 | | 0.999 | 0.001 | 267.408 | 1.000 | 0.606 |
| nc20 | ll200 | | 1.000 | 0.001 | 379.749 | 1.000 | 0.555 |
| | ll1000 | | 0.999 | 0.000 | 228.323 | 1.000 | 0.646 |
| nc50 | ll200 | | 0.999 | 0.002 | 316.463 | 1.000 | 0.555 |
| | ll1000 | | 0.998 | 0.000 | 215.947 | 1.000 | 0.625 |

**Supplementary Table 3. Accuracy of the estimated parameters of the Bottleneck model assessed by 1,000 pods.** Combinations of experimental parameters considering 1,000 loci.

| | | Coverage 1x | | | | | |
|---|---|---|---|---|---|---|---|
| | | Parameter | $R^2$ | Bias | RMSE | Factor2 | Coverage50% |
| nc10 | ll200 | *N1* | 0.653 | 0.168 | 868.519 | 0.932 | 0.552 |
| | | *T* | 0.655 | 0.277 | 3695.004 | 0.875 | 0.538 |
| | | *NaBott* | 0.597 | 0.059 | 14885.458 | 0.976 | 0.515 |
| | ll1000 | *N1* | 0.795 | 0.100 | 728.413 | 0.968 | 0.495 |
| | | *T* | 0.861 | 0.149 | 2522.540 | 0.939 | 0.521 |
| | | *NaBott* | 0.819 | 0.021 | 9948.504 | 0.994 | 0.626 |
| nc20 | ll200 | *N1* | 0.660 | 0.156 | 853.297 | 0.941 | 0.527 |
| | | *T* | 0.703 | 0.279 | 3707.116 | 0.875 | 0.485 |
| | | *NaBott* | 0.622 | 0.062 | 14135.565 | 0.984 | 0.537 |
| | ll1000 | *N1* | 0.683 | 0.175 | 902.776 | 0.926 | 0.513 |
| | | *T* | 0.777 | 0.216 | 3153.159 | 0.903 | 0.522 |
| | | *NaBott* | 0.769 | 0.027 | 10732.898 | 0.984 | 0.561 |
| nc50 | ll200 | *N1* | 0.565 | 0.218 | 980.206 | 0.894 | 0.529 |
| | | *T* | 0.524 | 0.470 | 4187.524 | 0.810 | 0.503 |
| | | *NaBott* | 0.559 | 0.070 | 17791.336 | 0.983 | 0.531 |
| | ll1000 | *N1* | 0.781 | 0.118 | 740.648 | 0.959 | 0.493 |
| | | *T* | 0.911 | 0.175 | 2353.816 | 0.934 | 0.547 |
| | | *NaBott* | 0.800 | 0.041 | 10184.577 | 0.987 | 0.545 |

| | | Coverage 2x | | | | | |
|---|---|---|---|---|---|---|---|
| | | Parameter | $R^2$ | Bias | RMSE | Factor2 | Coverage50% |
| nc10 | ll200 | *N1* | 0.528 | 0.213 | 1024.312 | 0.893 | 0.491 |
| | | *T* | 0.632 | 0.381 | 4088.446 | 0.822 | 0.464 |
| | | *NaBott* | 0.562 | 0.073 | 15641.259 | 0.969 | 0.533 |
| | ll1000 | *N1* | 0.765 | 0.126 | 749.295 | 0.954 | 0.506 |
| | | *T* | 0.846 | 0.159 | 2152.658 | 0.941 | 0.510 |
| | | *NaBott* | 0.780 | 0.019 | 9889.567 | 0.991 | 0.599 |
| nc20 | ll200 | *N1* | 0.630 | 0.187 | 931.103 | 0.929 | 0.469 |
| | | *T* | 0.733 | 0.277 | 3697.900 | 0.872 | 0.492 |
| | | *NaBott* | 0.719 | 0.043 | 13001.665 | 0.987 | 0.559 |
| | ll1000 | *N1* | 0.771 | 0.106 | 709.390 | 0.964 | 0.531 |
| | | *T* | 0.869 | 0.135 | 2089.638 | 0.947 | 0.567 |
| | | *NaBott* | 0.831 | 0.030 | 9436.385 | 0.988 | 0.601 |
| nc50 | ll200 | *N1* | 0.606 | 0.202 | 950.933 | 0.915 | 0.542 |
| | | *T* | 0.624 | 0.318 | 3898.129 | 0.856 | 0.513 |
| | | *NaBott* | 0.531 | 0.064 | 16497.826 | 0.979 | 0.526 |
| | ll1000 | *N1* | 0.740 | 0.152 | 823.798 | 0.940 | 0.463 |
| | | *T* | 0.772 | 0.189 | 2941.760 | 0.923 | 0.471 |
| | | *NaBott* | 0.822 | 0.022 | 10573.725 | 0.992 | 0.537 |

| | | Coverage 5x | | | | | |
|---|---|---|---|---|---|---|---|
| | | Parameter | $R^2$ | Bias | RMSE | Factor2 | Coverage50% |
| nc10 | ll200 | *N1* | 0.670 | 0.209 | 975.256 | 0.905 | 0.521 |
| | | *T* | 0.688 | 0.298 | 3861.590 | 0.842 | 0.486 |
| | | *NaBott* | 0.582 | 0.073 | 15124.538 | 0.976 | 0.533 |
| | ll1000 | *N1* | 0.808 | 0.101 | 688.596 | 0.959 | 0.514 |
| | | *T* | 0.892 | 0.135 | 2047.077 | 0.942 | 0.520 |
| | | *NaBott* | 0.801 | 0.026 | 9778.200 | 0.991 | 0.602 |
| nc20 | ll200 | *N1* | 0.611 | 0.200 | 964.997 | 0.915 | 0.472 |
| | | *T* | 0.694 | 0.276 | 3786.642 | 0.872 | 0.505 |
| | | *NaBott* | 0.596 | 0.053 | 15257.369 | 0.983 | 0.493 |
| | ll1000 | *N1* | 0.821 | 0.084 | 626.388 | 0.973 | 0.532 |
| | | *T* | 0.907 | 0.109 | 1935.173 | 0.958 | 0.546 |
| | | *NaBott* | 0.818 | 0.021 | 9603.793 | 0.988 | 0.625 |
| nc50 | ll200 | *N1* | 0.643 | 0.187 | 918.610 | 0.931 | 0.487 |
| | | *T* | 0.668 | 0.282 | 3617.178 | 0.878 | 0.500 |
| | | *NaBott* | 0.681 | 0.040 | 12577.801 | 0.992 | 0.562 |
| | ll1000 | *N1* | 0.805 | 0.098 | 702.097 | 0.963 | 0.509 |
| | | *T* | 0.878 | 0.140 | 1981.857 | 0.939 | 0.503 |
| | | *NaBott* | 0.832 | 0.010 | 10151.471 | 0.994 | 0.566 |

| | | Coverage 30x | | | | | |
|---|---|---|---|---|---|---|---|
| | | Parameter | $R^2$ | Bias | RMSE | Factor2 | Coverage50% |
| nc10 | ll200 | *N1* | 0.709 | 0.177 | 866.735 | 0.924 | 0.540 |
| | | *T* | 0.717 | 0.260 | 3334.580 | 0.887 | 0.536 |
| | | *NaBott* | 0.709 | 0.048 | 12218.474 | 0.984 | 0.542 |
| | ll1000 | *N1* | 0.801 | 0.072 | 613.803 | 0.974 | 0.572 |
| | | *T* | 0.929 | 0.099 | 1884.530 | 0.961 | 0.584 |
| | | *NaBott* | 0.793 | 0.018 | 9409.010 | 0.995 | 0.661 |
| nc20 | ll200 | *N1* | 0.676 | 0.162 | 847.922 | 0.936 | 0.525 |
| | | *T* | 0.745 | 0.257 | 3329.713 | 0.894 | 0.514 |
| | | *NaBott* | 0.688 | 0.043 | 12887.452 | 0.990 | 0.545 |
| | ll1000 | *N1* | 0.815 | 0.083 | 656.112 | 0.969 | 0.522 |
| | | *T* | 0.909 | 0.096 | 1783.618 | 0.958 | 0.555 |
| | | *NaBott* | 0.870 | 0.015 | 8601.397 | 0.992 | 0.640 |
| nc50 | ll200 | *N1* | 0.732 | 0.139 | 817.584 | 0.948 | 0.521 |
| | | *T* | 0.710 | 0.215 | 3320.334 | 0.894 | 0.534 |
| | | *NaBott* | 0.690 | 0.037 | 13136.178 | 0.984 | 0.546 |
| | ll1000 | *N1* | 0.793 | 0.097 | 644.400 | 0.968 | 0.536 |
| | | *T* | 0.877 | 0.132 | 1907.834 | 0.956 | 0.546 |
| | | *NaBott* | 0.774 | 0.015 | 9593.672 | 0.994 | 0.617 |

**Supplementary Table 4. Accuracy of the estimated parameters of the Bottleneck model assessed by 1,000 pods.** Combinations of experimental parameters considering 5,000 loci.

| | | Coverage 1x | | | | | |
|---|---|---|---|---|---|---|---|
| | | **Parameter** | **$R^2$** | **Bias** | **RMSE** | **Factor2** | **Coverage50%** |
| nc10 | ll200 | *N1* | 0.689 | 0.200 | 886.946 | 0.922 | 0.539 |
| | | *T* | 0.759 | 0.265 | 3207.655 | 0.887 | 0.516 |
| | | *NaBott* | 0.790 | 0.034 | 10771.029 | 0.990 | 0.562 |
| | ll1000 | *N1* | 0.799 | 0.112 | 708.054 | 0.965 | 0.556 |
| | | *T* | 0.882 | 0.136 | 1873.723 | 0.949 | 0.567 |
| | | *NaBott* | 0.860 | 0.016 | 7959.279 | 0.991 | 0.633 |
| nc20 | ll200 | *N1* | 0.723 | 0.135 | 809.505 | 0.948 | 0.501 |
| | | *T* | 0.761 | 0.198 | 3042.307 | 0.911 | 0.517 |
| | | *NaBott* | 0.722 | 0.030 | 11906.801 | 0.988 | 0.578 |
| | ll1000 | *N1* | 0.757 | 0.132 | 770.193 | 0.946 | 0.535 |
| | | *T* | 0.854 | 0.161 | 2400.323 | 0.927 | 0.532 |
| | | *NaBott* | 0.843 | 0.033 | 8244.129 | 0.989 | 0.633 |
| nc50 | ll200 | *N1* | 0.456 | 0.232 | 1017.675 | 0.887 | 0.515 |
| | | *T* | 0.534 | 0.375 | 4169.887 | 0.830 | 0.523 |
| | | *NaBott* | 0.424 | 0.071 | 15794.643 | 0.992 | 0.551 |
| | ll1000 | *N1* | 0.778 | 0.095 | 690.049 | 0.966 | 0.543 |
| | | *T* | 0.875 | 0.120 | 2090.666 | 0.948 | 0.553 |
| | | *NaBott* | 0.872 | 0.014 | 8264.823 | 0.993 | 0.577 |

| | | Coverage 2x | | | | | |
|---|---|---|---|---|---|---|---|
| | | **Parameter** | **$R^2$** | **Bias** | **RMSE** | **Factor2** | **Coverage50%** |
| nc10 | ll200 | *N1* | 0.682 | 0.185 | 892.240 | 0.929 | 0.487 |
| | | *T* | 0.771 | 0.252 | 3204.015 | 0.888 | 0.501 |
| | | *NaBott* | 0.779 | 0.031 | 10773.951 | 0.989 | 0.533 |
| | ll1000 | *N1* | 0.799 | 0.082 | 640.848 | 0.964 | 0.541 |
| | | *T* | 0.895 | 0.098 | 1517.488 | 0.956 | 0.565 |
| | | *NaBott* | 0.900 | 0.011 | 7402.099 | 0.995 | 0.663 |
| nc20 | ll200 | *N1* | 0.717 | 0.157 | 836.943 | 0.940 | 0.524 |
| | | *T* | 0.756 | 0.198 | 2866.225 | 0.914 | 0.525 |
| | | *NaBott* | 0.791 | 0.017 | 10213.974 | 0.993 | 0.582 |
| | ll1000 | *N1* | 0.742 | 0.122 | 724.023 | 0.960 | 0.560 |
| | | *T* | 0.882 | 0.147 | 2010.084 | 0.951 | 0.567 |
| | | *NaBott* | 0.867 | 0.021 | 7348.372 | 0.992 | 0.640 |
| nc50 | ll200 | *N1* | 0.629 | 0.247 | 977.828 | 0.909 | 0.502 |
| | | *T* | 0.650 | 0.329 | 3812.149 | 0.861 | 0.490 |
| | | *NaBott* | 0.665 | 0.047 | 11944.613 | 0.989 | 0.580 |
| | ll1000 | *N1* | 0.728 | 0.151 | 811.795 | 0.942 | 0.534 |
| | | *T* | 0.810 | 0.191 | 2696.072 | 0.926 | 0.531 |
| | | *NaBott* | 0.889 | 0.012 | 7586.827 | 0.996 | 0.582 |

| Coverage 5x | | | | | |
|---|---|---|---|---|---|
| **Parameter** | **$R^2$** | **Bias** | **RMSE** | **Factor2** | **Coverage50%** |

| | | | | | | | |
|---|---|---|---|---|---|---|---|
| nc10 | ll200 | *N1* | 0.747 | 0.103 | 774.891 | 0.959 | 0.510 |
| | | *T* | 0.833 | 0.164 | 2884.581 | 0.928 | 0.508 |
| | | *NaBott* | 0.779 | 0.035 | 11222.058 | 0.987 | 0.554 |
| | ll1000 | *N1* | 0.767 | 0.149 | 746.970 | 0.951 | 0.580 |
| | | *T* | 0.847 | 0.165 | 1962.224 | 0.934 | 0.589 |
| | | *NaBott* | 0.905 | 0.016 | 6977.608 | 0.994 | 0.700 |
| nc20 | ll200 | *N1* | 0.746 | 0.163 | 818.798 | 0.943 | 0.526 |
| | | *T* | 0.759 | 0.220 | 2789.918 | 0.908 | 0.504 |
| | | *NaBott* | 0.794 | 0.026 | 10171.610 | 0.988 | 0.591 |
| | ll1000 | *N1* | 0.767 | 0.107 | 688.589 | 0.953 | 0.575 |
| | | *T* | 0.877 | 0.121 | 1809.353 | 0.949 | 0.569 |
| | | *NaBott* | 0.882 | 0.019 | 7305.038 | 0.994 | 0.684 |
| nc50 | ll200 | *N1* | 0.643 | 0.179 | 901.868 | 0.926 | 0.514 |
| | | *T* | 0.735 | 0.244 | 3610.010 | 0.888 | 0.500 |
| | | *NaBott* | 0.730 | 0.047 | 11336.572 | 0.985 | 0.548 |
| | ll1000 | *N1* | 0.826 | 0.079 | 584.968 | 0.973 | 0.538 |
| | | *T* | 0.938 | 0.089 | 1359.596 | 0.965 | 0.553 |
| | | *NaBott* | 0.890 | 0.020 | 7389.880 | 0.988 | 0.666 |

| | | **Coverage 30x** | | | | | |
|---|---|---|---|---|---|---|---|
| | | **Parameter** | **$R^2$** | **Bias** | **RMSE** | **Factor2** | **Coverage50%** |
| nc10 | ll200 | *N1* | 0.698 | 0.151 | 809.926 | 0.942 | 0.510 |
| | | *T* | 0.801 | 0.191 | 2776.364 | 0.921 | 0.520 |
| | | *NaBott* | 0.782 | 0.022 | 9809.908 | 0.994 | 0.571 |
| | ll1000 | *N1* | 0.842 | 0.073 | 583.787 | 0.972 | 0.571 |
| | | *T* | 0.951 | 0.085 | 1212.466 | 0.968 | 0.600 |
| | | *NaBott* | 0.895 | 0.023 | 6657.717 | 0.993 | 0.710 |
| nc20 | ll200 | *N1* | 0.751 | 0.153 | 832.845 | 0.943 | 0.518 |
| | | *T* | 0.857 | 0.205 | 2786.045 | 0.913 | 0.531 |
| | | *NaBott* | 0.796 | 0.034 | 9999.243 | 0.990 | 0.572 |
| | ll1000 | *N1* | 0.812 | 0.052 | 549.215 | 0.976 | 0.593 |
| | | *T* | 0.945 | 0.066 | 1167.277 | 0.971 | 0.614 |
| | | *NaBott* | 0.892 | 0.011 | 7109.018 | 0.994 | 0.691 |
| nc50 | ll200 | *N1* | 0.757 | 0.138 | 783.094 | 0.949 | 0.535 |
| | | *T* | 0.818 | 0.202 | 2733.306 | 0.916 | 0.526 |
| | | *NaBott* | 0.811 | 0.031 | 9703.053 | 0.994 | 0.594 |
| | ll1000 | *N1* | 0.871 | 0.068 | 537.977 | 0.979 | 0.578 |
| | | *T* | 0.926 | 0.085 | 1259.342 | 0.973 | 0.578 |
| | | *NaBott* | 0.867 | 0.026 | 8417.561 | 0.986 | 0.696 |

**Supplementary Table 5. Accuracy of the estimated parameters of the Exponential Growth model assessed by 1,000 pods.** Combinations of experimental parameters considering 1,000 loci.

| | | Coverage 1x | | | | | |
|---|---|---|---|---|---|---|---|
| | | Parameter | $R^2$ | Bias | RMSE | Factor2 | Coverage50% |
| nc10 | ll200 | *N1* | 0.025 | 0.156 | 21116.612 | 0.927 | 0.510 |
| | | *T* | 0.469 | 0.935 | 3795.697 | 0.785 | 0.559 |
| | | *NaExp* | 0.238 | 0.251 | 1112.305 | 0.856 | 0.560 |
| | ll1000 | *N1* | 0.063 | 0.152 | 21461.613 | 0.928 | 0.466 |
| | | *T* | 0.733 | 0.307 | 3254.462 | 0.872 | 0.515 |
| | | *NaExp* | 0.448 | 0.276 | 1005.609 | 0.860 | 0.512 |
| nc20 | ll200 | *N1* | 0.064 | 0.124 | 21139.963 | 0.935 | 0.511 |
| | | *T* | 0.715 | 0.442 | 3284.099 | 0.859 | 0.550 |
| | | *NaExp* | 0.322 | 0.263 | 1135.775 | 0.855 | 0.521 |
| | ll1000 | *N1* | 0.237 | 0.107 | 21450.154 | 0.946 | 0.466 |
| | | *T* | 0.804 | 0.313 | 3074.880 | 0.889 | 0.504 |
| | | *NaExp* | 0.507 | 0.194 | 1019.320 | 0.875 | 0.496 |
| nc50 | ll200 | *N1* | 0.244 | 0.107 | 19227.428 | 0.970 | 0.501 |
| | | *T* | 0.646 | 0.268 | 3729.226 | 0.835 | 0.558 |
| | | *NaExp* | 0.137 | 0.326 | 1189.785 | 0.812 | 0.514 |
| | ll1000 | *N1* | 0.182 | 0.125 | 21122.796 | 0.952 | 0.497 |
| | | *T* | 0.799 | 0.260 | 2944.336 | 0.896 | 0.475 |
| | | *NaExp* | 0.298 | 0.280 | 1154.632 | 0.828 | 0.488 |

| | | Coverage 2x | | | | | |
|---|---|---|---|---|---|---|---|
| | | Parameter | $R^2$ | Bias | RMSE | Factor2 | Coverage50% |
| nc10 | ll200 | *N1* | 0.047 | 0.139 | 21084.219 | 0.926 | 0.506 |
| | | *T* | 0.643 | 0.483 | 3944.169 | 0.818 | 0.509 |
| | | *NaExp* | 0.312 | 0.280 | 1117.932 | 0.826 | 0.509 |
| | ll1000 | *N1* | 0.124 | 0.148 | 22106.251 | 0.929 | 0.484 |
| | | *T* | 0.804 | 0.303 | 3144.395 | 0.900 | 0.496 |
| | | *NaExp* | 0.497 | 0.210 | 1025.683 | 0.886 | 0.523 |
| nc20 | ll200 | *N1* | 0.079 | 0.140 | 20489.851 | 0.939 | 0.517 |
| | | *T* | 0.743 | 0.323 | 3321.596 | 0.880 | 0.524 |
| | | *NaExp* | 0.295 | 0.300 | 1118.425 | 0.839 | 0.514 |
| | ll1000 | *N1* | 0.126 | 0.130 | 21538.449 | 0.945 | 0.480 |
| | | *T* | 0.771 | 0.279 | 3031.834 | 0.901 | 0.489 |
| | | *NaExp* | 0.329 | 0.284 | 1106.004 | 0.831 | 0.499 |
| nc50 | ll200 | *N1* | 0.102 | 0.119 | 20480.930 | 0.950 | 0.511 |
| | | *T* | 0.587 | 0.504 | 3130.020 | 0.856 | 0.615 |
| | | *NaExp* | 0.123 | 0.337 | 1175.402 | 0.826 | 0.524 |
| | ll1000 | *N1* | 0.221 | 0.093 | 20576.524 | 0.958 | 0.489 |
| | | *T* | 0.793 | 0.192 | 2897.855 | 0.911 | 0.549 |
| | | *NaExp* | 0.236 | 0.315 | 1145.096 | 0.828 | 0.533 |

| | | Coverage 5x | | | | | |
|---|---|---|---|---|---|---|---|
| | | Parameter | $R^2$ | Bias | RMSE | Factor2 | Coverage50% |
| nc10 | ll200 | *N1* | 0.086 | 0.151 | 21265.463 | 0.939 | 0.491 |

| | | | | | | | |
|---|---|---|---|---|---|---|---|
| | | *T* | 0.727 | 0.444 | 3688.859 | 0.823 | 0.494 |
| | | *NaExp* | 0.300 | 0.323 | 1156.656 | 0.816 | 0.488 |
| | ll1000 | *N1* | 0.147 | 0.140 | 21139.987 | 0.925 | 0.516 |
| | | *T* | 0.765 | 0.369 | 2946.913 | 0.879 | 0.511 |
| | | *NaExp* | 0.510 | 0.189 | 999.537 | 0.877 | 0.531 |
| nc20 | ll200 | *N1* | 0.184 | 0.088 | 20573.404 | 0.959 | 0.497 |
| | | *T* | 0.775 | 0.292 | 3324.238 | 0.862 | 0.481 |
| | | *NaExp* | 0.311 | 0.287 | 1156.554 | 0.823 | 0.494 |
| | ll1000 | *N1* | 0.210 | 0.108 | 21900.028 | 0.936 | 0.495 |
| | | *T* | 0.803 | 0.265 | 2960.026 | 0.888 | 0.494 |
| | | *NaExp* | 0.396 | 0.277 | 1142.955 | 0.836 | 0.476 |
| nc50 | ll200 | *N1* | 0.199 | 0.081 | 20230.472 | 0.963 | 0.520 |
| | | *T* | 0.793 | 0.247 | 3004.159 | 0.894 | 0.515 |
| | | *NaExp* | 0.197 | 0.334 | 1219.434 | 0.817 | 0.494 |
| | ll1000 | *N1* | 0.319 | 0.085 | 19967.123 | 0.962 | 0.503 |
| | | *T* | 0.795 | 0.177 | 2998.871 | 0.910 | 0.487 |
| | | *NaExp* | 0.339 | 0.285 | 1214.465 | 0.803 | 0.491 |

| | | **Coverage 30x** | | | | | |
|---|---|---|---|---|---|---|---|
| | | **Parameter** | **$R^2$** | **Bias** | **RMSE** | **Factor2** | **Coverage50%** |
| nc10 | ll200 | *N1* | 0.091 | 0.129 | 21239.587 | 0.940 | 0.501 |
| | | *T* | 0.710 | 0.332 | 3222.802 | 0.873 | 0.540 |
| | | *NaExp* | 0.298 | 0.307 | 1155.671 | 0.834 | 0.494 |
| | ll1000 | *N1* | 0.136 | 0.117 | 21817.818 | 0.938 | 0.481 |
| | | *T* | 0.782 | 0.306 | 2973.979 | 0.891 | 0.503 |
| | | *NaExp* | 0.441 | 0.261 | 1067.447 | 0.847 | 0.470 |
| nc20 | ll200 | *N1* | 0.141 | 0.110 | 20669.920 | 0.959 | 0.491 |
| | | *T* | 0.757 | 0.323 | 3062.532 | 0.880 | 0.517 |
| | | *NaExp* | 0.257 | 0.288 | 1179.409 | 0.833 | 0.508 |
| | ll1000 | *N1* | 0.176 | 0.124 | 21391.027 | 0.947 | 0.493 |
| | | *T* | 0.789 | 0.166 | 2961.960 | 0.914 | 0.503 |
| | | *NaExp* | 0.332 | 0.304 | 1182.575 | 0.829 | 0.502 |
| nc50 | ll200 | *N1* | 0.300 | 0.103 | 21308.067 | 0.951 | 0.488 |
| | | *T* | 0.805 | 0.161 | 3066.184 | 0.907 | 0.491 |
| | | *NaExp* | 0.270 | 0.311 | 1262.277 | 0.792 | 0.489 |
| | ll1000 | *N1* | 0.272 | 0.090 | 20182.885 | 0.962 | 0.514 |
| | | *T* | 0.779 | 0.142 | 2911.160 | 0.917 | 0.505 |
| | | *NaExp* | 0.280 | 0.310 | 1168.234 | 0.822 | 0.538 |

**Supplementary Table 6. Accuracy of the estimated parameters of the Exponential Growth model assessed by 1,000 pods.** Combinations of experimental parameters considering 5,000 loci.

| | | Coverage 1x | | | | | |
|---|---|---|---|---|---|---|---|
| | | **Parameter** | **$R^2$** | **Bias** | **RMSE** | **Factor2** | **Coverage50%** |
| nc10 | ll200 | *N1* | 0.069 | 0.127 | 21227.819 | 0.934 | 0.505 |
| | | *T* | 0.743 | 0.499 | 3390.393 | 0.843 | 0.524 |
| | | *NaExp* | 0.399 | 0.271 | 1075.252 | 0.857 | 0.477 |
| | ll1000 | *N1* | 0.085 | 0.140 | 21437.573 | 0.936 | 0.506 |
| | | *T* | 0.816 | 0.251 | 2908.127 | 0.901 | 0.520 |
| | | *NaExp* | 0.577 | 0.210 | 911.310 | 0.888 | 0.537 |
| nc20 | ll200 | *N1* | 0.193 | 0.108 | 18917.859 | 0.970 | 0.550 |
| | | *T* | 0.695 | 0.209 | 2865.130 | 0.915 | 0.581 |
| | | *NaExp* | 0.241 | 0.264 | 1038.725 | 0.864 | 0.549 |
| | ll1000 | *N1* | 0.226 | 0.126 | 20288.809 | 0.944 | 0.508 |
| | | *T* | 0.817 | 0.366 | 2940.792 | 0.876 | 0.520 |
| | | *NaExp* | 0.573 | 0.164 | 971.513 | 0.878 | 0.509 |
| nc50 | ll200 | *N1* | 0.187 | 0.107 | 19408.133 | 0.964 | 0.525 |
| | | *T* | 0.677 | 0.346 | 3207.315 | 0.859 | 0.558 |
| | | *NaExp* | 0.193 | 0.301 | 1171.286 | 0.825 | 0.530 |
| | ll1000 | *N1* | 0.318 | 0.100 | 17965.959 | 0.967 | 0.504 |
| | | *T* | 0.812 | 0.254 | 2823.050 | 0.889 | 0.520 |
| | | *NaExp* | 0.323 | 0.306 | 1105.199 | 0.851 | 0.512 |

| | | Coverage 2x | | | | | |
|---|---|---|---|---|---|---|---|
| | | **Parameter** | **$R^2$** | **Bias** | **RMSE** | **Factor2** | **Coverage50%** |
| nc10 | ll200 | *N1* | 0.075 | 0.112 | 21122.708 | 0.943 | 0.500 |
| | | *T* | 0.730 | 0.454 | 3245.578 | 0.869 | 0.520 |
| | | *NaExp* | 0.335 | 0.316 | 1084.128 | 0.846 | 0.505 |
| | ll1000 | *N1* | 0.125 | 0.121 | 21941.546 | 0.946 | 0.482 |
| | | *T* | 0.819 | 0.247 | 2904.739 | 0.889 | 0.483 |
| | | *NaExp* | 0.630 | 0.211 | 938.403 | 0.884 | 0.502 |
| nc20 | ll200 | *N1* | 0.129 | 0.142 | 21867.864 | 0.926 | 0.496 |
| | | *T* | 0.785 | 0.307 | 2910.053 | 0.889 | 0.488 |
| | | *NaExp* | 0.323 | 0.288 | 1125.399 | 0.839 | 0.482 |
| | ll1000 | *N1* | 0.454 | 0.083 | 19287.059 | 0.953 | 0.487 |
| | | *T* | 0.854 | 0.091 | 2736.813 | 0.939 | 0.487 |
| | | *NaExp* | 0.685 | 0.154 | 873.470 | 0.904 | 0.512 |
| nc50 | ll200 | *N1* | 0.161 | 0.106 | 19833.435 | 0.959 | 0.521 |
| | | *T* | 0.657 | 0.424 | 2949.957 | 0.885 | 0.608 |
| | | *NaExp* | 0.183 | 0.322 | 1163.026 | 0.834 | 0.532 |
| | ll1000 | *N1* | 0.542 | 0.066 | 17767.134 | 0.967 | 0.488 |
| | | *T* | 0.836 | 0.115 | 2823.295 | 0.939 | 0.485 |
| | | *NaExp* | 0.465 | 0.255 | 1104.797 | 0.845 | 0.464 |

| | | Coverage 5x | | | | | |
|---|---|---|---|---|---|---|---|
| | | **Parameter** | **$R^2$** | **Bias** | **RMSE** | **Factor2** | **Coverage50%** |
| nc10 | ll200 | *N1* | 0.128 | 0.139 | 21602.752 | 0.926 | 0.489 |

| | | | | | | | |
|---|---|---|---|---|---|---|---|
| | | *T* | 0.764 | 0.336 | 3208.567 | 0.872 | 0.511 |
| | | *NaExp* | 0.371 | 0.306 | 1123.043 | 0.833 | 0.492 |
| | ll1000 | *N1* | 0.213 | 0.131 | 22100.502 | 0.928 | 0.500 |
| | | *T* | 0.844 | 0.190 | 2744.433 | 0.895 | 0.494 |
| | | *NaExp* | 0.651 | 0.168 | 894.740 | 0.893 | 0.505 |
| nc20 | ll200 | *N1* | 0.140 | 0.122 | 20460.829 | 0.936 | 0.519 |
| | | *T* | 0.789 | 0.202 | 2970.885 | 0.913 | 0.508 |
| | | *NaExp* | 0.290 | 0.283 | 1131.855 | 0.851 | 0.521 |
| | ll1000 | *N1* | 0.391 | 0.082 | 20204.232 | 0.945 | 0.483 |
| | | *T* | 0.829 | 0.091 | 2752.506 | 0.940 | 0.468 |
| | | *NaExp* | 0.616 | 0.203 | 936.682 | 0.880 | 0.493 |
| nc50 | ll200 | *N1* | 0.357 | 0.085 | 20097.029 | 0.954 | 0.476 |
| | | *T* | 0.825 | 0.137 | 2872.786 | 0.916 | 0.513 |
| | | *NaExp* | 0.309 | 0.278 | 1137.105 | 0.839 | 0.517 |
| | ll1000 | *N1* | 0.488 | 0.060 | 18362.734 | 0.965 | 0.486 |
| | | *T* | 0.835 | 0.118 | 2823.749 | 0.942 | 0.486 |
| | | *NaExp* | 0.506 | 0.266 | 1055.913 | 0.827 | 0.509 |

| | | **Coverage 30x** | | | | | |
|---|---|---|---|---|---|---|---|
| | | **Parameter** | **$R^2$** | **Bias** | **RMSE** | **Factor2** | **Coverage50%** |
| nc10 | ll200 | *N1* | 0.102 | 0.120 | 21027.171 | 0.954 | 0.490 |
| | | *T* | 0.770 | 0.298 | 3008.685 | 0.893 | 0.525 |
| | | *NaExp* | 0.315 | 0.269 | 1113.259 | 0.855 | 0.517 |
| | ll1000 | *N1* | 0.184 | 0.123 | 21545.419 | 0.934 | 0.507 |
| | | *T* | 0.881 | 0.155 | 2418.797 | 0.935 | 0.531 |
| | | *NaExp* | 0.653 | 0.133 | 819.388 | 0.914 | 0.511 |
| nc20 | ll200 | *N1* | 0.148 | 0.135 | 20712.768 | 0.939 | 0.506 |
| | | *T* | 0.815 | 0.213 | 2844.496 | 0.904 | 0.514 |
| | | *NaExp* | 0.273 | 0.287 | 1142.299 | 0.843 | 0.522 |
| | ll1000 | *N1* | 0.212 | 0.119 | 21445.524 | 0.936 | 0.497 |
| | | *T* | 0.820 | 0.144 | 2709.066 | 0.927 | 0.510 |
| | | *NaExp* | 0.490 | 0.249 | 1025.747 | 0.851 | 0.495 |
| nc50 | ll200 | *N1* | 0.304 | 0.092 | 21360.327 | 0.948 | 0.461 |
| | | *T* | 0.822 | 0.157 | 2926.798 | 0.912 | 0.490 |
| | | *NaExp* | 0.287 | 0.322 | 1248.717 | 0.811 | 0.501 |
| | ll1000 | *N1* | 0.294 | 0.069 | 20772.366 | 0.958 | 0.493 |
| | | *T* | 0.821 | 0.116 | 2823.599 | 0.942 | 0.513 |
| | | *NaExp* | 0.378 | 0.255 | 1097.335 | 0.857 | 0.511 |

**Supplementary Table 7. Accuracy of the estimated parameters of the Divergence model assessed by 1,000 pods.** Combinations of experimental parameters considering 1,000 loci.

| | | Coverage 1x | | | | | |
|---|---|---|---|---|---|---|---|
| | | Parameter | R2 | Bias | RMSE | Factor2 | Coverage50% |
| nc10 | ll200 | *N1* | 0.654 | 0.161 | 9155.313 | 0.904 | 0.514 |
| | | *N2* | 0.726 | 0.132 | 8607.355 | 0.932 | 0.499 |
| | | *Nanc* | 0.953 | 0.029 | 2190.376 | 0.985 | 0.542 |
| | | *Tsep* | 0.874 | 0.150 | 2342.224 | 0.930 | 0.554 |
| | ll1000 | *N1* | 0.774 | 0.110 | 6909.512 | 0.962 | 0.594 |
| | | *N2* | 0.783 | 0.091 | 6874.929 | 0.967 | 0.550 |
| | | *Nanc* | 0.982 | 0.011 | 1368.092 | 0.998 | 0.653 |
| | | *Tsep* | 0.888 | 0.079 | 1569.978 | 0.972 | 0.629 |
| nc20 | ll200 | *N1* | 0.750 | 0.120 | 8345.493 | 0.942 | 0.502 |
| | | *N2* | 0.789 | 0.123 | 8075.368 | 0.947 | 0.516 |
| | | *Nanc* | 0.987 | 0.028 | 2400.162 | 0.984 | 0.539 |
| | | *Tsep* | 0.892 | 0.070 | 1940.873 | 0.972 | 0.564 |
| | ll1000 | *N1* | 0.819 | 0.082 | 6234.466 | 0.979 | 0.592 |
| | | *N2* | 0.822 | 0.077 | 6398.797 | 0.969 | 0.573 |
| | | *Nanc* | 0.980 | 0.028 | 1451.580 | 0.993 | 0.627 |
| | | *Tsep* | 0.900 | 0.065 | 1414.270 | 0.977 | 0.659 |
| nc50 | ll200 | *N1* | 0.748 | 0.163 | 9159.231 | 0.908 | 0.502 |
| | | *N2* | 0.677 | 0.252 | 10046.658 | 0.869 | 0.493 |
| | | *Nanc* | 0.935 | 0.076 | 4559.102 | 0.942 | 0.495 |
| | | *Tsep* | 0.797 | 0.154 | 3274.603 | 0.918 | 0.477 |
| | ll1000 | *N1* | 0.765 | 0.146 | 7193.454 | 0.946 | 0.531 |
| | | *N2* | 0.755 | 0.118 | 7284.723 | 0.952 | 0.535 |
| | | *Nanc* | 0.949 | 0.033 | 2430.447 | 0.983 | 0.522 |
| | | *Tsep* | 0.816 | 0.168 | 2378.697 | 0.937 | 0.608 |

| | | Coverage 2x | | | | | |
|---|---|---|---|---|---|---|---|
| | | Parameter | R2 | Bias | RMSE | Factor2 | Coverage50% |
| nc10 | ll200 | *N1* | 0.771 | 0.139 | 8006.414 | 0.950 | 0.537 |
| | | *N2* | 0.722 | 0.164 | 8227.542 | 0.943 | 0.504 |
| | | *Nanc* | 0.970 | 0.024 | 2102.069 | 0.982 | 0.556 |
| | | *Tsep* | 0.921 | 0.079 | 1842.791 | 0.970 | 0.551 |
| | ll1000 | *N1* | 0.848 | 0.077 | 6051.145 | 0.974 | 0.586 |
| | | *N2* | 0.792 | 0.076 | 5915.952 | 0.979 | 0.586 |
| | | *Nanc* | 0.978 | 0.015 | 1326.036 | 0.997 | 0.686 |
| | | *Tsep* | 0.938 | 0.050 | 1183.651 | 0.987 | 0.634 |
| nc20 | ll200 | *N1* | 0.692 | 0.181 | 8445.646 | 0.928 | 0.539 |
| | | *N2* | 0.708 | 0.145 | 8163.903 | 0.929 | 0.537 |
| | | *Nanc* | 0.986 | 0.034 | 2648.142 | 0.981 | 0.526 |
| | | *Tsep* | 0.905 | 0.050 | 1770.296 | 0.980 | 0.533 |
| | ll1000 | *N1* | 0.881 | 0.074 | 5561.477 | 0.985 | 0.588 |
| | | *N2* | 0.813 | 0.075 | 6166.650 | 0.975 | 0.568 |
| | | *Nanc* | 0.979 | 0.008 | 1550.086 | 0.996 | 0.651 |
| | | *Tsep* | 0.933 | 0.036 | 1181.370 | 0.994 | 0.661 |

| | | | | | | | |
|---|---|---|---|---|---|---|---|
| nc50 | ll200 | *N1* | 0.793 | 0.061 | 7811.865 | 0.956 | 0.537 |
| | | *N2* | 0.676 | 0.213 | 9380.944 | 0.888 | 0.510 |
| | | *Nanc* | 0.938 | 0.081 | 4182.347 | 0.959 | 0.525 |
| | | *Tsep* | 0.737 | 0.170 | 3203.063 | 0.906 | 0.541 |
| | ll1000 | *N1* | 0.809 | 0.086 | 6216.943 | 0.974 | 0.574 |
| | | *N2* | 0.795 | 0.105 | 6435.257 | 0.971 | 0.543 |
| | | *Nanc* | 0.961 | 0.014 | 2313.210 | 0.992 | 0.586 |
| | | *Tsep* | 0.875 | 0.068 | 1819.484 | 0.980 | 0.641 |

| | | **Coverage 5x** | | | | | |
|---|---|---|---|---|---|---|---|
| | | **Parameter** | **R2** | **Bias** | **RMSE** | **Factor2** | **Coverage50%** |
| nc10 | ll200 | *N1* | 0.737 | 0.142 | 7468.780 | 0.946 | 0.524 |
| | | *N2* | 0.785 | 0.085 | 7241.530 | 0.957 | 0.543 |
| | | *Nanc* | 0.968 | 0.013 | 2432.867 | 0.992 | 0.569 |
| | | *Tsep* | 0.881 | 0.073 | 1695.713 | 0.969 | 0.601 |
| | ll1000 | *N1* | 0.867 | 0.067 | 5827.342 | 0.984 | 0.583 |
| | | *N2* | 0.822 | 0.058 | 5438.979 | 0.984 | 0.605 |
| | | *Nanc* | 0.976 | 0.002 | 1318.983 | 1.000 | 0.701 |
| | | *Tsep* | 0.936 | 0.023 | 922.973 | 0.996 | 0.734 |
| nc20 | ll200 | *N1* | 0.794 | 0.087 | 7304.609 | 0.961 | 0.516 |
| | | *N2* | 0.801 | 0.107 | 7280.575 | 0.958 | 0.527 |
| | | *Nanc* | 0.950 | 0.036 | 2776.566 | 0.979 | 0.543 |
| | | *Tsep* | 0.922 | 0.044 | 1527.070 | 0.984 | 0.585 |
| | ll1000 | *N1* | 0.850 | 0.061 | 5486.603 | 0.982 | 0.591 |
| | | *N2* | 0.843 | 0.061 | 5345.512 | 0.985 | 0.626 |
| | | *Nanc* | 0.974 | 0.020 | 1469.626 | 0.994 | 0.676 |
| | | *Tsep* | 0.944 | 0.018 | 880.199 | 0.997 | 0.729 |
| nc50 | ll200 | *N1* | 0.761 | 0.092 | 7296.399 | 0.958 | 0.532 |
| | | *N2* | 0.822 | 0.072 | 6637.222 | 0.978 | 0.525 |
| | | *Nanc* | 0.943 | 0.051 | 3306.031 | 0.976 | 0.549 |
| | | *Tsep* | 0.907 | 0.036 | 1731.411 | 0.992 | 0.557 |
| | ll1000 | *N1* | 0.849 | 0.035 | 4603.966 | 0.989 | 0.630 |
| | | *N2* | 0.872 | 0.034 | 4497.882 | 0.993 | 0.626 |
| | | *Nanc* | 0.974 | 0.006 | 1760.389 | 0.998 | 0.680 |
| | | *Tsep* | 0.969 | 0.012 | 857.029 | 0.998 | 0.719 |

| | | **Coverage 30x** | | | | | |
|---|---|---|---|---|---|---|---|
| | | **Parameter** | **R2** | **Bias** | **RMSE** | **Factor2** | **Coverage50%** |
| nc10 | ll200 | *N1* | 0.739 | 0.158 | 7842.213 | 0.945 | 0.554 |
| | | *N2* | 0.765 | 0.096 | 7788.829 | 0.954 | 0.543 |
| | | *Nanc* | 0.970 | 0.025 | 2148.120 | 0.993 | 0.568 |
| | | *Tsep* | 0.911 | 0.057 | 1686.612 | 0.972 | 0.578 |
| | ll1000 | *N1* | 0.835 | 0.039 | 5170.745 | 0.990 | 0.617 |
| | | *N2* | 0.852 | 0.059 | 5825.852 | 0.982 | 0.601 |
| | | *Nanc* | 0.975 | 0.004 | 1339.137 | 0.998 | 0.709 |
| | | *Tsep* | 0.939 | 0.030 | 881.715 | 0.996 | 0.742 |
| nc20 | ll200 | *N1* | 0.812 | 0.065 | 7073.019 | 0.972 | 0.543 |
| | | *N2* | 0.775 | 0.113 | 7329.004 | 0.958 | 0.543 |

| | | | | | | | |
|---|---|---|---|---|---|---|---|
| | | *Nanc* | 0.976 | 0.035 | 2514.890 | 0.988 | 0.553 |
| | | *Tsep* | 0.922 | 0.050 | 1340.213 | 0.986 | 0.612 |
| | ll1000 | *N1* | 0.843 | 0.044 | 4985.146 | 0.989 | 0.610 |
| | | *N2* | 0.832 | 0.047 | 5365.946 | 0.988 | 0.622 |
| | | *Nanc* | 0.984 | 0.007 | 1505.727 | 0.998 | 0.693 |
| | | *Tsep* | 0.962 | 0.014 | 769.479 | 1.000 | 0.790 |
| nc50 | ll200 | *N1* | 0.813 | 0.062 | 6301.582 | 0.976 | 0.549 |
| | | *N2* | 0.827 | 0.064 | 6040.040 | 0.975 | 0.584 |
| | | *Nanc* | 0.954 | 0.018 | 2938.266 | 0.990 | 0.558 |
| | | *Tsep* | 0.922 | 0.027 | 1359.340 | 0.995 | 0.630 |
| | ll1000 | *N1* | 0.873 | 0.066 | 5139.122 | 0.984 | 0.622 |
| | | *N2* | 0.842 | 0.060 | 5303.823 | 0.982 | 0.626 |
| | | *Nanc* | 0.967 | 0.025 | 1981.573 | 0.988 | 0.635 |
| | | *Tsep* | 0.923 | 0.025 | 1207.228 | 0.995 | 0.696 |

**Supplementary Table 8. Accuracy of the estimated parameters of the Divergence model assessed by 1,000 pods.** Combinations of experimental parameters considering 5,000 loci.

| | | Coverage 1x | | | | | |
|---|---|---|---|---|---|---|---|
| | | **Parameter** | **R2** | **Bias** | **RMSE** | **Factor2** | **Coverage50%** |
| nc10 | ll200 | *N1* | 0.807 | 0.071 | 6178.998 | 0.968 | 0.541 |
| | | *N2* | 0.753 | 0.119 | 7303.880 | 0.943 | 0.549 |
| | | *Nanc* | 0.993 | 0.007 | 1287.755 | 0.997 | 0.571 |
| | | *Tsep* | 0.901 | 0.072 | 1762.754 | 0.973 | 0.594 |
| | ll1000 | *N1* | 0.857 | 0.045 | 5122.714 | 0.992 | 0.656 |
| | | *N2* | 0.854 | 0.047 | 4828.987 | 0.991 | 0.621 |
| | | *Nanc* | 0.991 | 0.004 | 826.891 | 1.000 | 0.706 |
| | | *Tsep* | 0.921 | 0.048 | 1071.065 | 0.993 | 0.724 |
| nc20 | ll200 | *N1* | 0.792 | 0.138 | 6912.684 | 0.954 | 0.572 |
| | | *N2* | 0.837 | 0.057 | 5927.945 | 0.980 | 0.551 |
| | | *Nanc* | 0.992 | 0.017 | 1503.813 | 0.993 | 0.626 |
| | | *Tsep* | 0.897 | 0.062 | 1634.868 | 0.979 | 0.591 |
| | ll1000 | *N1* | 0.889 | 0.042 | 4605.538 | 0.989 | 0.650 |
| | | *N2* | 0.847 | 0.048 | 4801.478 | 0.990 | 0.642 |
| | | *Nanc* | 0.993 | 0.006 | 926.783 | 0.998 | 0.691 |
| | | *Tsep* | 0.946 | 0.024 | 781.951 | 0.997 | 0.695 |
| nc50 | ll200 | *N1* | 0.702 | 0.172 | 8230.381 | 0.919 | 0.517 |
| | | *N2* | 0.826 | 0.088 | 6455.792 | 0.953 | 0.564 |
| | | *Nanc* | 0.981 | 0.015 | 2692.224 | 0.977 | 0.541 |
| | | *Tsep* | 0.811 | 0.097 | 2580.004 | 0.941 | 0.541 |
| | ll1000 | *N1* | 0.849 | 0.062 | 4687.746 | 0.983 | 0.617 |
| | | *N2* | 0.877 | 0.062 | 4378.261 | 0.984 | 0.643 |
| | | *Nanc* | 0.977 | 0.009 | 1335.554 | 0.995 | 0.671 |
| | | *Tsep* | 0.911 | 0.060 | 1522.001 | 0.986 | 0.668 |

| | | Coverage 2x | | | | | |
|---|---|---|---|---|---|---|---|
| | | Parameter | R2 | Bias | RMSE | Factor2 | Coverage50% |
| nc10 | ll200 | *N1* | 0.840 | 0.076 | 5928.333 | 0.973 | 0.576 |
| | | *N2* | 0.783 | 0.097 | 6285.699 | 0.965 | 0.582 |
| | | *Nanc* | 0.987 | 0.014 | 1306.460 | 0.997 | 0.603 |
| | | *Tsep* | 0.918 | 0.068 | 1395.310 | 0.980 | 0.647 |
| | ll1000 | *N1* | 0.896 | 0.052 | 4385.383 | 0.981 | 0.646 |
| | | *N2* | 0.890 | 0.049 | 4507.875 | 0.987 | 0.670 |
| | | *Nanc* | 0.987 | 0.001 | 745.340 | 0.999 | 0.766 |
| | | *Tsep* | 0.960 | 0.037 | 716.737 | 0.992 | 0.787 |
| nc20 | ll200 | *N1* | 0.809 | 0.068 | 6364.359 | 0.974 | 0.567 |
| | | *N2* | 0.803 | 0.078 | 6314.701 | 0.971 | 0.572 |
| | | *Nanc* | 0.982 | 0.006 | 1668.263 | 0.997 | 0.582 |
| | | *Tsep* | 0.919 | 0.046 | 1223.175 | 0.989 | 0.653 |
| | ll1000 | *N1* | 0.866 | 0.042 | 4630.029 | 0.987 | 0.670 |
| | | *N2* | 0.884 | 0.050 | 4982.631 | 0.987 | 0.646 |
| | | *Nanc* | 0.989 | 0.008 | 988.891 | 0.997 | 0.703 |
| | | *Tsep* | 0.968 | 0.026 | 766.731 | 0.998 | 0.749 |
| nc50 | ll200 | *N1* | 0.850 | 0.061 | 6320.253 | 0.980 | 0.560 |
| | | *N2* | 0.821 | 0.067 | 6513.861 | 0.979 | 0.546 |
| | | *Nanc* | 0.966 | 0.034 | 2686.499 | 0.978 | 0.539 |
| | | *Tsep* | 0.938 | 0.032 | 1555.236 | 0.994 | 0.603 |
| | ll1000 | *N1* | 0.837 | 0.042 | 4607.084 | 0.988 | 0.642 |
| | | *N2* | 0.854 | 0.046 | 4621.131 | 0.989 | 0.606 |
| | | *Nanc* | 0.976 | 0.027 | 1427.035 | 0.993 | 0.654 |
| | | *Tsep* | 0.927 | 0.035 | 1199.911 | 0.992 | 0.690 |

| | | Coverage 5x | | | | | |
|---|---|---|---|---|---|---|---|
| | | Parameter | R2 | Bias | RMSE | Factor2 | Coverage50% |
| nc10 | ll200 | *N1* | 0.867 | 0.054 | 5591.553 | 0.981 | 0.600 |
| | | *N2* | 0.803 | 0.096 | 6650.584 | 0.954 | 0.577 |
| | | *Nanc* | 0.988 | 0.001 | 1289.186 | 0.999 | 0.648 |
| | | *Tsep* | 0.943 | 0.043 | 1039.058 | 0.989 | 0.682 |
| | ll1000 | *N1* | 0.875 | 0.034 | 4574.602 | 0.993 | 0.639 |
| | | *N2* | 0.910 | 0.029 | 3978.111 | 0.996 | 0.665 |
| | | *Nanc* | 0.991 | 0.002 | 789.572 | 0.998 | 0.742 |
| | | *Tsep* | 0.964 | 0.015 | 569.830 | 1.000 | 0.818 |
| nc20 | ll200 | *N1* | 0.856 | 0.068 | 5841.968 | 0.984 | 0.537 |
| | | *N2* | 0.828 | 0.071 | 5930.085 | 0.979 | 0.552 |
| | | *Nanc* | 0.972 | 0.004 | 1723.490 | 1.000 | 0.592 |
| | | *Tsep* | 0.959 | 0.030 | 981.257 | 0.996 | 0.633 |
| | ll1000 | *N1* | 0.896 | 0.029 | 4100.869 | 0.993 | 0.671 |
| | | *N2* | 0.883 | 0.035 | 4308.574 | 0.990 | 0.642 |
| | | *Nanc* | 0.987 | 0.008 | 988.304 | 0.998 | 0.717 |
| | | *Tsep* | 0.959 | 0.013 | 615.090 | 0.999 | 0.807 |
| nc50 | ll200 | *N1* | 0.857 | 0.054 | 5796.990 | 0.979 | 0.595 |
| | | *N2* | 0.859 | 0.059 | 5646.839 | 0.984 | 0.566 |

| | | | | | | | |
|---|---|---|---|---|---|---|---|
| | | *Nanc* | 0.973 | 0.021 | 2170.771 | 0.993 | 0.584 |
| | | *Tsep* | 0.951 | 0.017 | 1162.794 | 0.999 | 0.655 |
| | ll1000 | *N1* | 0.881 | 0.059 | 4261.002 | 0.984 | 0.653 |
| | | *N2* | 0.890 | 0.056 | 4507.761 | 0.983 | 0.641 |
| | | *Nanc* | 0.977 | 0.007 | 1280.497 | 0.994 | 0.690 |
| | | *Tsep* | 0.956 | 0.023 | 883.002 | 0.997 | 0.729 |

| | | **Coverage 30x** | | | | | |
|---|---|---|---|---|---|---|---|
| | | **Parameter** | **R2** | **Bias** | **RMSE** | **Factor2** | **Coverage50%** |
| nc10 | ll200 | *N1* | 0.825 | 0.090 | 6416.640 | 0.966 | 0.610 |
| | | *N2* | 0.857 | 0.056 | 5657.955 | 0.982 | 0.577 |
| | | *Nanc* | 0.981 | 0.004 | 1416.384 | 0.998 | 0.620 |
| | | *Tsep* | 0.951 | 0.039 | 1075.387 | 0.991 | 0.705 |
| | ll1000 | *N1* | 0.873 | 0.045 | 4352.897 | 0.988 | 0.683 |
| | | *N2* | 0.885 | 0.032 | 4208.989 | 0.993 | 0.652 |
| | | *Nanc* | 0.987 | 0.003 | 824.537 | 0.998 | 0.722 |
| | | *Tsep* | 0.963 | 0.029 | 606.403 | 0.997 | 0.837 |
| nc20 | ll200 | *N1* | 0.872 | 0.045 | 5595.820 | 0.987 | 0.561 |
| | | *N2* | 0.862 | 0.063 | 5662.925 | 0.980 | 0.590 |
| | | *Nanc* | 0.977 | 0.004 | 1576.824 | 0.998 | 0.603 |
| | | *Tsep* | 0.945 | 0.022 | 908.414 | 0.994 | 0.714 |
| | ll1000 | *N1* | 0.913 | 0.035 | 3782.402 | 0.995 | 0.668 |
| | | *N2* | 0.908 | 0.034 | 4001.358 | 0.992 | 0.663 |
| | | *Nanc* | 0.991 | 0.004 | 913.430 | 0.998 | 0.736 |
| | | *Tsep* | 0.976 | 0.019 | 536.396 | 0.999 | 0.849 |
| nc50 | ll200 | *N1* | 0.863 | 0.044 | 5264.049 | 0.985 | 0.604 |
| | | *N2* | 0.877 | 0.040 | 5228.546 | 0.990 | 0.601 |
| | | *Nanc* | 0.981 | 0.006 | 1856.134 | 0.997 | 0.617 |
| | | *Tsep* | 0.958 | 0.013 | 980.051 | 0.997 | 0.703 |
| | ll1000 | *N1* | 0.895 | 0.018 | 3594.999 | 0.996 | 0.691 |
| | | *N2* | 0.898 | 0.020 | 3546.578 | 0.998 | 0.681 |
| | | *Nanc* | 0.978 | 0.003 | 1020.853 | 1.000 | 0.734 |
| | | *Tsep* | 0.968 | 0.013 | 582.188 | 1.000 | 0.818 |

**Supplementary Table 9. Accuracy of the estimated parameters of the Divergence with migration model assessed by 1,000 pods.**
Combinations of experimental parameters considering 1,000 loci.

| | | Coverage 1x | | | | | |
|---|---|---|---|---|---|---|---|
| | | Parameter | R2 | Bias | RMSE | Factor2 | Coverage50% |
| nc10 | ll200 | *N1* | 0.500 | 0.461 | 11412.411 | 0.794 | 0.505 |
| | | *N2* | 0.461 | 0.445 | 11673.160 | 0.781 | 0.509 |
| | | *Nanc* | 0.932 | 0.079 | 4415.267 | 0.964 | 0.504 |
| | | *Tsep* | 0.274 | 0.627 | 5309.609 | 0.731 | 0.507 |
| | | *m12* | 0.125 | 7.595 | 40.622 | 0.516 | 0.510 |
| | | *m21* | 0.123 | 3.611 | 40.730 | 0.465 | 0.494 |
| | ll1000 | *N1* | 0.622 | 0.356 | 9862.770 | 0.829 | 0.525 |
| | | *N2* | 0.538 | 0.340 | 10646.689 | 0.826 | 0.513 |
| | | *Nanc* | 0.979 | 0.029 | 2451.871 | 0.973 | 0.577 |
| | | *Tsep* | 0.369 | 0.583 | 4917.960 | 0.748 | 0.538 |
| | | *m12* | 0.184 | 1.410 | 35.288 | 0.562 | 0.534 |
| | | *m21* | 0.132 | 3.127 | 41.252 | 0.483 | 0.502 |
| nc20 | ll200 | *N1* | 0.605 | 0.342 | 10495.142 | 0.826 | 0.484 |
| | | *N2* | 0.576 | 0.256 | 10938.138 | 0.823 | 0.527 |
| | | *Nanc* | 0.955 | 0.062 | 3516.476 | 0.969 | 0.546 |
| | | *Tsep* | 0.355 | 0.695 | 5147.629 | 0.725 | 0.492 |
| | | *m12* | 0.180 | 15.968 | 38.718 | 0.510 | 0.503 |
| | | *m21* | 0.160 | 4.655 | 39.264 | 0.475 | 0.492 |
| | ll1000 | *N1* | 0.622 | 0.249 | 9548.361 | 0.855 | 0.509 |
| | | *N2* | 0.592 | 0.375 | 10354.273 | 0.842 | 0.487 |
| | | *Nanc* | 0.967 | 0.068 | 2019.813 | 0.975 | 0.597 |
| | | *Tsep* | 0.396 | 0.672 | 4684.152 | 0.769 | 0.513 |
| | | *m12* | 0.208 | 4.353 | 38.277 | 0.582 | 0.529 |
| | | *m21* | 0.153 | 2.616 | 37.795 | 0.521 | 0.516 |
| nc50 | ll200 | *N1* | 0.714 | 0.293 | 9933.899 | 0.859 | 0.475 |
| | | *N2* | 0.490 | 0.555 | 12289.071 | 0.743 | 0.488 |
| | | *Nanc* | 0.916 | 0.139 | 5337.672 | 0.927 | 0.488 |
| | | *Tsep* | 0.274 | 0.986 | 5662.100 | 0.685 | 0.477 |
| | | *m12* | 0.187 | 2.639 | 40.058 | 0.485 | 0.471 |
| | | *m21* | 0.165 | 3.192 | 40.944 | 0.477 | 0.482 |
| | ll1000 | *N1* | 0.675 | 0.269 | 9051.272 | 0.853 | 0.535 |
| | | *N2* | 0.639 | 0.234 | 9598.678 | 0.882 | 0.523 |
| | | *Nanc* | 0.966 | 0.066 | 2526.095 | 0.971 | 0.587 |
| | | *Tsep* | 0.355 | 0.669 | 4966.753 | 0.731 | 0.520 |
| | | *m12* | 0.205 | 2.606 | 39.451 | 0.561 | 0.522 |
| | | *m21* | 0.153 | 3.609 | 39.414 | 0.481 | 0.491 |

| | | Coverage 2x | | | | | |
|---|---|---|---|---|---|---|---|
| | | Parameter | R2 | Bias | RMSE | Factor2 | Coverage50% |
| nc10 | ll200 | *N1* | 0.618 | 0.357 | 10659.104 | 0.814 | 0.490 |
| | | *N2* | 0.513 | 0.426 | 11703.306 | 0.793 | 0.518 |

| | | | | | | | |
|---|---|---|---|---|---|---|---|
| | | *Nanc* | 0.946 | 0.088 | 3409.459 | 0.970 | 0.521 |
| | | *Tsep* | 0.329 | 0.762 | 5262.514 | 0.732 | 0.501 |
| | | *m12* | 0.172 | 3.183 | 38.482 | 0.550 | 0.536 |
| | | *m21* | 0.128 | 5.198 | 39.217 | 0.485 | 0.505 |
| | ll1000 | *N1* | 0.647 | 0.314 | 9762.591 | 0.851 | 0.518 |
| | | *N2* | 0.586 | 0.247 | 10268.196 | 0.853 | 0.520 |
| | | *Nanc* | 0.962 | 0.049 | 1987.866 | 0.980 | 0.601 |
| | | *Tsep* | 0.433 | 0.482 | 4938.470 | 0.754 | 0.513 |
| | | *m12* | 0.207 | 1.432 | 40.742 | 0.590 | 0.532 |
| | | *m21* | 0.134 | 4.083 | 38.691 | 0.524 | 0.523 |
| nc20 | ll200 | *N1* | 0.637 | 0.317 | 10269.919 | 0.832 | 0.522 |
| | | *N2* | 0.540 | 0.327 | 11170.528 | 0.803 | 0.511 |
| | | *Nanc* | 0.932 | 0.046 | 3930.110 | 0.969 | 0.519 |
| | | *Tsep* | 0.304 | 0.855 | 5449.880 | 0.698 | 0.480 |
| | | *m12* | 0.176 | 2.745 | 38.719 | 0.504 | 0.485 |
| | | *m21* | 0.141 | 1.577 | 40.170 | 0.511 | 0.516 |
| | ll1000 | *N1* | 0.690 | 0.250 | 9128.129 | 0.861 | 0.507 |
| | | *N2* | 0.562 | 0.291 | 10281.309 | 0.842 | 0.489 |
| | | *Nanc* | 0.956 | 0.056 | 1997.829 | 0.981 | 0.619 |
| | | *Tsep* | 0.402 | 0.587 | 4685.042 | 0.751 | 0.542 |
| | | *m12* | 0.242 | 2.782 | 32.390 | 0.603 | 0.540 |
| | | *m21* | 0.143 | 3.635 | 41.568 | 0.513 | 0.504 |
| nc50 | ll200 | *N1* | 0.752 | 0.248 | 8814.155 | 0.887 | 0.502 |
| | | *N2* | 0.438 | 0.437 | 12448.803 | 0.765 | 0.507 |
| | | *Nanc* | 0.904 | 0.180 | 4931.489 | 0.934 | 0.509 |
| | | *Tsep* | 0.246 | 0.934 | 5691.736 | 0.715 | 0.477 |
| | | *m12* | 0.166 | 15.790 | 42.416 | 0.498 | 0.499 |
| | | *m21* | 0.162 | 6.794 | 38.286 | 0.494 | 0.510 |
| | ll1000 | *N1* | 0.777 | 0.172 | 8194.833 | 0.903 | 0.538 |
| | | *N2* | 0.633 | 0.278 | 9615.627 | 0.870 | 0.493 |
| | | *Nanc* | 0.945 | 0.136 | 2660.518 | 0.960 | 0.599 |
| | | *Tsep* | 0.340 | 0.712 | 5013.077 | 0.759 | 0.490 |
| | | *m12* | 0.234 | 1.685 | 35.069 | 0.596 | 0.521 |
| | | *m21* | 0.175 | 1.799 | 38.409 | 0.571 | 0.548 |

| | | **Coverage 5x** | | | | | |
|---|---|---|---|---|---|---|---|
| | | **Parameter** | **R2** | **Bias** | **RMSE** | **Factor2** | **Coverage50%** |
| nc10 | ll200 | *N1* | 0.608 | 0.305 | 10754.244 | 0.808 | 0.503 |
| | | *N2* | 0.554 | 0.361 | 11440.532 | 0.787 | 0.491 |
| | | *Nanc* | 0.956 | 0.056 | 3325.220 | 0.973 | 0.575 |
| | | *Tsep* | 0.326 | 0.877 | 5322.455 | 0.697 | 0.489 |
| | | *m12* | 0.157 | 2.909 | 36.520 | 0.516 | 0.507 |
| | | *m21* | 0.143 | 2.542 | 39.800 | 0.511 | 0.513 |
| | ll1000 | *N1* | 0.656 | 0.270 | 9010.178 | 0.873 | 0.488 |
| | | *N2* | 0.581 | 0.246 | 9780.536 | 0.890 | 0.516 |
| | | *Nanc* | 0.954 | 0.086 | 2648.879 | 0.971 | 0.592 |
| | | *Tsep* | 0.467 | 0.575 | 4864.997 | 0.749 | 0.487 |
| | | *m12* | 0.230 | 2.382 | 33.928 | 0.604 | 0.535 |

| | | | | | | | |
|---|---|---|---|---|---|---|---|
| | | *m21* | 0.156 | 76.879 | 39.386 | 0.502 | 0.484 |
| nc20 | ll200 | *N1* | 0.719 | 0.238 | 9298.476 | 0.868 | 0.511 |
| | | *N2* | 0.626 | 0.227 | 10084.465 | 0.848 | 0.508 |
| | | *Nanc* | 0.934 | 0.104 | 4007.075 | 0.956 | 0.525 |
| | | *Tsep* | 0.328 | 0.743 | 5295.169 | 0.705 | 0.468 |
| | | *m12* | 0.245 | 3.023 | 32.680 | 0.584 | 0.542 |
| | | *m21* | 0.160 | 2.974 | 40.577 | 0.498 | 0.498 |
| | ll1000 | *N1* | 0.721 | 0.192 | 8568.671 | 0.901 | 0.503 |
| | | *N2* | 0.671 | 0.191 | 9186.787 | 0.888 | 0.498 |
| | | *Nanc* | 0.958 | 0.052 | 2187.812 | 0.976 | 0.595 |
| | | *Tsep* | 0.444 | 0.512 | 4539.489 | 0.785 | 0.552 |
| | | *m12* | 0.228 | 2.535 | 35.677 | 0.596 | 0.511 |
| | | *m21* | 0.183 | 195.083 | 41.262 | 0.533 | 0.526 |
| nc50 | ll200 | *N1* | 0.740 | 0.214 | 8796.621 | 0.891 | 0.499 |
| | | *N2* | 0.729 | 0.159 | 9062.182 | 0.900 | 0.488 |
| | | *Nanc* | 0.912 | 0.135 | 3990.508 | 0.944 | 0.520 |
| | | *Tsep* | 0.281 | 0.956 | 5234.892 | 0.711 | 0.479 |
| | | *m12* | 0.203 | 2.608 | 37.965 | 0.564 | 0.511 |
| | | *m21* | 0.180 | 7.142 | 36.108 | 0.508 | 0.501 |
| | ll1000 | *N1* | 0.781 | 0.163 | 7593.648 | 0.910 | 0.489 |
| | | *N2* | 0.714 | 0.172 | 8653.066 | 0.922 | 0.510 |
| | | *Nanc* | 0.936 | 0.078 | 2855.482 | 0.961 | 0.584 |
| | | *Tsep* | 0.346 | 0.696 | 4810.749 | 0.779 | 0.523 |
| | | *m12* | 0.260 | 2.745 | 33.679 | 0.641 | 0.560 |
| | | *m21* | 0.202 | 2.342 | 35.991 | 0.560 | 0.521 |

| | | Coverage 30x | | | | | |
|---|---|---|---|---|---|---|---|
| | | **Parameter** | **R2** | **Bias** | **RMSE** | **Factor2** | **Coverage50%** |
| nc10 | ll200 | *N1* | 0.597 | 0.368 | 10510.496 | 0.817 | 0.522 |
| | | *N2* | 0.586 | 0.309 | 10604.448 | 0.813 | 0.511 |
| | | *Nanc* | 0.953 | 0.066 | 3713.520 | 0.969 | 0.522 |
| | | *Tsep* | 0.296 | 0.897 | 5190.186 | 0.716 | 0.513 |
| | | *m12* | 0.166 | 2.962 | 38.768 | 0.514 | 0.494 |
| | | *m21* | 0.137 | 14.923 | 41.897 | 0.475 | 0.493 |
| | ll1000 | *N1* | 0.678 | 0.220 | 9135.007 | 0.882 | 0.497 |
| | | *N2* | 0.618 | 0.214 | 9430.574 | 0.869 | 0.527 |
| | | *Nanc* | 0.969 | 0.049 | 1965.322 | 0.977 | 0.630 |
| | | *Tsep* | 0.409 | 0.544 | 4644.812 | 0.783 | 0.500 |
| | | *m12* | 0.222 | 2.115 | 33.930 | 0.617 | 0.543 |
| | | *m21* | 0.157 | 9.580 | 36.890 | 0.529 | 0.531 |
| nc20 | ll200 | *N1* | 0.736 | 0.228 | 8727.574 | 0.884 | 0.512 |
| | | *N2* | 0.573 | 0.289 | 10578.938 | 0.848 | 0.517 |
| | | *Nanc* | 0.923 | 0.114 | 3606.423 | 0.947 | 0.538 |
| | | *Tsep* | 0.263 | 0.950 | 5299.382 | 0.717 | 0.477 |
| | | *m12* | 0.220 | 2.912 | 33.331 | 0.580 | 0.519 |
| | | *m21* | 0.165 | 2.322 | 37.555 | 0.506 | 0.507 |
| | ll1000 | *N1* | 0.700 | 0.202 | 8604.772 | 0.880 | 0.496 |
| | | *N2* | 0.691 | 0.171 | 8584.153 | 0.916 | 0.521 |

| | | | | | | | |
|---|---|---|---|---|---|---|---|
| | | *Nanc* | 0.960 | 0.053 | 2110.011 | 0.983 | 0.652 |
| | | *Tsep* | 0.407 | 0.574 | 4585.663 | 0.760 | 0.536 |
| | | *m12* | 0.235 | 2.053 | 34.106 | 0.619 | 0.530 |
| | | *m21* | 0.181 | 5.190 | 38.455 | 0.531 | 0.494 |
| nc50 | ll200 | *N1* | 0.765 | 0.138 | 8063.151 | 0.912 | 0.482 |
| | | *N2* | 0.664 | 0.182 | 9667.601 | 0.880 | 0.487 |
| | | *Nanc* | 0.897 | 0.147 | 4698.857 | 0.930 | 0.545 |
| | | *Tsep* | 0.276 | 1.000 | 5231.801 | 0.733 | 0.505 |
| | | *m12* | 0.245 | 3.198 | 33.545 | 0.583 | 0.534 |
| | | *m21* | 0.188 | 5.971 | 38.807 | 0.499 | 0.492 |
| | ll1000 | *N1* | 0.794 | 0.151 | 7100.099 | 0.929 | 0.475 |
| | | *N2* | 0.741 | 0.132 | 8051.725 | 0.934 | 0.519 |
| | | *Nanc* | 0.955 | 0.084 | 2756.833 | 0.965 | 0.613 |
| | | *Tsep* | 0.365 | 0.631 | 4760.645 | 0.766 | 0.528 |
| | | *m12* | 0.249 | 2.100 | 31.714 | 0.628 | 0.539 |
| | | *m21* | 0.210 | 2.429 | 34.605 | 0.579 | 0.544 |

**Supplementary Table 10. Accuracy of the estimated parameters of the Divergence with migration model assessed by 1,000 pods.** Combinations of experimental parameters considering 5,000 loci.

| | | Coverage 1x | | | | | |
|---|---|---|---|---|---|---|---|
| | | Parameter | R2 | Bias | RMSE | Factor2 | Coverage50% |
| nc10 | ll200 | *N1* | 0.659 | 0.292 | 9835.659 | 0.849 | 0.534 |
| | | *N2* | 0.489 | 0.383 | 11685.146 | 0.782 | 0.486 |
| | | *Nanc* | 0.973 | 0.041 | 2800.031 | 0.976 | 0.556 |
| | | *Tsep* | 0.402 | 0.670 | 4996.280 | 0.738 | 0.516 |
| | | *m12* | 0.204 | 2.162 | 36.645 | 0.597 | 0.554 |
| | | *m21* | 0.153 | 5.423 | 39.303 | 0.503 | 0.505 |
| | ll1000 | *N1* | 0.652 | 0.222 | 9375.775 | 0.877 | 0.515 |
| | | *N2* | 0.663 | 0.258 | 9396.072 | 0.871 | 0.527 |
| | | *Nanc* | 0.979 | 0.047 | 1383.401 | 0.988 | 0.659 |
| | | *Tsep* | 0.513 | 0.575 | 4245.307 | 0.791 | 0.518 |
| | | *m12* | 0.239 | 2.926 | 35.944 | 0.640 | 0.540 |
| | | *m21* | 0.165 | 1.956 | 39.744 | 0.517 | 0.507 |
| nc20 | ll200 | *N1* | 0.588 | 0.327 | 10518.363 | 0.838 | 0.533 |
| | | *N2* | 0.641 | 0.204 | 9987.678 | 0.869 | 0.522 |
| | | *Nanc* | 0.971 | 0.048 | 2666.986 | 0.978 | 0.524 |
| | | *Tsep* | 0.436 | 0.722 | 5029.804 | 0.738 | 0.496 |
| | | *m12* | 0.180 | 2.541 | 39.894 | 0.535 | 0.491 |
| | | *m21* | 0.132 | 3.519 | 38.910 | 0.508 | 0.517 |
| | ll1000 | *N1* | 0.724 | 0.193 | 8262.431 | 0.917 | 0.525 |
| | | *N2* | 0.635 | 0.208 | 8673.650 | 0.888 | 0.545 |
| | | *Nanc* | 0.985 | 0.019 | 1259.140 | 0.990 | 0.598 |
| | | *Tsep* | 0.551 | 0.514 | 4210.267 | 0.796 | 0.552 |
| | | *m12* | 0.294 | 2.388 | 29.948 | 0.653 | 0.541 |
| | | *m21* | 0.192 | 2.356 | 36.484 | 0.557 | 0.539 |

| | | | | | | | |
|---|---|---|---|---|---|---|---|
| nc50 | ll200 | *N1* | 0.634 | 0.372 | 10247.254 | 0.808 | 0.532 |
| | | *N2* | 0.699 | 0.258 | 10416.350 | 0.819 | 0.489 |
| | | *Nanc* | 0.942 | 0.081 | 3315.686 | 0.954 | 0.512 |
| | | *Tsep* | 0.334 | 1.043 | 5415.652 | 0.704 | 0.485 |
| | | *m12* | 0.169 | 3.378 | 40.292 | 0.498 | 0.516 |
| | | *m21* | 0.153 | 2.111 | 38.613 | 0.492 | 0.507 |
| | ll1000 | *N1* | 0.746 | 0.103 | 7380.514 | 0.936 | 0.533 |
| | | *N2* | 0.724 | 0.127 | 8399.388 | 0.922 | 0.533 |
| | | *Nanc* | 0.966 | 0.048 | 1614.420 | 0.981 | 0.631 |
| | | *Tsep* | 0.488 | 0.467 | 4369.081 | 0.809 | 0.534 |
| | | *m12* | 0.260 | 1.748 | 33.814 | 0.643 | 0.535 |
| | | *m21* | 0.204 | 3.665 | 35.868 | 0.551 | 0.531 |

| | | **Coverage 2x** | | | | | |
|---|---|---|---|---|---|---|---|
| | | **Parameter** | **R2** | **Bias** | **RMSE** | **Factor2** | **Coverage50%** |
| nc10 | ll200 | *N1* | 0.668 | 0.285 | 8982.826 | 0.873 | 0.538 |
| | | *N2* | 0.548 | 0.332 | 10507.895 | 0.839 | 0.525 |
| | | *Nanc* | 0.963 | 0.040 | 2720.449 | 0.975 | 0.568 |
| | | *Tsep* | 0.383 | 0.701 | 4951.037 | 0.722 | 0.488 |
| | | *m12* | 0.198 | 2.869 | 34.815 | 0.603 | 0.541 |
| | | *m21* | 0.143 | 2.483 | 40.548 | 0.505 | 0.520 |
| | ll1000 | *N1* | 0.672 | 0.257 | 8977.575 | 0.879 | 0.502 |
| | | *N2* | 0.632 | 0.203 | 9747.851 | 0.872 | 0.502 |
| | | *Nanc* | 0.975 | 0.017 | 1114.137 | 0.994 | 0.661 |
| | | *Tsep* | 0.516 | 0.470 | 4179.326 | 0.798 | 0.539 |
| | | *m12* | 0.267 | 2.364 | 31.909 | 0.654 | 0.536 |
| | | *m21* | 0.163 | 4.465 | 37.771 | 0.531 | 0.520 |
| nc20 | ll200 | *N1* | 0.691 | 0.212 | 9139.610 | 0.885 | 0.501 |
| | | *N2* | 0.685 | 0.183 | 9300.829 | 0.878 | 0.537 |
| | | *Nanc* | 0.958 | 0.063 | 2348.220 | 0.977 | 0.544 |
| | | *Tsep* | 0.397 | 0.657 | 4771.680 | 0.768 | 0.515 |
| | | *m12* | 0.250 | 3.360 | 34.448 | 0.589 | 0.523 |
| | | *m21* | 0.152 | 4.472 | 37.814 | 0.489 | 0.491 |
| | ll1000 | *N1* | 0.703 | 0.212 | 8657.876 | 0.882 | 0.509 |
| | | *N2* | 0.644 | 0.181 | 8841.140 | 0.895 | 0.528 |
| | | *Nanc* | 0.985 | 0.021 | 1330.530 | 0.990 | 0.641 |
| | | *Tsep* | 0.534 | 0.497 | 4325.052 | 0.780 | 0.524 |
| | | *m12* | 0.264 | 2.785 | 33.939 | 0.627 | 0.503 |
| | | *m21* | 0.194 | 2.251 | 36.925 | 0.529 | 0.489 |
| nc50 | ll200 | *N1* | 0.825 | 0.236 | 8125.193 | 0.895 | 0.505 |
| | | *N2* | 0.701 | 0.233 | 9345.243 | 0.887 | 0.502 |
| | | *Nanc* | 0.936 | 0.084 | 3350.919 | 0.966 | 0.547 |
| | | *Tsep* | 0.338 | 0.621 | 5292.303 | 0.733 | 0.490 |
| | | *m12* | 0.239 | 2.691 | 34.784 | 0.579 | 0.517 |
| | | *m21* | 0.179 | 13.599 | 37.181 | 0.548 | 0.524 |
| | ll1000 | *N1* | 0.733 | 0.175 | 7535.241 | 0.923 | 0.523 |
| | | *N2* | 0.731 | 0.158 | 7988.443 | 0.921 | 0.515 |
| | | *Nanc* | 0.969 | 0.057 | 1753.408 | 0.974 | 0.648 |

| | | | | | | | |
|---|---|---|---|---|---|---|---|
| | | *Tsep* | 0.472 | 0.551 | 4234.751 | 0.807 | 0.563 |
| | | *m12* | 0.303 | 2.304 | 33.166 | 0.634 | 0.493 |
| | | *m21* | 0.210 | 24.453 | 37.708 | 0.543 | 0.514 |

| | | **Coverage 5x** | | | | | |
|---|---|---|---|---|---|---|---|
| | | **Parameter** | **R2** | **Bias** | **RMSE** | **Factor2** | **Coverage50%** |
| nc10 | ll200 | *N1* | 0.689 | 0.277 | 9196.270 | 0.876 | 0.509 |
| | | *N2* | 0.579 | 0.311 | 10645.089 | 0.831 | 0.503 |
| | | *Nanc* | 0.974 | 0.045 | 2500.958 | 0.975 | 0.566 |
| | | *Tsep* | 0.421 | 0.625 | 4911.774 | 0.756 | 0.519 |
| | | *m12* | 0.227 | 2.306 | 36.768 | 0.566 | 0.508 |
| | | *m21* | 0.159 | 3.653 | 38.352 | 0.500 | 0.500 |
| | ll1000 | *N1* | 0.701 | 0.253 | 8582.010 | 0.877 | 0.486 |
| | | *N2* | 0.662 | 0.224 | 8724.177 | 0.895 | 0.511 |
| | | *Nanc* | 0.980 | 0.034 | 1429.570 | 0.984 | 0.637 |
| | | *Tsep* | 0.537 | 0.454 | 4143.882 | 0.801 | 0.537 |
| | | *m12* | 0.270 | 13.950 | 30.847 | 0.661 | 0.546 |
| | | *m21* | 0.177 | 4.090 | 38.814 | 0.542 | 0.502 |
| nc20 | ll200 | *N1* | 0.744 | 0.198 | 8410.893 | 0.900 | 0.508 |
| | | *N2* | 0.711 | 0.166 | 8911.047 | 0.907 | 0.522 |
| | | *Nanc* | 0.949 | 0.061 | 2721.146 | 0.966 | 0.564 |
| | | *Tsep* | 0.405 | 0.616 | 5004.993 | 0.750 | 0.520 |
| | | *m12* | 0.248 | 2.999 | 36.054 | 0.619 | 0.511 |
| | | *m21* | 0.197 | 1.780 | 36.265 | 0.541 | 0.526 |
| | ll1000 | *N1* | 0.740 | 0.186 | 7615.388 | 0.909 | 0.556 |
| | | *N2* | 0.669 | 0.175 | 8939.756 | 0.918 | 0.494 |
| | | *Nanc* | 0.992 | 0.030 | 1442.174 | 0.985 | 0.650 |
| | | *Tsep* | 0.504 | 0.413 | 4394.375 | 0.810 | 0.530 |
| | | *m12* | 0.281 | 71.587 | 33.364 | 0.697 | 0.558 |
| | | *m21* | 0.157 | 3.236 | 43.540 | 0.575 | 0.529 |
| nc50 | ll200 | *N1* | 0.767 | 0.192 | 8068.101 | 0.882 | 0.490 |
| | | *N2* | 0.711 | 0.141 | 8757.809 | 0.916 | 0.516 |
| | | *Nanc* | 0.957 | 0.080 | 3161.402 | 0.958 | 0.557 |
| | | *Tsep* | 0.340 | 0.784 | 5189.352 | 0.734 | 0.498 |
| | | *m12* | 0.262 | 8.020 | 35.131 | 0.599 | 0.494 |
| | | *m21* | 0.190 | 11.098 | 36.162 | 0.539 | 0.496 |
| | ll1000 | *N1* | 0.799 | 0.123 | 6963.919 | 0.940 | 0.532 |
| | | *N2* | 0.709 | 0.146 | 8016.151 | 0.927 | 0.498 |
| | | *Nanc* | 0.986 | 0.037 | 1568.912 | 0.986 | 0.647 |
| | | *Tsep* | 0.496 | 0.494 | 4183.320 | 0.832 | 0.539 |
| | | *m12* | 0.306 | 7.787 | 32.411 | 0.666 | 0.525 |
| | | *m21* | 0.226 | 21.369 | 33.942 | 0.578 | 0.521 |

| | | **Coverage 30x** | | | | | |
|---|---|---|---|---|---|---|---|
| | | **Parameter** | **R2** | **Bias** | **RMSE** | **Factor2** | **Coverage50%** |
| nc10 | ll200 | *N1* | 0.655 | 0.252 | 9444.570 | 0.864 | 0.520 |
| | | *N2* | 0.640 | 0.269 | 9837.096 | 0.858 | 0.524 |
| | | *Nanc* | 0.973 | 0.048 | 2304.978 | 0.972 | 0.585 |

| | | | | | | | |
|---|---|---|---|---|---|---|---|
| | | *Tsep* | 0.417 | 0.559 | 4886.560 | 0.740 | 0.507 |
| | | *m12* | 0.227 | 3.177 | 35.416 | 0.578 | 0.496 |
| | | *m21* | 0.167 | 4.275 | 38.643 | 0.506 | 0.503 |
| | ll1000 | *N1* | 0.703 | 0.244 | 7829.534 | 0.907 | 0.524 |
| | | *N2* | 0.686 | 0.170 | 8311.963 | 0.912 | 0.535 |
| | | *Nanc* | 0.975 | 0.040 | 1161.914 | 0.984 | 0.670 |
| | | *Tsep* | 0.527 | 0.331 | 4058.775 | 0.820 | 0.558 |
| | | *m12* | 0.264 | 4.496 | 32.820 | 0.690 | 0.566 |
| | | *m21* | 0.167 | 5.145 | 31.451 | 0.606 | 0.503 |
| nc20 | ll200 | *N1* | 0.740 | 0.216 | 8358.351 | 0.886 | 0.481 |
| | | *N2* | 0.655 | 0.183 | 9419.081 | 0.883 | 0.511 |
| | | *Nanc* | 0.943 | 0.145 | 2908.572 | 0.958 | 0.570 |
| | | *Tsep* | 0.378 | 0.687 | 4804.651 | 0.748 | 0.510 |
| | | *m12* | 0.255 | 4.623 | 35.286 | 0.604 | 0.508 |
| | | *m21* | 0.197 | 2.054 | 34.490 | 0.547 | 0.521 |
| | ll1000 | *N1* | 0.770 | 0.217 | 7404.215 | 0.910 | 0.521 |
| | | *N2* | 0.688 | 0.179 | 8458.851 | 0.920 | 0.500 |
| | | *Nanc* | 0.983 | 0.038 | 1638.021 | 0.981 | 0.662 |
| | | *Tsep* | 0.498 | 0.457 | 4181.855 | 0.809 | 0.562 |
| | | *m12* | 0.336 | 1.612 | 27.841 | 0.697 | 0.552 |
| | | *m21* | 0.205 | 9.499 | 35.989 | 0.556 | 0.503 |
| nc50 | ll200 | *N1* | 0.809 | 0.179 | 7504.063 | 0.921 | 0.530 |
| | | *N2* | 0.764 | 0.142 | 8448.758 | 0.924 | 0.496 |
| | | *Nanc* | 0.932 | 0.153 | 3263.440 | 0.948 | 0.574 |
| | | *Tsep* | 0.341 | 0.659 | 5193.381 | 0.752 | 0.497 |
| | | *m12* | 0.233 | 1.497 | 34.498 | 0.646 | 0.542 |
| | | *m21* | 0.200 | 4.439 | 37.357 | 0.546 | 0.525 |
| | ll1000 | *N1* | 0.813 | 0.122 | 6706.854 | 0.947 | 0.529 |
| | | *N2* | 0.725 | 0.104 | 7704.226 | 0.945 | 0.525 |
| | | *Nanc* | 0.970 | 0.041 | 1678.281 | 0.984 | 0.669 |
| | | *Tsep* | 0.466 | 0.525 | 4308.982 | 0.821 | 0.553 |
| | | *m12* | 0.331 | 2.110 | 27.455 | 0.703 | 0.564 |
| | | *m21* | 0.220 | 34.586 | 34.958 | 0.584 | 0.525 |

**Supplementary Table 11. Accuracy of the estimated parameters of the Divergence with pulse of admixture model assessed by 1,000 pods.** Combinations of experimental parameters considering 1,000 loci.

| | | Coverage 1x | | | | | |
|---|---|---|---|---|---|---|---|
| | | **Parameter** | **R2** | **Bias** | **RMSE** | **Factor2** | **Coverage50%** |
| nc10 | ll200 | *N1* | 0.546 | 0.262 | 11130.775 | 0.848 | 0.487 |
| | | *N2* | 0.599 | 0.214 | 10091.491 | 0.876 | 0.509 |
| | | *Nanc* | 0.974 | 0.017 | 1825.056 | 0.994 | 0.540 |
| | | *Tadm* | 0.475 | 0.170 | 466.348 | 0.938 | 0.511 |
| | | *Tsep* | 0.605 | 0.155 | 1702.689 | 0.929 | 0.538 |
| | | *adm12* | 0.076 | 0.146 | 0.043 | 0.930 | 0.526 |
| | | *adm21* | 0.077 | 0.151 | 0.045 | 0.924 | 0.502 |
| | ll1000 | *N1* | 0.672 | 0.177 | 8194.508 | 0.923 | 0.567 |
| | | *N2* | 0.716 | 0.124 | 8229.567 | 0.948 | 0.554 |
| | | *Nanc* | 0.985 | 0.005 | 1271.255 | 0.997 | 0.634 |
| | | *Tadm* | 0.525 | 0.134 | 424.338 | 0.952 | 0.510 |
| | | *Tsep* | 0.658 | 0.122 | 1401.779 | 0.948 | 0.587 |
| | | *adm12* | 0.067 | 0.149 | 0.043 | 0.922 | 0.499 |
| | | *adm21* | 0.065 | 0.163 | 0.044 | 0.915 | 0.493 |
| nc20 | ll200 | *N1* | 0.644 | 0.180 | 10111.944 | 0.901 | 0.492 |
| | | *N2* | 0.658 | 0.158 | 10211.257 | 0.893 | 0.480 |
| | | *Nanc* | 0.969 | 0.006 | 2005.487 | 0.997 | 0.535 |
| | | *Tadm* | 0.552 | 0.137 | 441.278 | 0.940 | 0.504 |
| | | *Tsep* | 0.724 | 0.087 | 1457.427 | 0.952 | 0.530 |
| | | *adm12* | 0.098 | 0.161 | 0.045 | 0.905 | 0.480 |
| | | *adm21* | 0.108 | 0.144 | 0.045 | 0.914 | 0.491 |
| | ll1000 | *N1* | 0.721 | 0.105 | 8001.344 | 0.945 | 0.548 |
| | | *N2* | 0.717 | 0.100 | 8057.902 | 0.944 | 0.533 |
| | | *Nanc* | 0.992 | 0.002 | 1254.815 | 1.000 | 0.604 |
| | | *Tadm* | 0.559 | 0.113 | 398.900 | 0.965 | 0.512 |
| | | *Tsep* | 0.722 | 0.056 | 1223.826 | 0.986 | 0.559 |
| | | *adm12* | 0.073 | 0.140 | 0.043 | 0.927 | 0.511 |
| | | *adm21* | 0.082 | 0.160 | 0.043 | 0.922 | 0.527 |
| nc50 | ll200 | *N1* | 0.654 | 0.154 | 9928.273 | 0.898 | 0.491 |
| | | *N2* | 0.556 | 0.357 | 11223.367 | 0.816 | 0.494 |
| | | *Nanc* | 0.990 | 0.022 | 3650.681 | 0.973 | 0.478 |
| | | *Tadm* | 0.521 | 0.150 | 495.954 | 0.922 | 0.475 |
| | | *Tsep* | 0.555 | 0.130 | 2015.829 | 0.905 | 0.511 |
| | | *adm12* | 0.134 | 0.154 | 0.046 | 0.912 | 0.493 |
| | | *adm21* | 0.137 | 0.147 | 0.046 | 0.901 | 0.496 |
| | ll1000 | *N1* | 0.731 | 0.115 | 7573.528 | 0.951 | 0.544 |
| | | *N2* | 0.787 | 0.131 | 7474.297 | 0.952 | 0.551 |
| | | *Nanc* | 0.972 | 0.013 | 1765.633 | 0.990 | 0.568 |
| | | *Tadm* | 0.439 | 0.156 | 440.803 | 0.930 | 0.525 |
| | | *Tsep* | 0.522 | 0.098 | 1650.991 | 0.948 | 0.583 |
| | | *adm12* | 0.067 | 0.146 | 0.045 | 0.914 | 0.484 |
| | | *adm21* | 0.081 | 0.153 | 0.044 | 0.918 | 0.498 |

| | | Coverage 2x | | | | | |
|---|---|---|---|---|---|---|---|
| | | Parameter | R2 | Bias | RMSE | Factor2 | Coverage50% |
| nc10 | ll200 | *N1* | 0.605 | 0.173 | 10167.407 | 0.889 | 0.475 |
| | | *N2* | 0.566 | 0.193 | 10406.735 | 0.905 | 0.496 |
| | | *Nanc* | 0.992 | 0.021 | 1813.034 | 0.993 | 0.527 |
| | | *Tadm* | 0.538 | 0.130 | 429.489 | 0.949 | 0.519 |
| | | *Tsep* | 0.697 | 0.100 | 1484.007 | 0.948 | 0.527 |
| | | *adm12* | 0.087 | 0.139 | 0.044 | 0.933 | 0.505 |
| | | *adm21* | 0.097 | 0.124 | 0.045 | 0.935 | 0.464 |
| | ll1000 | *N1* | 0.737 | 0.100 | 8002.048 | 0.963 | 0.584 |
| | | *N2* | 0.742 | 0.099 | 7963.528 | 0.952 | 0.556 |
| | | *Nanc* | 0.984 | 0.005 | 1166.919 | 1.000 | 0.644 |
| | | *Tadm* | 0.574 | 0.101 | 383.733 | 0.965 | 0.527 |
| | | *Tsep* | 0.733 | 0.056 | 1126.227 | 0.987 | 0.560 |
| | | *adm12* | 0.074 | 0.144 | 0.043 | 0.931 | 0.533 |
| | | *adm21* | 0.074 | 0.156 | 0.044 | 0.924 | 0.480 |
| nc20 | ll200 | *N1* | 0.612 | 0.212 | 10169.393 | 0.870 | 0.533 |
| | | *N2* | 0.599 | 0.193 | 9914.493 | 0.889 | 0.525 |
| | | *Nanc* | 0.979 | 0.013 | 2083.050 | 0.993 | 0.547 |
| | | *Tadm* | 0.587 | 0.089 | 428.780 | 0.960 | 0.494 |
| | | *Tsep* | 0.728 | 0.082 | 1372.957 | 0.972 | 0.527 |
| | | *adm12* | 0.096 | 0.146 | 0.044 | 0.917 | 0.509 |
| | | *adm21* | 0.094 | 0.143 | 0.044 | 0.922 | 0.518 |
| | ll1000 | *N1* | 0.735 | 0.087 | 7272.509 | 0.968 | 0.556 |
| | | *N2* | 0.691 | 0.085 | 7633.071 | 0.962 | 0.554 |
| | | *Nanc* | 0.982 | 0.002 | 1364.099 | 1.000 | 0.657 |
| | | *Tadm* | 0.608 | 0.050 | 385.242 | 0.975 | 0.494 |
| | | *Tsep* | 0.809 | 0.036 | 1028.959 | 0.992 | 0.594 |
| | | *adm12* | 0.076 | 0.162 | 0.043 | 0.915 | 0.511 |
| | | *adm21* | 0.089 | 0.160 | 0.043 | 0.918 | 0.525 |
| nc50 | ll200 | *N1* | 0.749 | 0.117 | 8569.528 | 0.925 | 0.494 |
| | | *N2* | 0.584 | 0.264 | 10501.004 | 0.851 | 0.521 |
| | | *Nanc* | 0.957 | 0.020 | 3187.190 | 0.974 | 0.517 |
| | | *Tadm* | 0.425 | 0.148 | 486.768 | 0.932 | 0.497 |
| | | *Tsep* | 0.443 | 0.128 | 2105.536 | 0.902 | 0.526 |
| | | *adm12* | 0.106 | 0.129 | 0.046 | 0.926 | 0.473 |
| | | *adm21* | 0.111 | 0.127 | 0.045 | 0.923 | 0.483 |
| | ll1000 | *N1* | 0.752 | 0.084 | 6634.115 | 0.973 | 0.586 |
| | | *N2* | 0.769 | 0.085 | 7223.169 | 0.968 | 0.565 |
| | | *Nanc* | 0.980 | 0.011 | 1809.860 | 0.993 | 0.586 |
| | | *Tadm* | 0.557 | 0.103 | 398.087 | 0.968 | 0.522 |
| | | *Tsep* | 0.678 | 0.063 | 1336.505 | 0.979 | 0.608 |
| | | *adm12* | 0.066 | 0.153 | 0.044 | 0.924 | 0.484 |
| | | *adm21* | 0.070 | 0.136 | 0.043 | 0.926 | 0.517 |

| | | Coverage 5x | | | | | |
|---|---|---|---|---|---|---|---|
| | | Parameter | R2 | Bias | RMSE | Factor2 | Coverage50% |
| nc10 | ll200 | *N1* | 0.606 | 0.189 | 9771.302 | 0.890 | 0.560 |

| | | | | | | | |
|---|---|---|---|---|---|---|---|
| | | *N2* | 0.661 | 0.154 | 9756.026 | 0.902 | 0.513 |
| | | *Nanc* | 0.983 | 0.009 | 1899.363 | 0.998 | 0.559 |
| | | *Tadm* | 0.566 | 0.110 | 426.074 | 0.959 | 0.497 |
| | | *Tsep* | 0.695 | 0.073 | 1392.094 | 0.972 | 0.525 |
| | | *adm12* | 0.092 | 0.147 | 0.042 | 0.927 | 0.538 |
| | | *adm21* | 0.099 | 0.130 | 0.043 | 0.932 | 0.505 |
| | ll1000 | *N1* | 0.757 | 0.071 | 7316.149 | 0.970 | 0.566 |
| | | *N2* | 0.695 | 0.069 | 7507.254 | 0.973 | 0.556 |
| | | *Nanc* | 0.983 | 0.004 | 1182.485 | 0.999 | 0.686 |
| | | *Tadm* | 0.652 | 0.084 | 359.958 | 0.971 | 0.519 |
| | | *Tsep* | 0.782 | 0.042 | 991.977 | 0.989 | 0.603 |
| | | *adm12* | 0.080 | 0.157 | 0.043 | 0.919 | 0.504 |
| | | *adm21* | 0.076 | 0.147 | 0.043 | 0.927 | 0.500 |
| nc20 | ll200 | *N1* | 0.708 | 0.111 | 8539.791 | 0.947 | 0.518 |
| | | *N2* | 0.652 | 0.151 | 9316.438 | 0.917 | 0.505 |
| | | *Nanc* | 0.976 | 0.019 | 2351.998 | 0.990 | 0.571 |
| | | *Tadm* | 0.631 | 0.077 | 400.676 | 0.964 | 0.484 |
| | | *Tsep* | 0.753 | 0.060 | 1297.350 | 0.986 | 0.520 |
| | | *adm12* | 0.103 | 0.144 | 0.044 | 0.923 | 0.508 |
| | | *adm21* | 0.094 | 0.200 | 0.045 | 0.885 | 0.488 |
| | ll1000 | *N1* | 0.792 | 0.102 | 6593.486 | 0.973 | 0.571 |
| | | *N2* | 0.764 | 0.109 | 7028.718 | 0.966 | 0.551 |
| | | *Nanc* | 0.981 | 0.008 | 1390.129 | 0.996 | 0.644 |
| | | *Tadm* | 0.675 | 0.080 | 349.530 | 0.980 | 0.539 |
| | | *Tsep* | 0.806 | 0.024 | 1049.131 | 0.991 | 0.568 |
| | | *adm12* | 0.077 | 0.165 | 0.044 | 0.914 | 0.482 |
| | | *adm21* | 0.101 | 0.173 | 0.042 | 0.922 | 0.506 |
| nc50 | ll200 | *N1* | 0.710 | 0.160 | 8195.573 | 0.928 | 0.540 |
| | | *N2* | 0.775 | 0.107 | 7717.886 | 0.949 | 0.553 |
| | | *Nanc* | 0.956 | 0.011 | 2912.344 | 0.985 | 0.545 |
| | | *Tadm* | 0.617 | 0.095 | 398.707 | 0.975 | 0.493 |
| | | *Tsep* | 0.661 | 0.033 | 1504.706 | 0.975 | 0.521 |
| | | *adm12* | 0.089 | 0.156 | 0.044 | 0.919 | 0.500 |
| | | *adm21* | 0.085 | 0.144 | 0.045 | 0.915 | 0.494 |
| | ll1000 | *N1* | 0.818 | 0.089 | 6160.797 | 0.976 | 0.596 |
| | | *N2* | 0.786 | 0.088 | 6650.572 | 0.968 | 0.591 |
| | | *Nanc* | 0.968 | 0.007 | 1720.445 | 0.991 | 0.626 |
| | | *Tadm* | 0.675 | 0.068 | 351.189 | 0.990 | 0.490 |
| | | *Tsep* | 0.739 | 0.043 | 1237.254 | 0.988 | 0.548 |
| | | *adm12* | 0.087 | 0.137 | 0.042 | 0.940 | 0.511 |
| | | *adm21* | 0.092 | 0.158 | 0.043 | 0.920 | 0.522 |

| | | Coverage 30x | | | | | |
|---|---|---|---|---|---|---|---|
| | | **Parameter** | **R2** | **Bias** | **RMSE** | **Factor2** | **Coverage50%** |
| nc10 | ll200 | *N1* | 0.606 | 0.183 | 9823.368 | 0.899 | 0.516 |
| | | *N2* | 0.708 | 0.153 | 9161.712 | 0.909 | 0.510 |
| | | *Nanc* | 0.976 | 0.006 | 2026.205 | 0.993 | 0.540 |
| | | *Tadm* | 0.548 | 0.122 | 434.992 | 0.950 | 0.491 |

| | | | | | | | |
|---|---|---|---|---|---|---|---|
| | | *Tsep* | 0.663 | 0.087 | 1484.241 | 0.966 | 0.536 |
| | | *adm12* | 0.095 | 0.164 | 0.043 | 0.925 | 0.516 |
| | | *adm21* | 0.088 | 0.167 | 0.045 | 0.916 | 0.492 |
| | ll1000 | *N1* | 0.739 | 0.080 | 7097.446 | 0.973 | 0.558 |
| | | *N2* | 0.763 | 0.097 | 6819.645 | 0.975 | 0.568 |
| | | *Nanc* | 0.984 | 0.001 | 1315.735 | 1.000 | 0.675 |
| | | *Tadm* | 0.600 | 0.083 | 376.392 | 0.978 | 0.504 |
| | | *Tsep* | 0.759 | 0.048 | 936.737 | 0.990 | 0.645 |
| | | *adm12* | 0.073 | 0.146 | 0.043 | 0.929 | 0.509 |
| | | *adm21* | 0.072 | 0.147 | 0.043 | 0.934 | 0.497 |
| nc20 | ll200 | *N1* | 0.729 | 0.105 | 8367.586 | 0.950 | 0.533 |
| | | *N2* | 0.637 | 0.128 | 8624.309 | 0.935 | 0.519 |
| | | *Nanc* | 0.963 | 0.022 | 2316.546 | 0.991 | 0.554 |
| | | *Tadm* | 0.628 | 0.103 | 381.771 | 0.968 | 0.551 |
| | | *Tsep* | 0.738 | 0.064 | 1242.293 | 0.988 | 0.552 |
| | | *adm12* | 0.088 | 0.190 | 0.045 | 0.901 | 0.500 |
| | | *adm21* | 0.089 | 0.162 | 0.043 | 0.914 | 0.510 |
| | ll1000 | *N1* | 0.802 | 0.067 | 6205.341 | 0.986 | 0.574 |
| | | *N2* | 0.805 | 0.077 | 6223.132 | 0.982 | 0.567 |
| | | *Nanc* | 0.970 | 0.003 | 1474.738 | 0.998 | 0.669 |
| | | *Tadm* | 0.654 | 0.057 | 352.295 | 0.988 | 0.509 |
| | | *Tsep* | 0.787 | 0.022 | 945.127 | 0.996 | 0.623 |
| | | *adm12* | 0.085 | 0.135 | 0.042 | 0.930 | 0.510 |
| | | *adm21* | 0.081 | 0.131 | 0.042 | 0.928 | 0.519 |
| nc50 | ll200 | *N1* | 0.742 | 0.107 | 7564.815 | 0.959 | 0.509 |
| | | *N2* | 0.745 | 0.144 | 7976.194 | 0.943 | 0.515 |
| | | *Nanc* | 0.949 | 0.000 | 3116.077 | 0.990 | 0.551 |
| | | *Tadm* | 0.663 | 0.087 | 370.928 | 0.976 | 0.495 |
| | | *Tsep* | 0.712 | 0.041 | 1354.106 | 0.984 | 0.557 |
| | | *adm12* | 0.097 | 0.145 | 0.043 | 0.918 | 0.525 |
| | | *adm21* | 0.084 | 0.158 | 0.044 | 0.916 | 0.478 |
| | ll1000 | *N1* | 0.817 | 0.056 | 5614.259 | 0.993 | 0.572 |
| | | *N2* | 0.839 | 0.057 | 5651.357 | 0.991 | 0.579 |
| | | *Nanc* | 0.974 | 0.002 | 1559.145 | 0.999 | 0.667 |
| | | *Tadm* | 0.717 | 0.059 | 335.390 | 0.991 | 0.533 |
| | | *Tsep* | 0.789 | 0.015 | 1093.603 | 0.993 | 0.572 |
| | | *adm12* | 0.089 | 0.152 | 0.043 | 0.920 | 0.498 |
| | | *adm21* | 0.093 | 0.146 | 0.042 | 0.931 | 0.531 |

**Supplementary Table 12. Accuracy of the estimated parameters of the Divergence with pulse of admixture model assessed by 1,000 pods.** Combinations of experimental parameters considering 5,000 loci.

| | | Coverage 1x | | | | | |
|---|---|---|---|---|---|---|---|
| | | **Parameter** | **R2** | **Bias** | **RMSE** | **Factor2** | **Coverage50%** |
| nc10 | ll200 | *N1* | 0.756 | 0.116 | 8291.115 | 0.953 | 0.531 |
| | | *N2* | 0.646 | 0.174 | 9134.845 | 0.911 | 0.541 |
| | | *Nanc* | 0.986 | 0.004 | 1148.603 | 0.998 | 0.557 |
| | | *Tadm* | 0.533 | 0.110 | 434.016 | 0.952 | 0.505 |
| | | *Tsep* | 0.701 | 0.099 | 1391.240 | 0.953 | 0.529 |
| | | *adm12* | 0.081 | 0.172 | 0.044 | 0.918 | 0.479 |
| | | *adm21* | 0.080 | 0.150 | 0.044 | 0.914 | 0.507 |
| | ll1000 | *N1* | 0.771 | 0.072 | 6647.890 | 0.978 | 0.565 |
| | | *N2* | 0.826 | 0.066 | 5883.734 | 0.985 | 0.573 |
| | | *Nanc* | 0.994 | 0.002 | 695.711 | 1.000 | 0.681 |
| | | *Tadm* | 0.547 | 0.104 | 403.042 | 0.975 | 0.467 |
| | | *Tsep* | 0.720 | 0.059 | 978.607 | 0.988 | 0.640 |
| | | *adm12* | 0.086 | 0.137 | 0.043 | 0.920 | 0.500 |
| | | *adm21* | 0.083 | 0.137 | 0.043 | 0.935 | 0.492 |
| nc20 | ll200 | *N1* | 0.670 | 0.163 | 8440.250 | 0.927 | 0.523 |
| | | *N2* | 0.756 | 0.078 | 8226.346 | 0.949 | 0.498 |
| | | *Nanc* | 0.997 | 0.005 | 1349.302 | 0.993 | 0.535 |
| | | *Tadm* | 0.577 | 0.124 | 419.847 | 0.948 | 0.516 |
| | | *Tsep* | 0.704 | 0.080 | 1377.175 | 0.966 | 0.551 |
| | | *adm12* | 0.104 | 0.181 | 0.045 | 0.914 | 0.479 |
| | | *adm21* | 0.121 | 0.151 | 0.044 | 0.913 | 0.509 |
| | ll1000 | *N1* | 0.829 | 0.058 | 5823.267 | 0.995 | 0.603 |
| | | *N2* | 0.814 | 0.062 | 6025.304 | 0.988 | 0.617 |
| | | *Nanc* | 0.993 | 0.002 | 745.688 | 0.998 | 0.668 |
| | | *Tadm* | 0.660 | 0.064 | 372.707 | 0.984 | 0.487 |
| | | *Tsep* | 0.857 | 0.037 | 858.195 | 0.993 | 0.602 |
| | | *adm12* | 0.100 | 0.177 | 0.044 | 0.913 | 0.486 |
| | | *adm21* | 0.090 | 0.168 | 0.044 | 0.927 | 0.488 |
| nc50 | ll200 | *N1* | 0.618 | 0.219 | 9066.509 | 0.887 | 0.519 |
| | | *N2* | 0.760 | 0.090 | 8247.314 | 0.936 | 0.497 |
| | | *Nanc* | 0.985 | 0.005 | 2025.551 | 0.992 | 0.552 |
| | | *Tadm* | 0.578 | 0.130 | 446.197 | 0.939 | 0.512 |
| | | *Tsep* | 0.630 | 0.106 | 1736.961 | 0.918 | 0.528 |
| | | *adm12* | 0.107 | 0.173 | 0.045 | 0.907 | 0.475 |
| | | *adm21* | 0.120 | 0.158 | 0.046 | 0.901 | 0.503 |
| | ll1000 | *N1* | 0.844 | 0.053 | 5236.700 | 0.988 | 0.600 |
| | | *N2* | 0.855 | 0.063 | 5225.656 | 0.988 | 0.599 |
| | | *Nanc* | 0.980 | 0.002 | 935.789 | 1.000 | 0.640 |
| | | *Tadm* | 0.650 | 0.081 | 389.137 | 0.972 | 0.504 |
| | | *Tsep* | 0.707 | 0.064 | 1199.189 | 0.987 | 0.596 |
| | | *adm12* | 0.089 | 0.139 | 0.043 | 0.934 | 0.516 |
| | | *adm21* | 0.098 | 0.123 | 0.044 | 0.922 | 0.517 |

| | | Coverage 2x | | | | | |
|---|---|---|---|---|---|---|---|
| | | Parameter | R2 | Bias | RMSE | Factor2 | Coverage50% |
| nc10 | ll200 | *N1* | 0.773 | 0.088 | 7339.078 | 0.967 | 0.561 |
| | | *N2* | 0.779 | 0.111 | 7043.112 | 0.961 | 0.605 |
| | | *Nanc* | 0.988 | 0.008 | 1088.319 | 0.997 | 0.602 |
| | | *Tadm* | 0.568 | 0.095 | 390.686 | 0.965 | 0.501 |
| | | *Tsep* | 0.751 | 0.077 | 1177.906 | 0.973 | 0.585 |
| | | *adm12* | 0.085 | 0.133 | 0.043 | 0.923 | 0.515 |
| | | *adm21* | 0.078 | 0.148 | 0.045 | 0.920 | 0.488 |
| | ll1000 | *N1* | 0.799 | 0.037 | 5861.431 | 0.989 | 0.598 |
| | | *N2* | 0.813 | 0.057 | 6125.758 | 0.986 | 0.580 |
| | | *Nanc* | 0.997 | 0.002 | 678.179 | 1.000 | 0.712 |
| | | *Tadm* | 0.672 | 0.077 | 363.650 | 0.983 | 0.481 |
| | | *Tsep* | 0.830 | 0.034 | 839.707 | 0.993 | 0.620 |
| | | *adm12* | 0.103 | 0.152 | 0.042 | 0.923 | 0.500 |
| | | *adm21* | 0.099 | 0.161 | 0.043 | 0.925 | 0.513 |
| nc20 | ll200 | *N1* | 0.776 | 0.102 | 7845.828 | 0.952 | 0.551 |
| | | *N2* | 0.730 | 0.126 | 8120.566 | 0.941 | 0.554 |
| | | *Nanc* | 0.991 | 0.002 | 1403.866 | 1.000 | 0.577 |
| | | *Tadm* | 0.638 | 0.087 | 388.444 | 0.978 | 0.490 |
| | | *Tsep* | 0.778 | 0.034 | 1074.476 | 0.990 | 0.577 |
| | | *adm12* | 0.113 | 0.136 | 0.044 | 0.924 | 0.503 |
| | | *adm21* | 0.107 | 0.149 | 0.044 | 0.922 | 0.500 |
| | ll1000 | *N1* | 0.815 | 0.055 | 5906.411 | 0.987 | 0.583 |
| | | *N2* | 0.798 | 0.059 | 5881.622 | 0.985 | 0.591 |
| | | *Nanc* | 0.994 | 0.001 | 842.357 | 0.999 | 0.675 |
| | | *Tadm* | 0.678 | 0.081 | 371.951 | 0.978 | 0.470 |
| | | *Tsep* | 0.865 | 0.045 | 860.378 | 0.993 | 0.580 |
| | | *adm12* | 0.093 | 0.172 | 0.042 | 0.925 | 0.519 |
| | | *adm21* | 0.096 | 0.163 | 0.043 | 0.924 | 0.502 |
| nc50 | ll200 | *N1* | 0.715 | 0.114 | 7655.604 | 0.950 | 0.516 |
| | | *N2* | 0.717 | 0.102 | 7897.550 | 0.948 | 0.527 |
| | | *Nanc* | 0.985 | 0.007 | 2314.283 | 0.990 | 0.554 |
| | | *Tadm* | 0.630 | 0.094 | 386.857 | 0.975 | 0.493 |
| | | *Tsep* | 0.727 | 0.026 | 1351.056 | 0.984 | 0.518 |
| | | *adm12* | 0.119 | 0.158 | 0.044 | 0.926 | 0.508 |
| | | *adm21* | 0.119 | 0.125 | 0.044 | 0.922 | 0.509 |
| | ll1000 | *N1* | 0.811 | 0.051 | 5546.903 | 0.992 | 0.594 |
| | | *N2* | 0.853 | 0.044 | 4628.994 | 0.997 | 0.618 |
| | | *Nanc* | 0.985 | 0.006 | 1098.199 | 0.999 | 0.641 |
| | | *Tadm* | 0.670 | 0.068 | 361.052 | 0.989 | 0.526 |
| | | *Tsep* | 0.734 | 0.039 | 1181.380 | 0.986 | 0.610 |
| | | *adm12* | 0.103 | 0.135 | 0.043 | 0.927 | 0.505 |
| | | *adm21* | 0.101 | 0.163 | 0.044 | 0.912 | 0.490 |

| | | Coverage 5x | | | | | |
|---|---|---|---|---|---|---|---|
| | | Parameter | R2 | Bias | RMSE | Factor2 | Coverage50% |
| nc10 | ll200 | *N1* | 0.753 | 0.106 | 8268.709 | 0.948 | 0.514 |

| | | Parameter | R2 | Bias | RMSE | Factor2 | Coverage50% |
|---|---|---|---|---|---|---|---|
| | | *N2* | 0.710 | 0.151 | 8307.573 | 0.936 | 0.552 |
| | | *Nanc* | 1.000 | 0.002 | 1128.252 | 0.998 | 0.581 |
| | | *Tadm* | 0.672 | 0.085 | 373.503 | 0.969 | 0.535 |
| | | *Tsep* | 0.789 | 0.050 | 1026.727 | 0.987 | 0.568 |
| | | *adm12* | 0.107 | 0.154 | 0.044 | 0.920 | 0.499 |
| | | *adm21* | 0.092 | 0.142 | 0.045 | 0.920 | 0.469 |
| | ll1000 | *N1* | 0.796 | 0.033 | 5726.459 | 0.992 | 0.602 |
| | | *N2* | 0.844 | 0.041 | 5831.839 | 0.995 | 0.578 |
| | | *Nanc* | 0.991 | 0.000 | 681.811 | 1.000 | 0.714 |
| | | *Tadm* | 0.668 | 0.066 | 366.881 | 0.987 | 0.466 |
| | | *Tsep* | 0.844 | 0.033 | 777.561 | 0.996 | 0.608 |
| | | *adm12* | 0.085 | 0.142 | 0.042 | 0.936 | 0.502 |
| | | *adm21* | 0.097 | 0.157 | 0.044 | 0.912 | 0.482 |
| nc20 | ll200 | *N1* | 0.812 | 0.085 | 7019.189 | 0.973 | 0.537 |
| | | *N2* | 0.741 | 0.096 | 7089.348 | 0.973 | 0.550 |
| | | *Nanc* | 0.977 | 0.012 | 1559.038 | 0.996 | 0.582 |
| | | *Tadm* | 0.669 | 0.069 | 364.814 | 0.991 | 0.480 |
| | | *Tsep* | 0.834 | 0.039 | 976.436 | 0.996 | 0.562 |
| | | *adm12* | 0.097 | 0.137 | 0.044 | 0.926 | 0.498 |
| | | *adm21* | 0.103 | 0.144 | 0.043 | 0.934 | 0.510 |
| | ll1000 | *N1* | 0.828 | 0.051 | 5122.222 | 0.988 | 0.612 |
| | | *N2* | 0.815 | 0.056 | 5475.032 | 0.985 | 0.614 |
| | | *Nanc* | 0.989 | 0.002 | 856.310 | 1.000 | 0.666 |
| | | *Tadm* | 0.680 | 0.064 | 357.623 | 0.985 | 0.479 |
| | | *Tsep* | 0.827 | 0.019 | 878.846 | 0.998 | 0.550 |
| | | *adm12* | 0.099 | 0.165 | 0.045 | 0.909 | 0.463 |
| | | *adm21* | 0.105 | 0.137 | 0.043 | 0.927 | 0.502 |
| nc50 | ll200 | *N1* | 0.793 | 0.109 | 7061.329 | 0.959 | 0.547 |
| | | *N2* | 0.766 | 0.092 | 7210.604 | 0.962 | 0.513 |
| | | *Nanc* | 0.964 | 0.013 | 2187.226 | 0.994 | 0.566 |
| | | *Tadm* | 0.704 | 0.044 | 371.245 | 0.983 | 0.498 |
| | | *Tsep* | 0.746 | 0.024 | 1275.213 | 0.989 | 0.534 |
| | | *adm12* | 0.102 | 0.156 | 0.045 | 0.917 | 0.481 |
| | | *adm21* | 0.099 | 0.153 | 0.045 | 0.920 | 0.475 |
| | ll1000 | *N1* | 0.876 | 0.054 | 4784.508 | 0.993 | 0.647 |
| | | *N2* | 0.861 | 0.049 | 4751.239 | 0.994 | 0.647 |
| | | *Nanc* | 0.985 | 0.008 | 1052.834 | 0.997 | 0.671 |
| | | *Tadm* | 0.678 | 0.072 | 337.184 | 0.989 | 0.524 |
| | | *Tsep* | 0.791 | 0.021 | 1094.193 | 0.994 | 0.559 |
| | | *adm12* | 0.116 | 0.136 | 0.042 | 0.930 | 0.507 |
| | | *adm21* | 0.114 | 0.137 | 0.043 | 0.931 | 0.499 |

| | | Coverage 30x | | | | | |
|---|---|---|---|---|---|---|---|
| | | **Parameter** | **R2** | **Bias** | **RMSE** | **Factor2** | **Coverage50%** |
| nc10 | ll200 | *N1* | 0.724 | 0.108 | 7912.677 | 0.959 | 0.545 |
| | | *N2* | 0.754 | 0.083 | 7804.380 | 0.969 | 0.527 |
| | | *Nanc* | 0.984 | 0.000 | 1318.414 | 0.999 | 0.575 |
| | | *Tadm* | 0.617 | 0.086 | 387.329 | 0.968 | 0.488 |

| | | | | | | | |
|---|---|---|---|---|---|---|---|
| | | *Tsep* | 0.742 | 0.049 | 1097.765 | 0.985 | 0.589 |
| | | *adm12* | 0.105 | 0.146 | 0.044 | 0.929 | 0.490 |
| | | *adm21* | 0.102 | 0.136 | 0.044 | 0.930 | 0.492 |
| | ll1000 | *N1* | 0.815 | 0.055 | 5372.516 | 0.993 | 0.589 |
| | | *N2* | 0.822 | 0.050 | 5364.411 | 0.992 | 0.585 |
| | | *Nanc* | 0.987 | 0.001 | 721.031 | 1.000 | 0.711 |
| | | *Tadm* | 0.643 | 0.086 | 359.445 | 0.983 | 0.505 |
| | | *Tsep* | 0.844 | 0.034 | 781.658 | 0.991 | 0.646 |
| | | *adm12* | 0.092 | 0.161 | 0.042 | 0.925 | 0.521 |
| | | *adm21* | 0.100 | 0.163 | 0.041 | 0.924 | 0.517 |
| nc20 | ll200 | *N1* | 0.819 | 0.084 | 6709.087 | 0.984 | 0.552 |
| | | *N2* | 0.780 | 0.105 | 7503.843 | 0.962 | 0.557 |
| | | *Nanc* | 0.974 | 0.003 | 1611.378 | 0.999 | 0.589 |
| | | *Tadm* | 0.651 | 0.080 | 361.369 | 0.972 | 0.499 |
| | | *Tsep* | 0.769 | 0.026 | 1039.546 | 0.992 | 0.572 |
| | | *adm12* | 0.109 | 0.123 | 0.044 | 0.919 | 0.500 |
| | | *adm21* | 0.105 | 0.138 | 0.044 | 0.942 | 0.484 |
| | ll1000 | *N1* | 0.849 | 0.054 | 5179.492 | 0.990 | 0.591 |
| | | *N2* | 0.841 | 0.056 | 5077.558 | 0.993 | 0.609 |
| | | *Nanc* | 0.991 | 0.003 | 800.275 | 0.999 | 0.712 |
| | | *Tadm* | 0.718 | 0.062 | 350.865 | 0.990 | 0.504 |
| | | *Tsep* | 0.863 | 0.023 | 882.932 | 0.996 | 0.578 |
| | | *adm12* | 0.111 | 0.137 | 0.042 | 0.935 | 0.505 |
| | | *adm21* | 0.096 | 0.166 | 0.044 | 0.918 | 0.465 |
| nc50 | ll200 | *N1* | 0.739 | 0.092 | 6888.367 | 0.967 | 0.536 |
| | | *N2* | 0.840 | 0.085 | 6545.927 | 0.976 | 0.529 |
| | | *Nanc* | 0.972 | 0.006 | 1966.477 | 0.996 | 0.576 |
| | | *Tadm* | 0.722 | 0.059 | 340.551 | 0.988 | 0.521 |
| | | *Tsep* | 0.791 | 0.016 | 1175.963 | 0.989 | 0.543 |
| | | *adm12* | 0.122 | 0.119 | 0.042 | 0.933 | 0.523 |
| | | *adm21* | 0.103 | 0.139 | 0.043 | 0.937 | 0.500 |
| | ll1000 | *N1* | 0.876 | 0.024 | 4378.537 | 0.996 | 0.620 |
| | | *N2* | 0.862 | 0.030 | 4576.962 | 0.994 | 0.618 |
| | | *Nanc* | 0.990 | 0.003 | 968.492 | 0.997 | 0.688 |
| | | *Tadm* | 0.719 | 0.054 | 324.168 | 0.992 | 0.518 |
| | | *Tsep* | 0.817 | 0.001 | 1027.107 | 0.998 | 0.526 |
| | | *adm12* | 0.127 | 0.117 | 0.042 | 0.944 | 0.500 |
| | | *adm21* | 0.115 | 0.135 | 0.042 | 0.938 | 0.529 |

**Supplementary Figure 1. Proportion of True Positives for the one-population models, obtained correcting the pods through the genotype likelihood computed by ANGSD.** The plots have the same features of Figure 1.

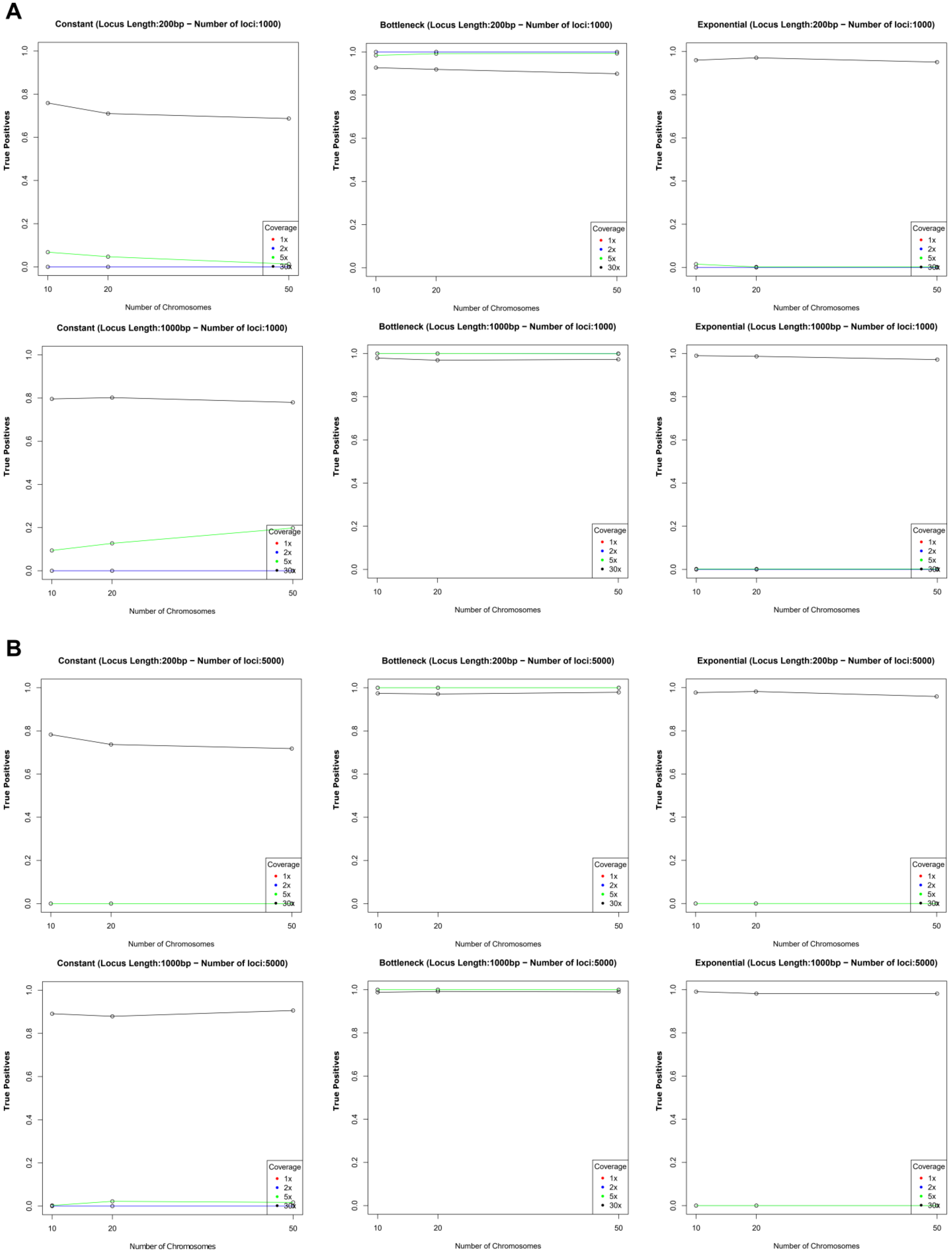

**Supplementary Figure 2. Proportion of True Positives for the two-population models, obtained correcting the pods through the genotype likelihood computed by ANGSD.** The plots have the same features of Figure 1.

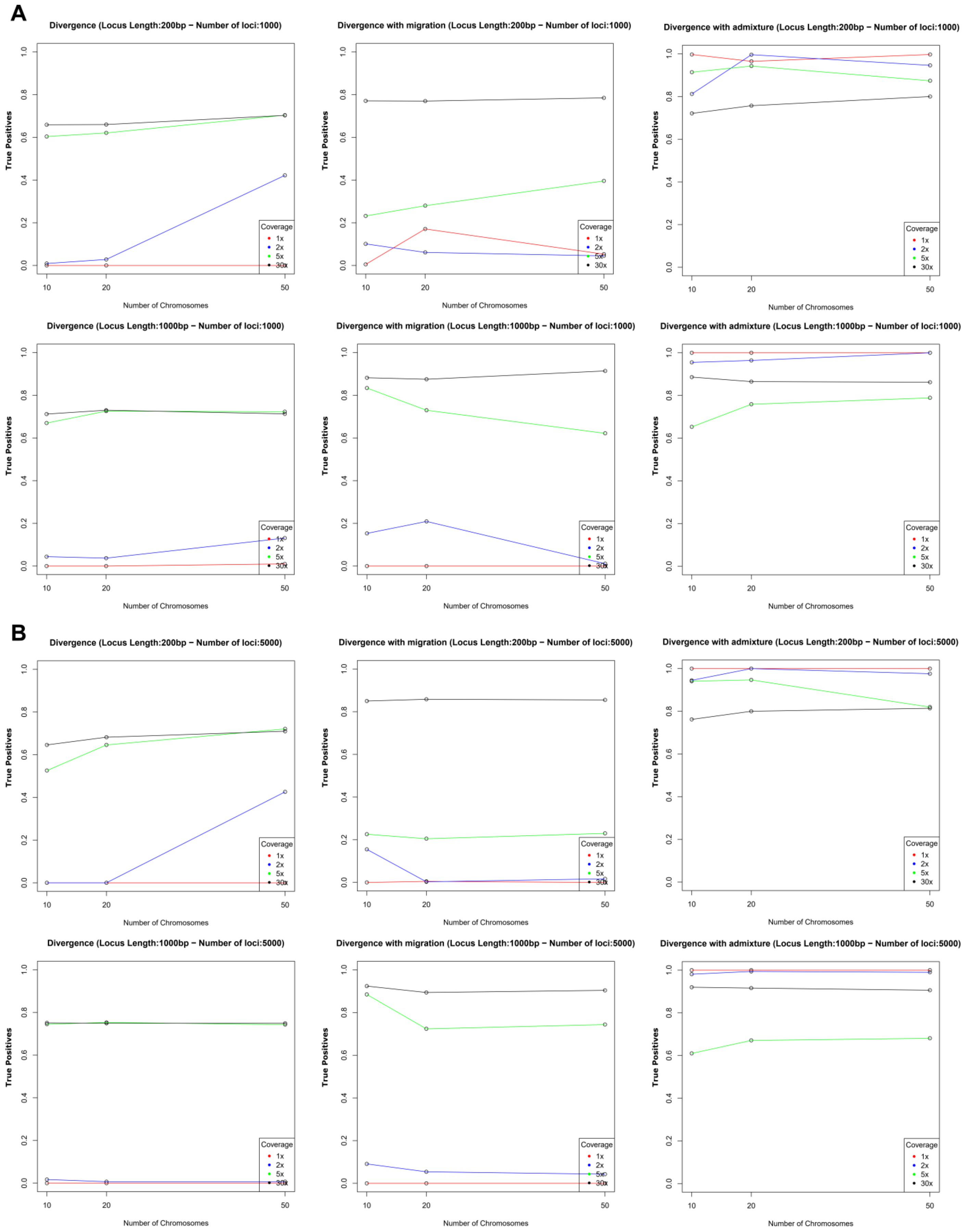
